# Targeted Protein Degradation of NUDT5 Reveals an Unexpected Non-Enzymatic Role in 6-Thioguanine-Mediated Toxicity

**DOI:** 10.1101/2025.03.16.643557

**Authors:** Anne-Sophie M. C. Marques, Ludwig G. Bauer, Tuan-Anh Nguyen, Alejandro Gonzalez Orta, Jan-Lennart Venne, Lai Cheng, Esra Balikci, Barr Tivon, Nir London, Stefan Kubicek, Kilian V. M. Huber

## Abstract

6-Thioguanine (6-TG), an FDA-approved antimetabolite drug, is widely used in the treatment of leukemia. Its cellular effects require metabolic activation and are regulated through interactions with various proteins such as NUDT15, which catalyzes the hydrolysis of the active 6-TG metabolites 6-thio-deoxyGTP (6-thio-dGTP) and 6-thio-GTP. Recent genome-wide CRISPR loss-of-function studies have identified another NUDIX hydrolase, NUDT5, as a crucial mediator of 6-TG toxicity. Here, we present the development and characterization of potent and selective NUDT5 degraders, guided by a cell-based assay screening strategy. These degraders, in conjunction with orthogonal CRISPR knock-out and reconstitution experiments, reveal a novel and unexpected, non-enzymatic role for NUDT5 in modulating the cellular response to 6-TG. Depletion of NUDT5 protein is antagonistic to NUDT15 inhibition, suggesting a distinct mode-of-action with potential implications for patient therapy.

## Introduction

NUDT5 is a conserved NUDIX hydrolase which cleaves the pyrophosphate bond within a range of nucleoside substrates such as ADP-ribose, ADP-glucose, and GDP-glucose^1^. Among the 22 human NUDIX proteins, NUDT5 plays a key role in ATP metabolism and chromatin remodelling, thereby conferring a potential vulnerability in hormone-dependent breast cancer^2,3^. Consistent with these data, NUDT5 knockdown suppresses migration and colony formation of T47D cells^1,3^. Recent reports suggest NUDT5 as a potential target in triple-negative breast cancer (TNBC) where its expression correlates with poor prognosis^4-6^. In addition to its role in cancer^7-9^, NUDT5 has been found to be an essential factor for modulating the toxicity of 6-thioguanine (6-TG)^10,11^. Introduced in the 1950s, 6-TG has been utilised as an antimetabolite drug to treat leukemia, autoimmune conditions, and inflammatory bowel disease (IBD) both in adults and children^12^. Mechanistically, 6-TG is thought to act as a prodrug, undergoing a cascade of metabolic conversions before being incorporated into DNA or RNA to exhibit its phenotypic effects. While NUDT5 inhibitors have been reported^1^, the impact of NUDT5 catalytic inhibition on 6-TG-mediated toxicity remains elusive. In this study, we describe the development of potent and selective NUDT5 PROTACs which are able to efficiently deplete NUDT5 protein from a range of cell types. In contrast to enzymatic inhibitors, NUDT5 PROTACs rescue cells from 6-TG-induced cell death, thus phenocopying previous CRISPR results. Our findings indicate a previously unidentified, non-enzymatic function for NUDT5 in purine metabolism that is distinct from the established role of NUDT15 as a genetic determinant and patient biomarker for 6-TG sensitivity.

## Results

In a previous CRISPR genome-wide loss-of-function screen conducted in HEK293T, HT29, and A375 cells, NUDT5 was identified as a crucial mediator of 6-TG toxicity^10^. Given the widespread use of 6-TG in leukemia treatment, we confirmed that knockout of NUDT5 confers resistance in HAP1 cells, a Philadelphia chromosome-positive, near-haploid human cell line derived from KBM-7 chronic myelogenous leukemia (CML) cells. We observed a four-fold increase in 6-TG IC_50_ values when comparing NUDT5 wild-type (WT) versus NUDT5 knockout (NUDT5 KO) cells (Fig. 1A). Next, we tested the published NUDT5 inhibitor TH5427 as well as a novel, chemically distinct NUDT5 chemical probe, MRK-952 (https://thesgc.org/chemical-probes/mrk-9520, to be published) for their ability to desensitize HAP1 cells to 6-TG treatment. Contrary to expectations, neither inhibitor was able to phenocopy the NUDT5 knockout in HAP1 or HL-60 cells (Fig. 1B). These results suggest that suppression of NUDT5 enzymatic activity alone is insufficient to rescue 6-TG-mediated toxicity. Intrigued by these findings, we decided to further dissect the role of NUDT5 in this context using a targeted protein degradation (TPD) strategy. We envisioned linking established NUDT5 ligands to E3 ligase moieties to generate proteolysis targeting chimeras (PROTACs), which would allow differentiation between enzymatic and potential non-enzymatic functions of NUDT5 (Fig. 1C).

**Figure 1:**
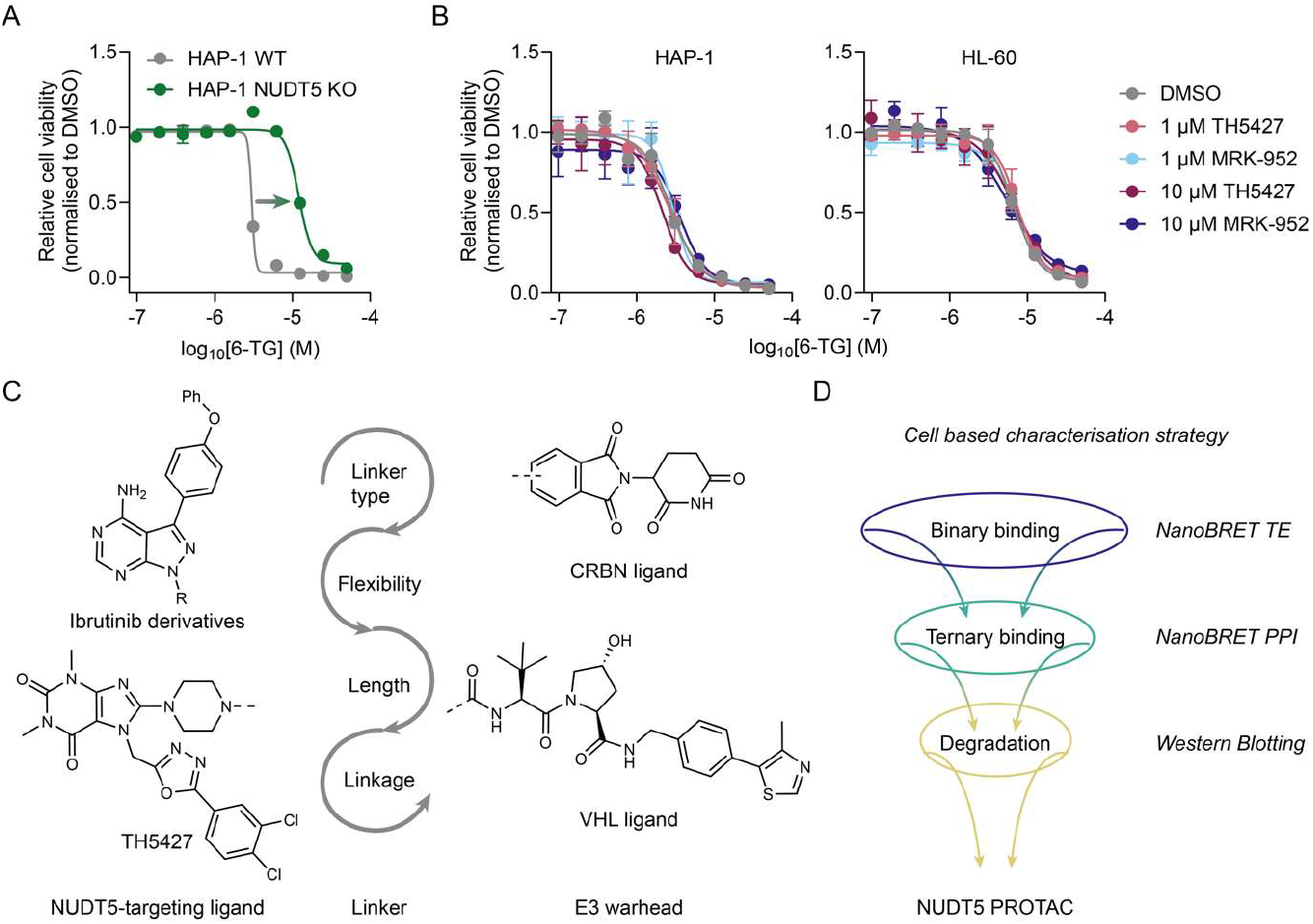
NUDT5 enzymatic activity is dispensable for 6-TG-mediated cancer cell cytotoxicity. A) NUDT5 knockout de-sensitizes HAP1 leukemia cells towards 6-TG. B) Inhibition of NUDT5 enzymatic activity fails to rescue 6-TG-mediated toxicity in HAP1 and HL-60 cells. C) NUDT5 PROTAC design strategy. D) Cell-based assay screening cascade for development of NUDT5 PROTACs.

We previously reported that ibrutinib, an FDA-approved BTK inhibitor, engages NUDT5 in cells^13^. This finding, in conjunction with the fact that multiple series of ibrutinib-based BTK PROTACs have been published^14,15^, prompted us to investigate if any such compounds might be able to also degrade NUDT5. Close inspection of co-crystal structures of ibrutinib bound to kinases compared to our recent NUDT5 complexes supported this hypothesis. We identified the N-3 and N-4 piperidine nitrogen in ibrutinib and compound 9^13^, respectively, as potential solvent exposed exit vectors for further functionalization (Supplementary Fig. S1). Next, we synthesized a library of VHL-based PROTACs featuring various PEG- and carbon-based linkers (Supplementary Information) and complemented this set with additional published ibrutinib based-degraders including the commercial MT-802^14^ and previously reported CRBN-targeting BTK degraders^15^. We assessed the performance of all PROTACs via a cell-based screening cascade surveying binary NUDT5 target engagement (TE) followed by ternary complex formation assays monitoring the interaction between NUDT5 and the respective E3 ligase. Active compounds were then further evaluated for NUDT5 degradation by Western blotting (Fig. 1D). However, although some ibrutinib-derived PROTACs were able to engage NUDT5 in cells, none of them appeared to induce formation of a productive E3-target ternary complex (Supplementary Table S1).

Next, we focused our synthetic strategy on the highly potent NUDT5 inhibitor TH5427 as a potential warhead for PROTACs^1^ (Fig. 1C). We obtained 15 bifunctional compounds with different linkers targeting either VHL or CRBN which we subsequently triaged in our cell-based screening cascade. While most compounds seemed to potently engage NUDT5, only CRBN-recruiting degraders efficiently promoted ternary complex formation after 4 h or 24 h (Fig. 2A and Supplementary Table S2). We assessed degradation for the most active compounds by Western Blotting across five concentrations (Fig. 2B and Supplementary Fig. S2). Overall, we observed that NUDT5 cellular engagement is not a primary determinant for degradation as many potent NUDT5 binders exhibited no significant effects on NUDT5 protein levels (e.g. ASM111, ASM122, ASM137). Alkyl linker-based PROTACs only induced moderate degradation independent of linker length, whereas for PEG-based compounds longer linkers were associated with higher D_max_ (DC_50_: ASM135 < ASM68 < ASM127 < ASM130) (Fig. 2B). For the most potent and efficient NUDT5 degraders from this series, ASM127 and ASM130, harboring three and four PEG units in the linker, respectively (Fig. 2C,D), we determined DC_50_ values of 12 nM and 7 nM, and apparent D_max_ values of 71% and 75%, respectively (Fig. 2E,F). No effects on NUDT5 protein levels were observed in HEK293 CRBN^KO^ cells indicating a CRBN-dependent degradation mechanism (Fig. 2G). Global proteomic analysis of HEK293 cells treated with either ASM127 or ASM130 revealed NUDT5 as the most significantly downregulated protein across 7,135 quantified proteins (Fig. 2H). The data also suggested a slight effect on the zinc finger protein ZFP91, a common off-target of PROTACs featuring an ortho-substituted phthalimide^16^. To rationalize the observed effects on NUDT5, we conducted ProsettaC^17^ analyses which provided plausible models, suggesting a similar binding mode for the active PEG-linker compounds (ASM68, ASM127, and ASM130) (Fig. 2D and Supplementary Fig. S3).

**Figure 2:**
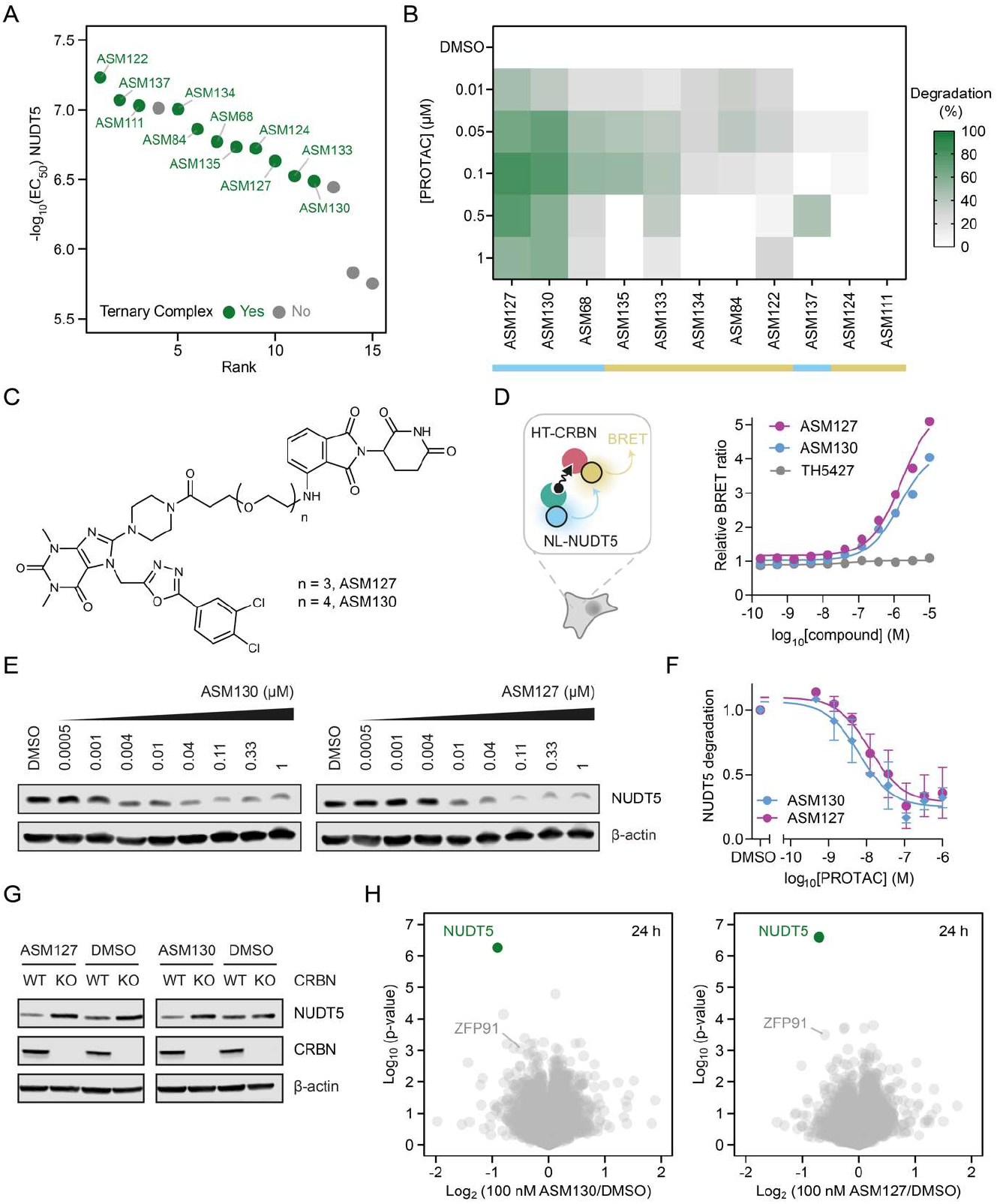
Characterization of the NUDT5 degraders ASM127 and ASM130. A) Multiparametric ranking of PROTACs based on binary NUDT5 target engagement and compound-induced NUDT5-E3 ternary complex formation in HEK293 cells. B) Heat map summary of PROTAC degradation efficiency across five concentrations after 20 h treatment in HEK293 cells. Yellow: PEG-based linker, Blue: carbon-based linker. Degradation was calculated from normalized Western blot intensities as the mean from two independent biological replicates (n = 2) (Supplementary Fig. S2). C) Chemical structures of ASM127 and ASM130. D) NanoBRET ternary complex formation assay results for ASM127, ASM130, and TH5427 using NL-NUDT5 and HT-CRBN after 4 h treatment in HEK293 cells. Data are shown as mean ± SD and based on three technical replicates. Graph is representative of two independent biological replicates (n = 2). E) Western blot showing dose-dependent degradation of NUDT5 in HEK293 cells after 20 h incubation with indicated compounds. Blot is representative of two independent biological replicates (n = 2). F) DC_50_ value determination for ASM127 and ASM130 (12 nM and 7 nM, respectively); estimated D_max_ 71% and 75%, respectively. Graphs show mean and SD of the normalized NUDT5 degradation efficiency derived from Western Blot band intensities (n = 2). G) CRBN knockout prevents PROTAC-mediated degradation of NUDT5 in HEK293 cells after 20 h of treatment with 100 nM ASM127 or ASM130. Western Blot is representative of two biological replicates (n = 2). H) Total proteome analysis results of HEK293 cells treated with 100 nM ASM127 or 100 nM ASM130 for 24 h (n = 4).

In an attempt to further improve degradation efficiency and potentially reduce the effect on ZFP91, we next turned our attention to more rigid linkers which have proven efficient in increasing therapeutic potency and metabolic stability^18,19^. The majority of new PROTACs from this series exhibited only limited effects with regard to NUDT5 degradation (Supplementary Table S3 and Supplementary Fig. S4). However, we identified one compound, hereafter referred to as dNUDT5 (Fig. 3A), featuring a tetrasubstituted urea linker motif, which induced strong and efficient downregulation of endogenous NUDT5 protein (Fig. 3B and Supplementary Fig. S5). To support further mechanistic studies, we synthesized a corresponding negative control, dNUDT5nc (Fig. 3A), by *N*-methylation of the imide ring to attenuate CRBN binding, thus preventing NUDT5 degradation (Fig. 3B and Supplementary Fig. S6A). Both dNUDT5 and dNUDT5nc engaged their cognate target NUDT5 in live cells as confirmed by NanoBRET assays (Supplementary Fig. S6B). As expected, only dNUDT5 induced proficient ternary complex formation between NUDT5 and CRBN in cells (Fig. 3C). We observed strong NUDT5 degradation in a concentration-dependent manner both in HAP1 cells (DC_50_ = 0.3 nM), as well as HEK293 cells (DC_50_ = 0.5 nM) (Fig. 3D and Supplementary Fig. S6C). Notably, concentrations above 300 nM appeared to result in a slight recovery of protein levels, likely due to the so-called hook effect commonly observed with PROTACs^20,21^. We conducted kinetic experiments indicating sustained degradation by dNUDT5 over a time course of 72 h (Fig. 3E and Supplementary Fig. S6D). Furthermore, we observed robust and consistent NUDT5 downregulation by dNUDT5 across a panel of cell lines such as A549, MCF7, and HAP1 representing different tissue backgrounds (Supplementary Fig. S7). We confirmed that degradation by dNUDT5 is UPS-dependent by performing a series of rescue experiments using neddylation, proteasome, and NUDT5 inhibitors as well as CRBN KO in HEK293 cells (Supplementary Fig. S6E). Global proteome analysis of HAP1 cells treated with dNUDT5 for 6 h and 24 h suggested exquisite selectivity with no other protein appearing significantly degraded in this context (Fig. 3F). To rationalize the markedly enhanced efficacy of dNUDT5 in comparison to previous compounds, we solved a co-crystal structure of dNUDT5 in complex with NUDT5, which indicated that dNUDT5 assumes a binding pose analogous to that of TH5427 (Figu. 3G).

**Figure 3:**
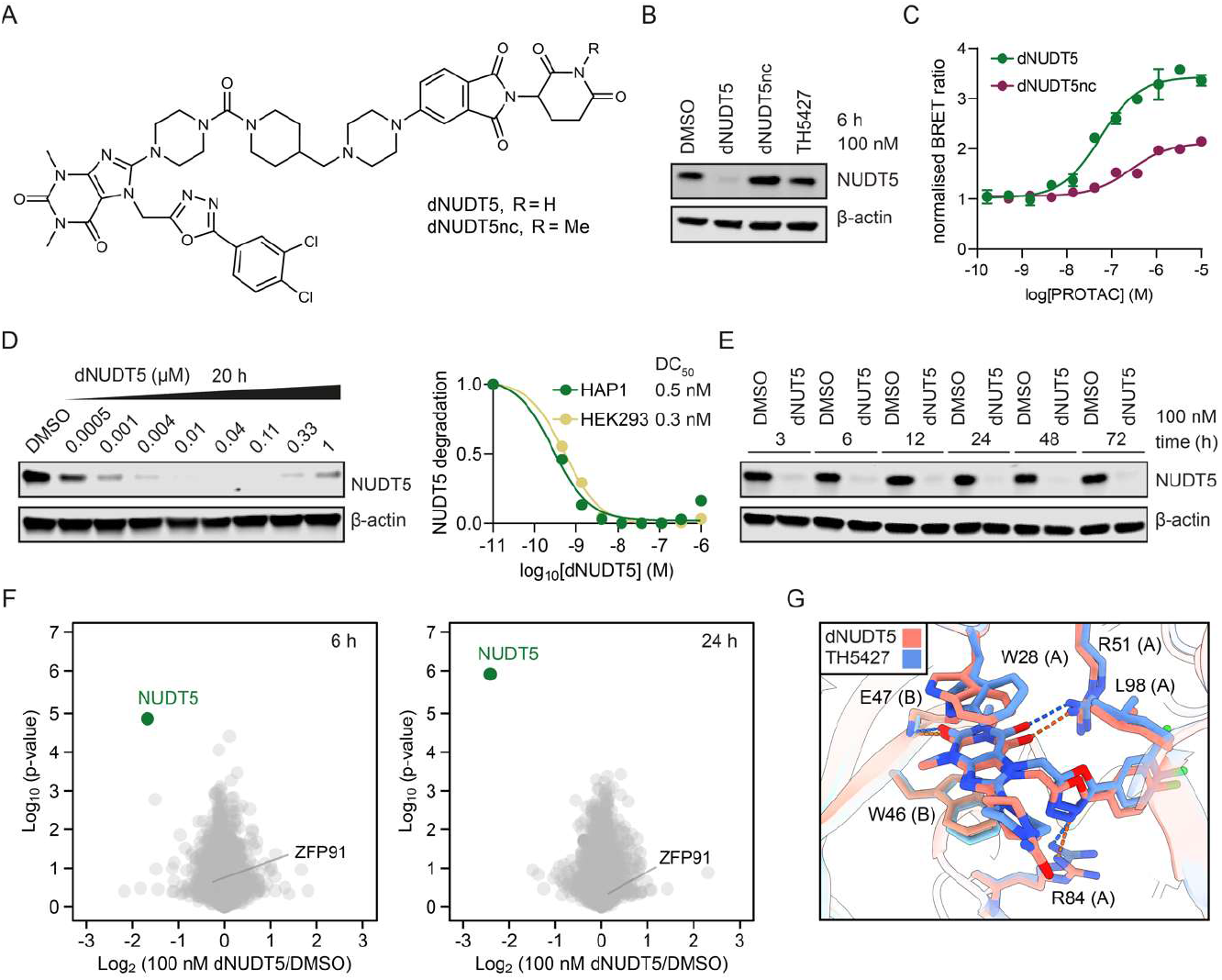
Characterization of dNUDT5. A) Chemical structures of dNUDT5 and dNUDT5nc. B) Western blot showing that dNUDT5, but not the methylated imide dNUDT5nc, is able to degrade NUDT5 in HAP1 cells. C) NanoBRET assay results demonstrating induced ternary complex formation by dNUDT5 in intact HEK293 cells. D) Western blot showing dose-dependent NUDT5 degradation by dNUDT5 in HAP1 and HEK293 cells. E) Western blot showing time-resolved NUDT5 degradation by dNUDT5 (100 nM) in HAP1 cells. F) Total proteome analysis of HAP1 cells treated with dNUDT5 for 6 h and 24 h, respectively (n = 3). G) Superimposition of dNUDT5 (pink, PDB: 9G6Y) and TH5427 (blue) NUDT5 co-crystal structures (PDB: 5NWH).

### Phenotypic evaluation of dNUDT5

With these potent and selective degraders in hand, we wondered whether our NUDT5 PROTACs would be able to rescue 6-TG-induced toxicity, analogous to CRISPR KO cells. We exposed HAP1 cells to escalating doses of our most potent compounds including dNUDT5, ASM127, and ASM130, in the presence of 2 µg/mL 6-TG for 72 h (Fig. 4A). Strikingly, the degraders exhibited a dose-dependent rescue of HAP1 cell viability, with maximal effects observed between concentrations of 50-100 nM, in accordance with their degradation efficiency (Supplementary Fig. S2, S5). Consistent with our previous results, dNUDT5 was the most efficient compound in abrogating 6-TG-induced cell death, whereas the inactive dNUDT5nc as well as the NUDT5 enzymatic inhibitor TH5427 failed to rescue cell viability. Notably, cell sensitivity towards 6-TG increased again at higher PROTAC concentrations almost perfectly aligning with the hook effect apparent in our Western blot experiments. Evaluation of an extended panel of cell lines confirmed that the rescue effect conferred by dNUDT5 is not cell-type specific since we observed similar results in HL-60, HCT116 and HEK293 cells (Fig. 4B). Importantly, treatment of naive HEK293 or HAP1 cells indicated that the degraders themselves did not affect cell viability *per se*, even when applied at high µM concentrations (Supplementary Fig. S8).

**Figure 4:**
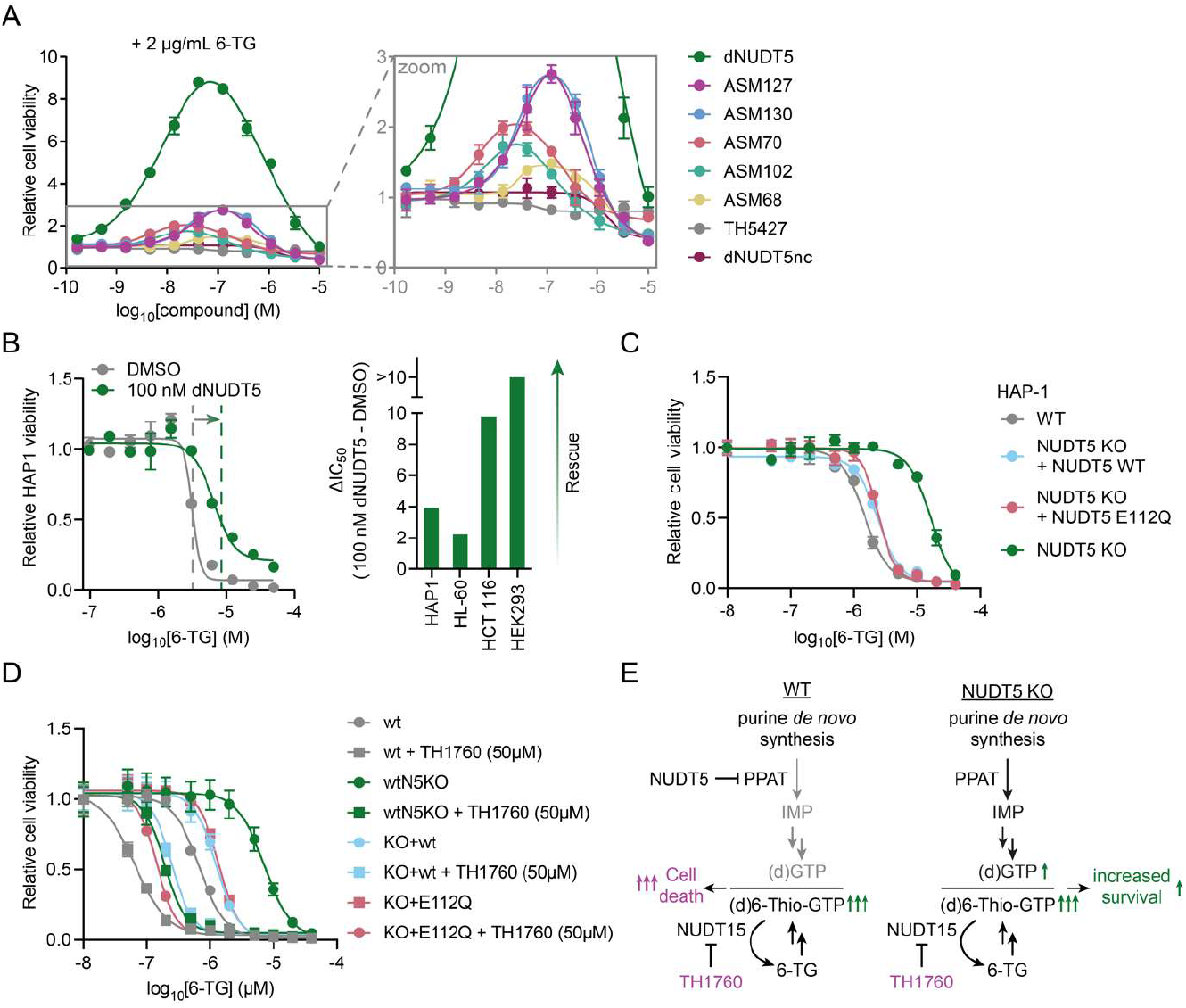
Depletion of NUDT5 protein by degraders or CRISPR KO rescues 6-TG-mediated toxicity and acts antagonistic to NUDT15 inhibition. A) Titration of indicated NUDT5 degraders and controls against a fixed concentration of 6-TG (2 µg/mL) in HAP1 cells for 72 h. B) dNUDT5 protects cells against 6-TG-induced toxicity across different cell lines. C) Reconstitution of HAP1 NUDT5 KO cells with either NUDT5 WT or the catalytically inactive E112Q mutant does not prevent 6-TG-dmediated cell death. Data are normalized to DMSO. Mean and standard deviation calculated from two independent biological replicates (n = 2). D) Effect of NUDT15 inhibition by TH1760 on 6-TG sensitivity across different cellular backgrounds. E) Proposed mechanism of NUDT5 and NUDT15 in 6-TG-mediated cell death.

These data strongly suggest that cell death mediated by 6-TG is dependent on a novel, non-enzymatic function of NUDT5. To corroborate this hypothesis, we reconstituted HAP1 NUDT5 KO cells with either wild-type (WT) or a catalytically inactive E112Q NUDT5 mutant and evaluated their sensitivity against 6-TG. Indeed, both the catalytically inactive mutant as well as the reconstituted WT exhibited comparable behaviour towards 6-TG treatment, similar to that observed in the parental cells (Fig. 4C).

### NUDT5 and NUDT15 act antagonistic with respect to 6-TG-induced toxicity

Interestingly, another member of the NUDIX family, NUDT15, has also been demonstrated to be a crucial factor influencing sensitivity towards 6-TG^22^. NUDT15 catalyses the conversion of 6-thio(d)GTP into 6-thio(d)GMP thereby preventing the incorporation of 6-TG metabolites into RNA and DNA. Importantly, recurring NUDT15 genetic variants observed in patients, which reduce NUDT15 catalytic activity, dramatically increase sensitivity towards 6-TG necessitating appropriate dose adjustments to prevent side effects such as leukopenia. Consequently, NUDT15 represents an established biomarker for thiopurine therapy. These observations have inspired the development of NUDT15 small molecule inhibitors such as TH1760^23^ to sensitise cancer cells towards 6-TG. Intrigued by these reports and our findings, we sought to investigate the relationship and potential mechanistic interplay between NUDT5 and NUDT15 with regard to 6-TG-mediated toxicity. We first titrated increasing concentrations of the NUDT15 inhibitor TH1760 against a fixed concentration of 6-TG (1 µM), which approximately corresponds to its cell viability IC_50_ value in HAP1 WT cells. Consistent with literature, we observed that inhibition of NUDT15 by TH1760 sensitised HAP1 WT cells towards 6-TG in a dose-dependent manner (Supplementary Fig. S9). Analogous to our previous results, NUDT5 KO attenuated 6-TG cytotoxicity, while reconstitution of NUDT5 KO cells with either NUDT5 WT or the catalytically inactive variant E112Q restored sensitivity (Fig. S9). Likewise, titration of 6-TG against a fixed concentration of TH1760 (50 µM) indicated cells treated with TH1760 were more sensitive toward 6-TG while this effect was partially rescued in NUDT5 KO cells (2.8-fold difference). Importantly, reconstitution of NUDT5 KO cells with the inactive NUDT5 E112Q mutant increased 6-TG-mediated cell death, further supporting the notion that NUDT5 enzymatic activity is dispensable in this context. Taken together, these results suggest that NUDT5 and NUDT15 act antagonistic with respect to 6-TG-mediated toxicity.

## Discussion

Targeted protein degradation (TPD) represents an intriguing alternative to classic small molecule inhibitors due to its distinct event-driven pharmacology. Whilst PROTACs are advancing in clinical trials^24^, well-characterized degraders also bear natural potential as chemical biology tools for drug target validation^25^ complementing genetic LOF screens^26,27^. Here, we apply this approach to the development and characterization of potent and highly selective NUDT5 degraders to uncover an unexpected non-enzymatic role of NUDT5 in 6-TG-mediated cell death, highlighting the power of TPD to dissect intricate domain-specific or scaffolding roles of human proteins in cellular signaling pathways^28,29^. While NUDT5 inhibitors have been tested extensively in various cancer models^1,3,5,6^, the exact mechanism by which NUDT5 KO confers resistance to 6-TG has not been elucidated^11,30^. Contrary to expectations, we find that inhibition of NUDT5 enzymatic activity fails to rescue cell death induced by 6-TG. Inspired by the success of TPD strategies in mimicking the effects of genetic LOF studies^31,32^, we developed a series of potent NUDT5 degraders to further elucidate its role in the cellular response to 6-TG. In a manner reminiscent of recent studies employing PROTACs to explore the non-catalytic functions of PARP1^33^ and Aurora kinase A^34^, we discover that only small-molecule-induced degradation, but not inhibition, of NUDT5 is able to recapitulate resistance to 6-TG analogous to CRISPR KO. Importantly, reconstitution of NUDT5 KO cells with either WT or a catalytically deficient NUDT5 mutant re-sensitizes cells towards 6-TG, confirming that NUDT5 enzymatic activity is not required to trigger cell death upon thiopurine treatment. Intriguingly, our results further establish NUDT5 as the second NUDIX family member influencing 6-TG pharmacology. Indeed, genetic mutations or catalytic inhibition of NUDT15 have been shown to enhance sensitivity towards 6-TG^35-40^. Notably, our data suggest NUDT5 and NUDT15 exert discrete and conceptually antagonistic effects toward 6-TG. While NUDT5 depletion results in 6-TG resistance, suppression of NUDT15 enzymatic activity sensitizes cells towards 6-TG treatment leading to increased cell death. In conjunction with results from another study (cf. complementary manuscript by Lin, Nguyen, Marques et al.), we hypothesise that cell sensitivity towards 6-TG is determined by the ratio between (d)GTP and 6-thio(d)GTP regulated by NUDT5 and NUD15, respectively (Fig. 4E). Systematic profiling of NUDT5 protein-protein interactions suggests an obligate interaction with PPAT, the enzyme that catalyses the first rate-limiting step in the purine biosynthesis pathway^41^. Under NUDT5 WT conditions, NUDT5 binds to PPAT and represses the formation of (d)GTP, allowing more 6-thio(d)GTP to be incorporated into RNA and DNA. In NUDT5 KO cells, increased purine *de novo* synthesis leads to elevated levels of (d)GTP effectively outcompeting toxic 6-TG metabolites. In summary, our collective findings suggest that NUDT5 fulfills an unprecedented fundamental, non-catalytic role in regulating purine biosynthesis, which in turn affects 6-TG antiproliferative activity with important considerations for thiopurine therapy in cancer and beyond.

## Acknowledgments

The authors thank the CeMM Proteomics Platform and Discovery Proteomics Facility, Oxford University, for their support. HEK293 CRBN KO cells were a kind gift from J. Qi, Dana-Farber Cancer Institute. This project received funding from the European Research Council (ERC) under the European Union’s Horizon 2020 research and innovation programme (SK, ERC-CoG-772437), a Marie Sklodowska-Curie Postdoctoral Fellowship (TAN, grant number: 101106260), and the Innovative Medicines Initiative 2 Joint Undertaking (JU) under grant agreement No 875510. The JU receives support from the European Union’s Horizon 2020 research and innovation program, the European Federation of Pharmaceutical Industries and Associations (EFPIA), the Ontario Institute for Cancer Research, the Royal Institution for the Advancement of Learning McGill University, the KTH Royal Institute of Technology (Kungliga Tekniska Hoegskolan) and Diamond Light Source Limited.

## The authors declare the following competing interests

International Patent Application No. WO2024127022A1, filed in the name of Oxford University Innovation Limited.

## Notes

### Competing Interest Statement

The authors have declared no competing interest.

