## Supplementary Information for "Targeted Protein Degradation of NUDT5 Reveals an Unexpected Non-Enzymatic Role in 6-Thioguanine-Mediated Toxicity"

**Table of Contents**

Figure S1. Design strategy for the synthesis of ibrutinib-derived PROTACs……………......………S3
Table S1. NanoBRET assay results for ibrutinib-derived PROTACs……………………..………….S4
Table S2. NanoBRET assay results for flexible linker-based PROTACs……………………….….…...S6
Figure S2. Western blot evaluation of NUDT5 degradation by flexible linker-based PROTAC in HEK293 cells…………………………………………………………………………….………S8
Figure S3. PRosettaC ternary complex prediction for active PROTACs.……………………….……S9
Table S3. NanoBRET assay results for rigid linker-based PROTACs.…………………….……S10
Figure S4. Western blot evaluation of NUDT5 degradation by rigid linker-based PROTACs in HEK293 cells……….………………………………………………………………………..S11
Figure S5. Western blot evaluation of NUDT5 degradation by dNUDT5 in HEK293 cells………...S11
Figure S6. Characterisation of dNUDT5 in HEK293 cells.…………………………………………S12
Figure S7. NUDT5 degradation efficiency of dNUDT5, ASM130, ASM127, ASM68, ASM102 and ASM137 assessed by Western Blotting in A549, HAP1, and MCF7 cells.…… ..…………………S13
Figure S8. NUDT5 PROTACs do not affect cell viability of HEK293 and HAP1 cells…………...S13
Figure S9: Effect of TH1760 in HAP1 WT, NUDT5 KO or reconstituted cells.……………………S14
Table S4. Data collection and refinement statistics for the co-crystal structure of dNUDT5 bound to NUDT5……………………..………………………………………………………...……S15
Methods.……………………………………………………………………………………..S16
Synthesis and compound characterization.…………………………………………………..S19
References……………………………………………………………………………...…….S61

**Supplementary Figures**

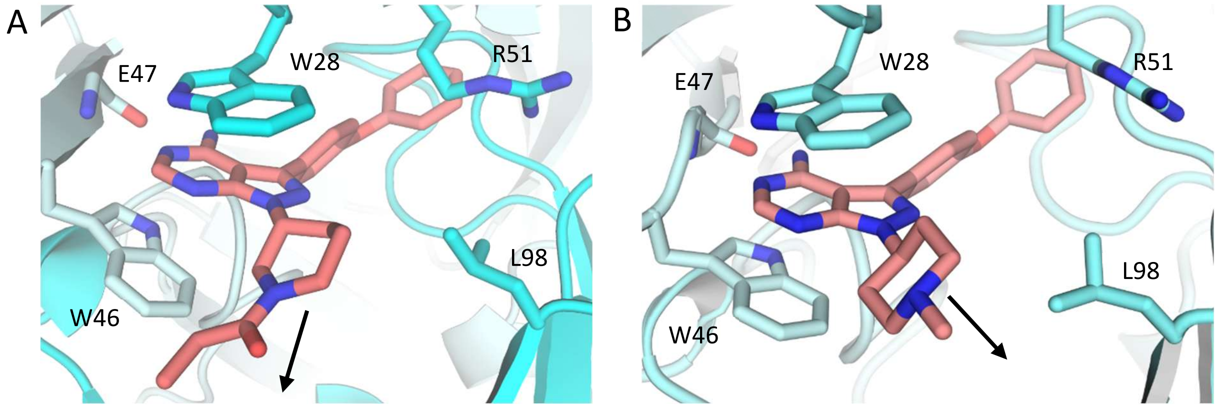

**Figure S1. Design strategy for the synthesis of ibrutinib-derived PROTACs**

(A) Co-crystal structure of NUDT5 bound to ibrutinib (PDB: 8RDZ). The arrow depicts the exit vector. (B) Co-crystal structure of NUDT5 bound to the ibrutinib analogue (compound 9) (PDB: 8RIY). The arrow depicts the exit vector.

**Table S1. NanoBRET assay results for ibrutinib-derived PROTACs.**

NUDT5 target engagement (TE) and PROTAC ternary complex formation measurements were conducted in HEK293 cells at the indicated time points. Results represent the mean of 3 technical replicates.

| Compound | Ligand | Linker | E3 ligase ligand | $EC_{50}$ (µM), TE 4h | Ternary complex formation below 1 µM (20 h) |
|---|---|---|---|---|---|
| **ASM32** | 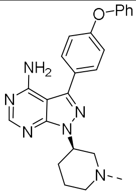 |  | 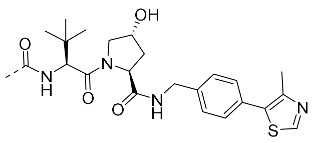 | >10 µM | No |
| **ASM72** | 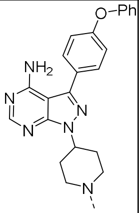 | 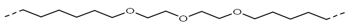 | 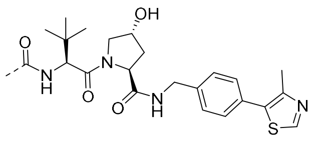 | 6 | No |
| **ASM73** | 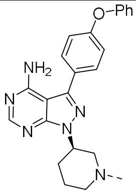 |  | 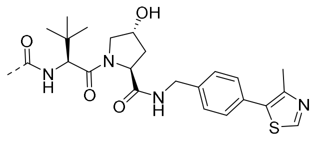 | >10 µM | No |
| **ASM74** | 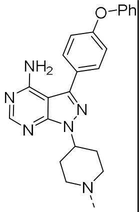 | 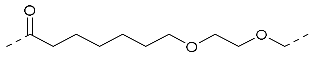 | 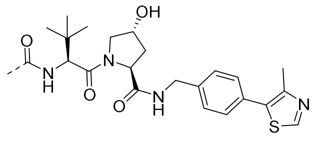 | >10 µM | No |
| **ASM75** | 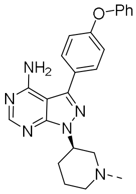 | 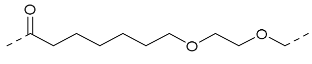 | 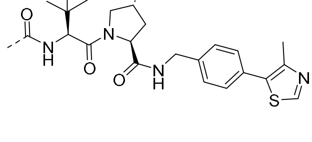 | >10 µM | No |

| | | | | | |
|---|---|---|---|---|---|
| **MT-802** | 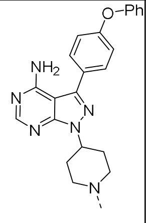 | 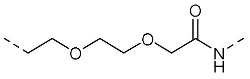 | 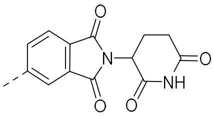 | 1.3 | No |
| **RG27** | 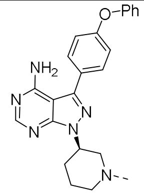 | 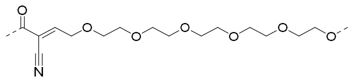 | 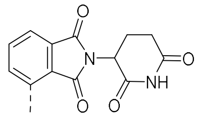 | >10 μM | No |
| **RG38** | 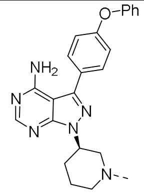 | 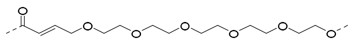 | 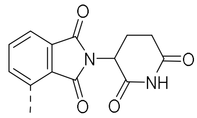 | >10 μM | No |
| **RG52** | 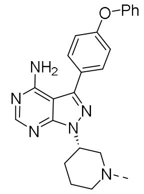 | 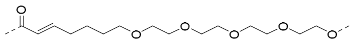 | 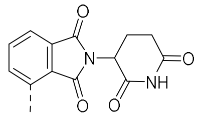 | >10 μM | No |
| **RG55** | 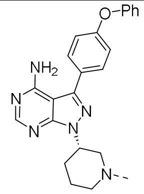 | 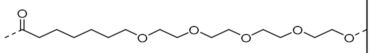 | 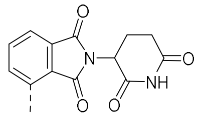 | >10 μM | No |
| **RG66** | 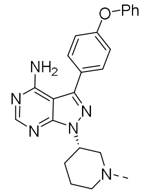 | 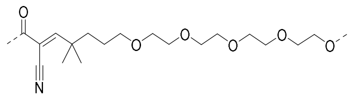 | 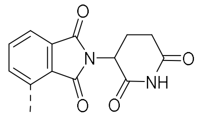 | >10 μM | No |
| **P13I[1]** | 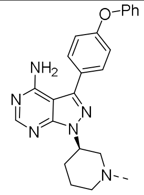 | 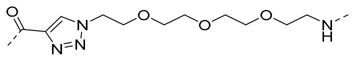 | 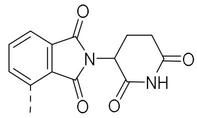 | 8.9 | No |

**Table S2. NanoBRET assay results for flexible linker-based PROTACs.**

NUDT5 TE and PROTAC ternary complex formation measurements were conducted in HEK293 cells at the indicated time points. Results represent the mean of 3 technical replicates.

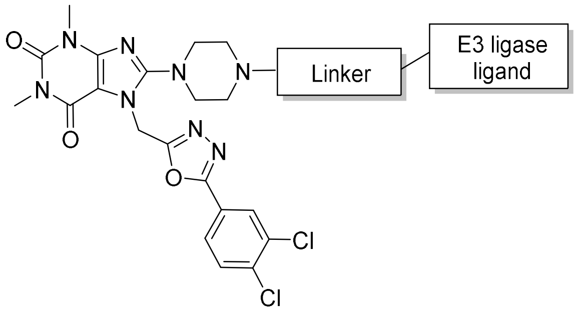

| Compound | Linker | E3 ligase ligand | $EC_{50}$ (nM) TE (4h) | Ternary complex formation below 1 µM (4 h) | Ternary complex formation below 1 µM (20 h) |
|---|---|---|---|---|---|
| **ASM076** | 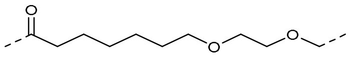 | 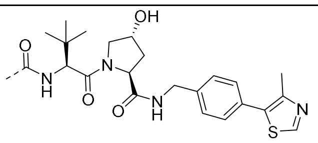 | 2000 | NT[a] | No |
| **ASM131** | 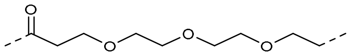 | 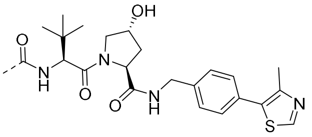 | 360 | No | No |
| **ASM137** | 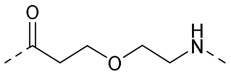 | 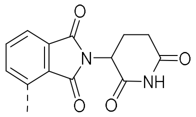 | 85 | Yes | Yes |
| **ASM68** | 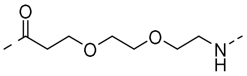 | 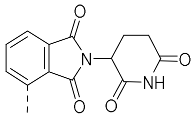 | 170 | NT | Yes |
| **ASM127** | 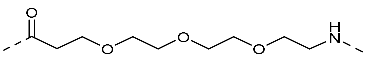 | 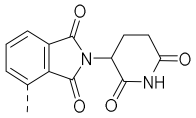 | 298 | Yes | Yes |
| **ASM130** | 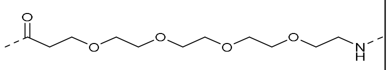 | 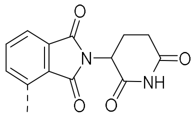 | 326 | Yes | Yes |
| **ASM111** | 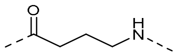 | 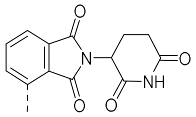 | 93 | NT | Yes |

| | | | | | |
|---|---|---|---|---|---|
| **ASM124** | 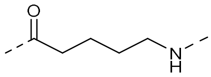 | 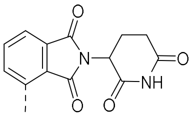 | 190 | Yes | Yes |
| **ASM122** | 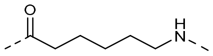 | 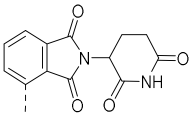 | 59 | Yes | Yes |
| **ASM84** | 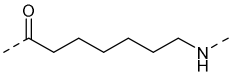 | 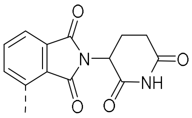 | 138 | NT | Yes |
| **ASM136** | 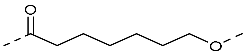 | 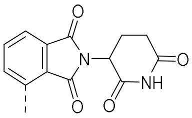 | 97 | No | No |
| **ASM134** | 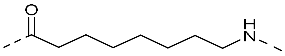 | 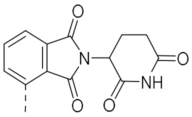 | 99 | Yes | Yes |
| **ASM57** | 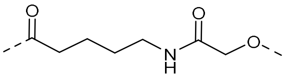 | 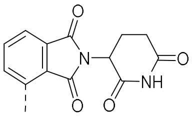 | 1770 | NT | No |
| **ASM133** | 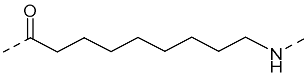 | 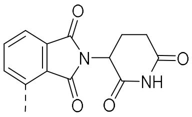 | 233 | Yes | Yes |
| **ASM135** | 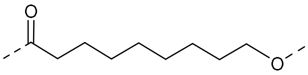 | 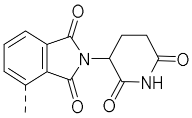 | 185 | Yes | Yes |

[a]NT, not tested.

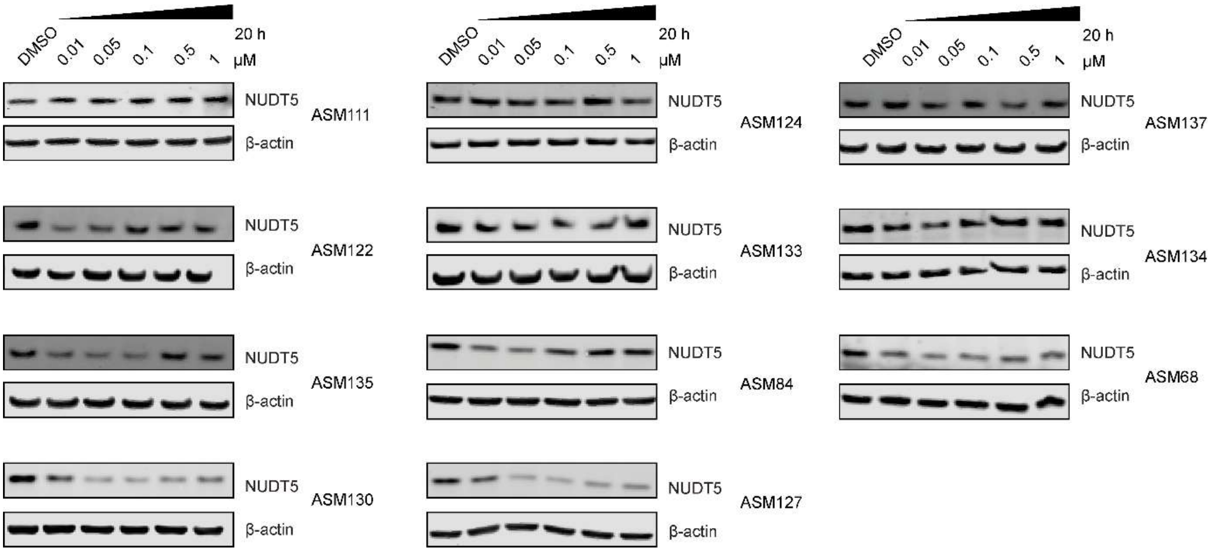

**Figure S2. Western blot evaluation of NUDT5 degradation by flexible linker-based PROTACs in HEK293 cells.** Western blots are representative of two independent biological replicates (n = 2).

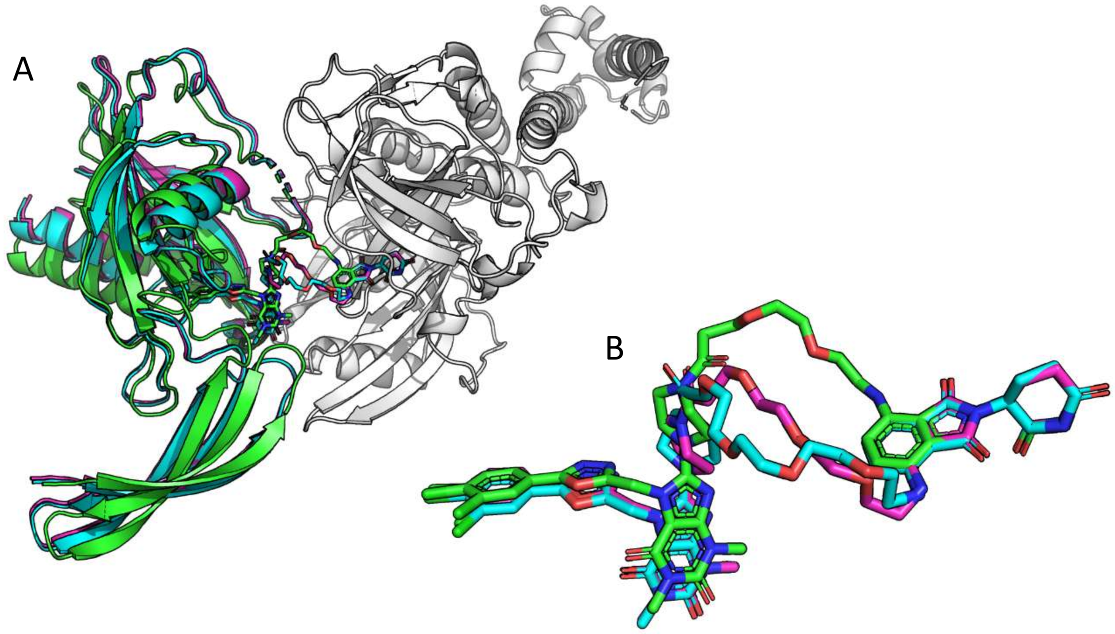

**Figure S3. PRosettaC ternary complex prediction for active PROTACs**. (A) Reclustering of the top-scoring models for ASM68 (green), ASM127 (magenta) and ASM130 (cyan) with an RMSD threshold of 2 Å. The top cluster includes 53 models (14 of ASM68, 24 of ASM127 and 15 of ASM130). The next clusters contain less than 20 structures and are typically limited to models arising from only one PROTAC, further increasing the confidence in this ternary complex prediction. (B) Zoom in on the PROTAC binding mode shows that all three PROTACs are able to bind similarly in this ternary complex. We should note that while ASM68 is predicted to have a flipped conformation of the phthalimide moiety, other members of the same cluster showed a similar conformation to those of ASM127 and ASM130, indicating both exits may be tolerated.

**Table S3. NanoBRET assay results for rigid linker-based PROTACs.**

NUDT5 TE and PROTAC ternary complex formation measurements were conducted in HEK293 cells at the indicated time points. Results are the mean of 3 technical replicates.

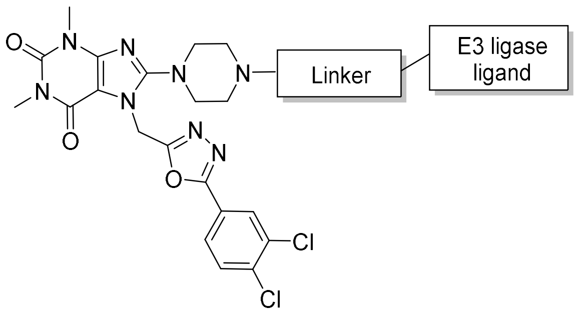

| Compound | Linker | E3 ligase ligand | $EC_{50}$ (nM)<br>TE (4h) | Ternary complex formation below 1 µM (20 h) |
|---|---|---|---|---|
| **ASM105** | 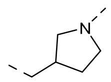 | 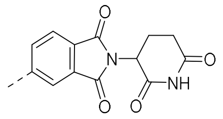 | 103 | No |
| **ASM110** | 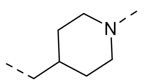 | 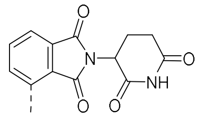 | 61 | No |
| **ASM70** | 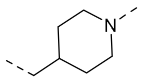 | 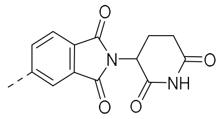 | 42 | Yes |
| **ASM71** | 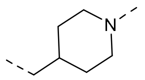 | 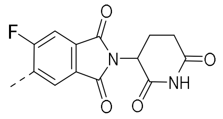 | 17 | No |
| **ASM102** | 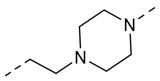 | 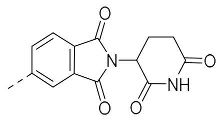 | 102 | Yes |

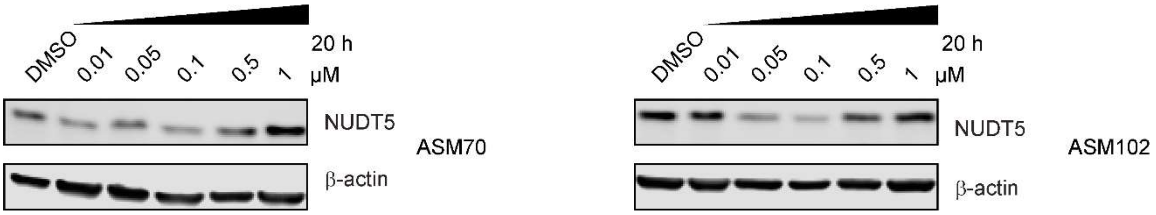

**Figure S4. Western blot evaluation of NUDT5 degradation by rigid linker-based PROTACs in HEK293 cells.** Western blots are representative of two independent biological replicates (n = 2).

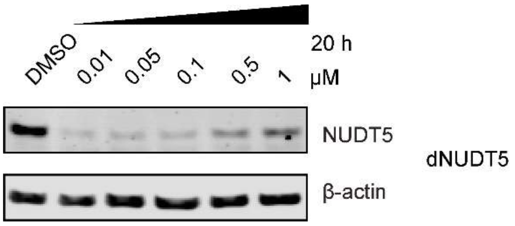

**Figure S5. Western blot evaluation of NUDT5 degradation by dNUDT5 in HEK293 cells.** Western blot is representative of two independent biological replicates (n = 2).

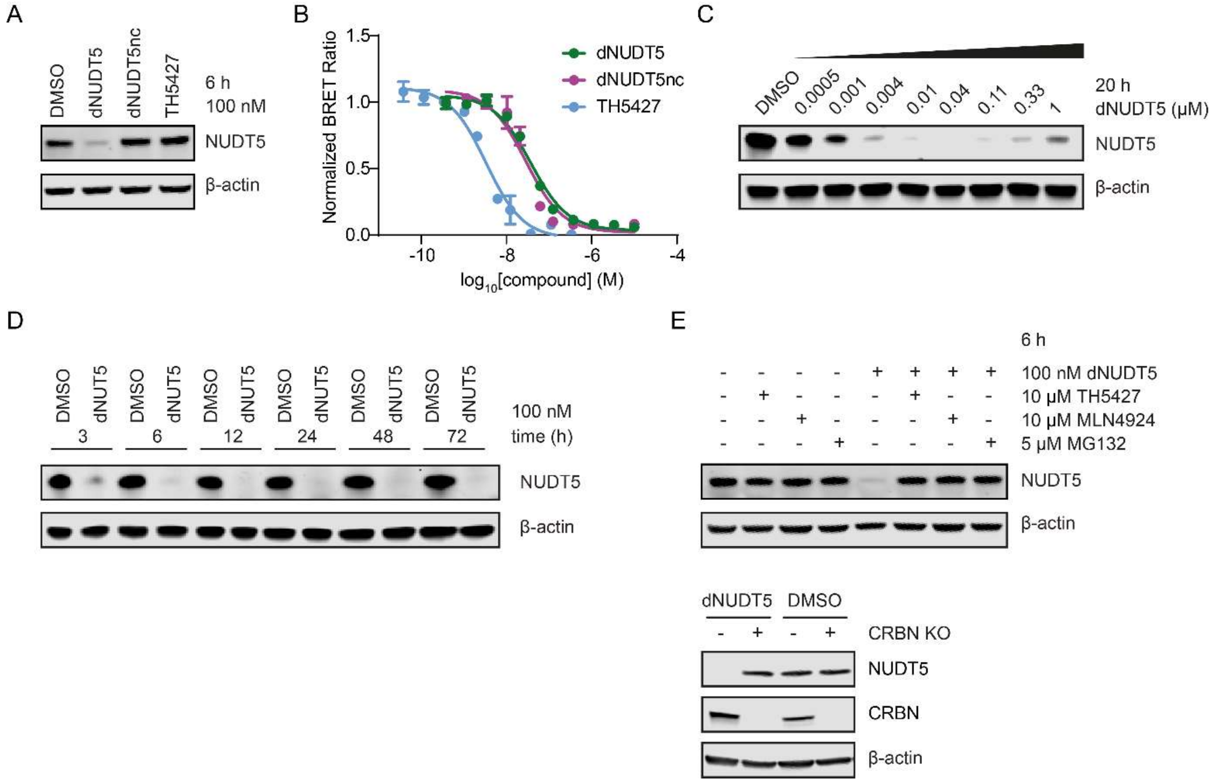

**Figure S6. Characterisation of dNUDT5 in HEK293 cells.**

A) Only dNUDT5, but not the CRBN-binding-deficient dNUDT5nc or the NUDT5 catalytic inhibitor TH5427, is able to degrade NUDT5 in HEK293 cells. B) NUDT5 NanoBRET TE assay results for dNUDT5 ($EC_{50}$ = 37 nM) and dNUDT5nc ($EC_{50}$ = 28 nM) compared to the positive control TH5427 ($EC_{50}$ = 3 nM). Data are shown as mean ± SD and are based on three technical replicates. Graphs are representative of two independent biological replicates (n = 2). C) Dose-dependent degradation of dNUDT5 in HEK293 cells. D) Time-resolved degradation of dNUDT5 in HEK293 cells at 100 nM. E) Rescue experiments in HEK293 cells with TH5427 (10 µM), the neddylation inhibitor MLN4924 (10 µM), the proteasome inhibitor MG132 (5 µM) and in $CRBN^{KO}$ HEK293 cells.

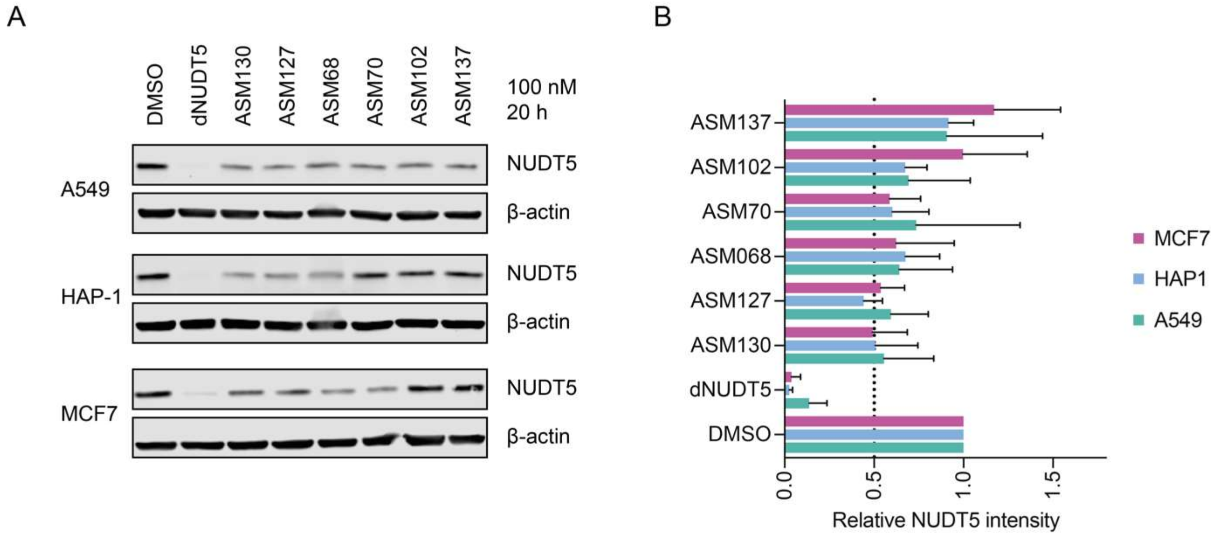

**Figure S7. NUDT5 degradation efficiency of dNUDT5, ASM130, ASM127, ASM68, ASM102 and ASM137 assessed by Western Blotting in A549, HAP1, and MCF7 cells.** A) Representative Western blot. B) Bar graph of mean and standard deviation of relative NUDT5 band intensities upon PROTAC treatment, normalised to DMSO (n = 4).

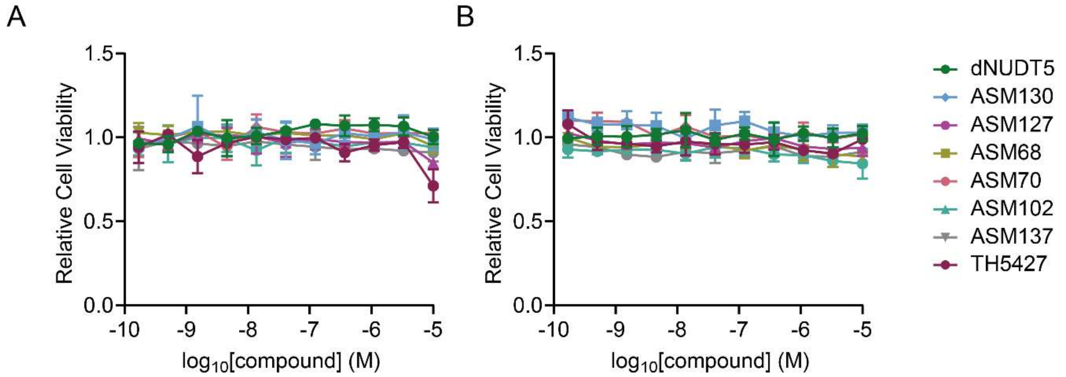

**Figure S8. NUDT5 PROTACs do not affect cell viability of HEK293 and HAP1 cells**.
Cell viability was measured by CellTiter Glo assay in (A) HEK293 and (B) HAP1 cells following PROTAC treatment for 72 h.

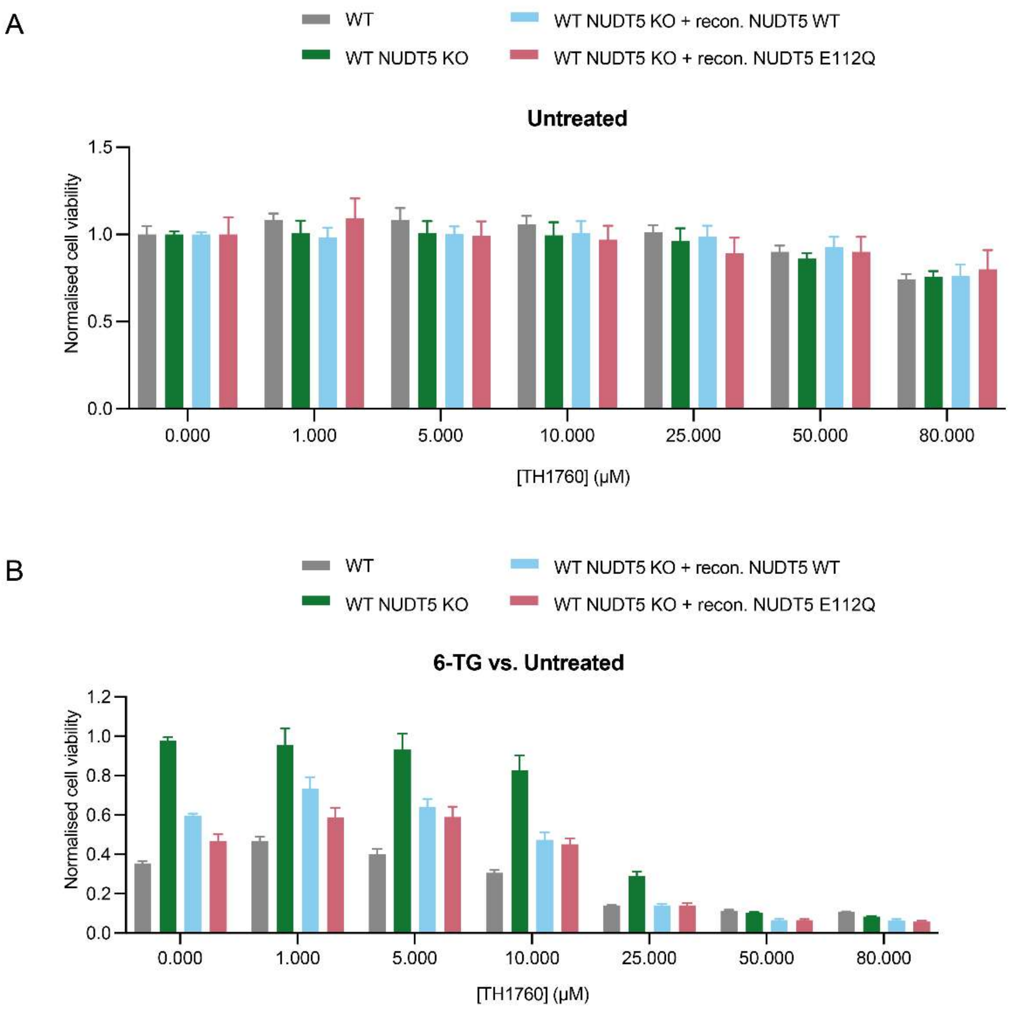

**Figure S9. Treatment with the NUDT15 inhibitor TH1760 increases cell sensitivity toward 6-TG, which can be reversed by NUDT5 depletion.** (A) Inhibition of NUDT15 at concentrations up to 50 µM does not affect HAP1 cell viability. Indicated cell lines were incubated with respective concentrations of TH1760 for 72 h. (B) Knockout of NUDT5 desensitizes cells towards 6-TG when NUDT15 is inhibited. The catalytic activity of NUDT5 (cf. NUDT5 KO vs. recon. NUDT5 WT and NUDT5 E112Q) is dispensable for the rescue mechanism. Indicated cell lines were treated with indicated amounts of TH1760 and in presence or absence of 6-TG (1 µM) for 72 h. Values were normalized to respective TH1760 concentration points.

**Table S4. Data collection and refinement statistics for the co-crystal structure of dNUDT5 bound to NUDT5.**

| | dNUDT5<br>PDB ID: 9G6Y |
|---|---|
| **Data collection** | |
| Space group | C 1 2 1 |
| Cell dimensions | |
| $a$, $b$, $c$ (Å) | 113.57, 40.21, 99.96 |
| α, β, γ (°) | 90.00, 121.92, 90.00 |
| Resolution (Å) | 37.11 - 1.90 (1.97 - 1.90) |
| $R_{sym}$ or $R_{merge}$ | 0.1344 (2.550) |
| $I / \sigma I$ | 7.7 (0.3) |
| Completeness (%) | 98.17% (83.55%) |
| Redundancy | 6.8 (6.7) |
| **Refinement** | |
| Resolution (Å) | 37.11 - 1.90 (1.97 - 1.90) |
| No. reflections | 29973 (2514) |
| $R_{work}$ / $R_{free}$ | 0.2124 (0.4314) / 0.2422 (0.4318) |
| No. atoms | 3278 |
| Protein | 3004 |
| Ligand/ion | 73 |
| Water | 201 |
| $B$-factors (Average) | 46.11 |
| Protein (mean) | 45.62 |
| Ligand/ion | 64.36 |
| Water | 46.80 |
| R.m.s. deviations | |
| Bond lengths (Å) | 0.003 |
| Bond angles (°) | 0.55 |
| Ramachandran Plot | |
| Favored (%) | 97.11 |
| Outliers (%) | 0.00 |

*Values in parentheses are for highest-resolution shell.

### Methods

#### Cell culture

HEK293, MCF7, A549 were cultured in DMEM + 10% FBS, HAP1 and HL-60 were cultured in IMDM + 10% FBS at 37 °C and 5% $CO_2$.

#### Western Blotting

Cells were harvested using TrypLE (Gibco, 12605010), washed 1x with PBS and frozen at –80 °C. Pellets were lysed on ice using RIPA buffer (25 mM Tris (pH 7.5), 150 mM NaCl, 1% Ipegal, 1% SDC, 0.1% SDC, 1x protease inhibitor (Roche, cOmplete mini, 11836170001), 1x phosphotase (Thermo Scientific, Halt, 78429), 0.1% benzonase (Merck, E1014-5KU)), incubated for 15 min and then centrifuged for 20 min at 4 °C at maximum speed in a table-top centrifuge. Protein concentration was determined by a DC assay (Bio-Rad, 5000112), adjusted, mixed with 1x Laemmli buffer (Bio-Rad, 1610747) and boiled for 5 min at 95 °C. Samples were resolved by SDS-PAGE (Bio-Rad, 3450125) and transferred to a nitrocellulose membrane (Cytiva, 10600014) via wet-transfer. Membranes were blocked with 6% milk in PBS-T for 1 h at RT, washed and then incubated with primary antibody diluted in PBS-T supplemented with 0.1% $NaN_3$ over night at 4 °C. Membranes were incubated with fluorophore-conjugated secondary antibodies, Alexa Fluor 680 anti-rabbit IgG (Life Technologies, A21109), Alexa Fluor 750 anti-mouse IgG (Life Technologies, A21037) diluted 1:5000 in 2 % milk PBS-T, washed and imaged with an Odyssey imager (Licor). Primary antibodies: 1:1000 NUDT5 (Abcam, ab129172), 1:200 β-Actin (Santa Cruz, sc-47778).

#### NanoBRET TE and ternary complex assays

NanoBRET target engagement assays were performed by transfection of NL-NUDT5 and 2.5 nM CBH-004 as previously described[2] (cite: Ibrutinib paper). NanoBRET ternary complex assays were performed following the manufacturer's instruction (Promega). In brief, HEK293 cells were transfected with NL-NUDT5 and either HL-VHL or HL-CRBN in a ratio of 1:100 and incubated overnight. Cells were adjusted to 200,000 cells/mL in Optimem + 4 % FBS, 100 nM HaloTag NanoBRET 618 Ligand spiked in and cells were seeded out in a 384 white well plate containing compounds that were pre-dispensed with an ECHO Liquid Handler. After 4 h or 20 h of incubation, a mix of substrate in assay medium was added into each well. Donor and acceptor signals were measured in a PHERAstar FSX or FS plate reader and analyzed by GraphPad Prism (v.9 or v.10). The acceptor/donor ratio was calculated, the no ligand background signal was subtracted and multiplied by 1000 to yield mBRET units. Data were then normalized to DMSO.

**Untargeted global proteomics**

*Sample preparation and acquisition*

Proteomics samples were digested with trypsin using STrap columns following the manufacturer's protocol (C02-micro, ProtiFi). For untargeted global proteomics HAP1 cells were treated with DMSO or 100 nM dNUDT5 for 6 h or 24 h. Then, the cell pellets were collected and lysed in 5 % SDS containing 1 µL/mL benzonase (E1014, Millipore) and subsequent short sonication to shear genomic DNA (1x 2 s pulse, 20 % amplitude, microtip, Sonics Vibra Cell). 50 µg protein from the total cell lysate or 80 % of the elution fracion of the chemical pulldown were first reduced by 20 mM dithiothreitol (M02712, Fluorochem), and alkylated by 40 mM iodoacetamide (I1149, Sigma). Phosphoric acid and S-Trap protein binding buffer were added into the sample lysates. Then, the SDS lysate/S-Trap buffer were loaded into a S-Trap column (C02-micro-80, ProtiFi). 1 µg of trypsin (V5111, Promega) was used to digest each sample at 37 °C for 20 h. The samples were then eluted and dried by vacuum centrifugation. Peptide pellets were resuspended in mass spectrometry grade water with 2 % acetonitrile (85188, Thermo Scientific) and 0.1 % trifluoroacetic acid (85183, Thermo Scientific). The samples were injected into Orbitrap Fusion Lumos Tribrid Mass Spectrometer (Thermo Scientific). Sample acquisition was performed as previously described[3].

*Data analysis*

Raw data were searched against the human database (UP000005640, downloaded 08/21) using DIA-NN (v.1.8.1)[4] by enabling FASTA digest for library free search and Deep learning-based spectra, RTs and IMs prediction. Trypsin/P was selected for the in silico digestion of the provided fasta file, allowing up to 2 missed cleavages. Fixed modifications included N-terminal methionine excision and cysteine carbamidomethylation. Variable modifications were methionine oxidation and N-terminal acetylation, with a maximum of one modification per precursor. Precursor FDR was kept at 1% and MBR enabled. The protein groups matrix was further analysed using Perseus (2.0.9.0)[5] or R studio (4.3.2). Potential contaminants were filtered against the top 300 most common Homo sapiens contaminants reported in CRAPome[6]. Proteins were removed if detected less than 70% across all runs. After column-wise normalisation by median subtraction, missing values were imputed by random numbers from a normal distribution (1.8 standard deviation, downshift width 0.3). Volcano plots were generated by calculating fold change and -log10(p-value) from a two-sided Student's t-test.

**Cell viability and 6-TG assays**

Cells were counted and adjusted to a density of 25,000 cells/mL in culture medium. A volume of 40 µL of the cell suspension was dispensed into white polypropylene 384-well plates (781207, Greiner) pre-dispensed with test compounds using an Echo Liquid Handler. Plates were sealed with breathable film and incubated at 37 °C with 5% $CO_2$ for 72 hours. Cell viability was assessed using the CellTiter-Glo Luminescent Cell Viability Assay, following the manufacturer's protocol (Promega). Luminescence was measured using a PHERAstar FS or FSX plate reader, and data were analyzed using GraphPad Prism software (versions 9 or 10). Results were normalized to the vehicle control (DMSO). To evaluate cell viability in the presence of 6-TG, the compound was spiked into the cell suspension at a concentration of 2 µg/mL or as otherwise specified, and the experiment was conducted as described above.

**TH1760/6-thioguanine assay:** HAP1 cell lines were cultured in IMDM full media in 96 well plates (1,200 cells/100 µl/well) containing indicated concentration series of TH1760/6-TG and where specifically stated additionally with corresponding concentration of 6-TG (1 µM) or TH1760 (50 µM), respectively. After 72 h incubation cell viability was measured using CellTiter-Glo assay (Promega). Signals were normalized to control condition.

**Protein expression and purification**

The gene encoding NUDT5 (ENSG00000165609) (residues 1-208) was sub-cloned into a pNIC28-Bsa4 vector using ligation-independent cloning (LIC), resulting in an N-terminal 6x histidine tag followed by a TEV cleavage site. The plasmid was transformed into Escherichia coli BL21 (DE3), and protein expression was induced at an optical density (OD600) of 0.8 with 0.3 mM isopropyl-β-D-1-thiogalactopyranoside (IPTG) at 18 °C overnight. Cell pellets were resuspended in lysis buffer (20 mM HEPES, 500 mM NaCl, 5% glycerol, 0.5 mM TCEP, pH 7.5) supplemented with lysozyme, a protease inhibitor cocktail (Set III, EDTA-free, Calbiochem), and benzonase (2 µL of 10,000 U/µL, Merck). Cells were lysed using an ultrasonic cell disruptor (Vibra-Cell, Sonics).

The clarified supernatant was applied to a Ni-NTA affinity chromatography column (HisTrap Crude FF, GE Healthcare) and eluted with an increasing gradient of imidazole concentrations. The affinity-purified protein was incubated with TEV protease overnight at 4 °C to cleave the histidine tag. Following cleavage, reverse nickel affinity chromatography was performed to separate the cleaved tag from the protein. The protein was further purified using size-exclusion chromatography (16/600 Superdex 75 PG, GE Healthcare) in a final buffer consisting of 20 mM HEPES, pH 7.5, 500 mM NaCl, and 0.5 mM TCEP. The purity of the protein was confirmed by SDS-PAGE, and the protein was concentrated to 26 mg/mL using 10,000 MWCO Vivaspin concentrators (Vivascience).

**Crystallization**

Prior to crystallization trials, NUDT5 (26 mg/mL) was mixed with dNUDT5 at a 1:2 molar ratio in the presence of 2 mM MgCl2 and incubated on ice for 1 hour. Crystallization screens were set up using the sitting-drop vapor diffusion method, with equal volumes of protein and precipitant solutions (1:1 ratio). Co-crystals were grown in a buffer solution comprising 0.1 M Tris-HCl (pH 8.5), 30% PEG 4000, and 0.2 M sodium acetate. Crystals were cryo-protected with the crystallization condition containing 20% glycerol and flash-frozen in liquid nitrogen.

**Data Collection, Structure Determination, and Refinement**

X-ray datasets were collected at Diamond Light Source (Harwell, U.K.) on I03 beamline using on Eiger2 XE 16M detector. Data processing and scaling were performed was performed using the program DIALS in combination with XIA2[7]. The structure was solved in Molecular replacement in PHASER[8] using PDB 5NWH as the search model. The model was further built in Coot[9] and refined in PHENIX[10]. Data collection and refinement statistics are summarized in Table 1. Figure was prepared using ChimeraX[11]. Structural data generated in this study have been deposited in the Protein Data Bank (PDB) under accession code 9G6Y.

**Synthesis and compound characterization**

**General remarks**

Commercial reagents and solvents were purchased from commercial suppliers and used without further purification. All reactions involving moisture sensitive reagents were carried out under a nitrogen atmosphere using standard vacuum line techniques and dry solvents. An Elga DV 25 system was used for deionising water. Thin layer chromatography was performed on aluminium plates coated with 60 $F_{254}$ silica gel. Plates were visualised using UV light (254 nm). Flash column chromatography was performed on a Biotage Isolera one flash column chromatography platform. $^{1}H$ and $^{13}C$ NMR spectra were obtained using Bruker NMR spectrometers (400 MHz). The proton and carbon chemical shift values are reported in parts per million (ppm, δ scale) downfield from tetramethylsilane (TMS) and the indicated solvent. NMR spectra were processed and analyzed using MestReNova software. Spin multiplicities are given as s (singlet), d (doublet), t (triplet), q (quartet), dd (doublet of doublets), m (multiplet) and b (broad), coupling constants J are given in hertz (Hz), and signal area integration in natural numbers.

LC-MS was performed with a Kinetex 5µm EVO C18 100A 100 x 3.0 mm column on a Waters SFO and 515 HPLC pump and Waters Binary Gradient 2545 device using linear gradient of solvent A (93 % water, 5 % acetonitrile and 2 % of 0.5 M ammonium acetate pH 6.0) and solvent B (18 % $H_2O$, 80 % acetonitrile and 2 % ammonium acetate pH 6.0), eluting at a flow rate of 2 mL/min: 5 % B for 0.35 min, 5% B to 95% B for 1 min, 95% to 5% B for 0.1 min and 5% B for 0.8 min. LCMS was used as a measure of compound purity using either UV absorbance (Waters UV/visible Detector 2489), ELSD signal (Waters ELS Detector 2424) or ESI+ TIC (SQ Detector 2). Preparative HPLC was performed on the same system with a Kinetex 5µm EVO C18 100A 150 x 21.2 mm column using linear gradient of solvent A over 20 min from 85% to 10% eluting at a flow rate of 20 mL/min. LC-MS was acquired using Waters FractionLynx software and processed using MestReNova software. HRMS was acquired with Agilent 6530 RapidFire QTOF mass spectrometer in 384-well polypropylene plates (Greiner, 781280) using an assay buffer consisting of 10 mM ammonium formate pH 7.5. The plate was transferred to a RapidFire RF360 high-throughput sampling robot. Samples were aspirated under vacuum and loaded onto a C4 solid-phase extraction (SPE) cartridge equilibrated and washed for 4.5 s with 0.1% formic acid in LC-MS grade water to remove non-volatile buffer components. After aqueous wash, analytes of interest were eluted from the C4 SPE onto an Agilent 6530 accurate mass Q-TOF in an organic elution step, 85% acetonitrile and 0.1% formic acid in LC-MS grade water. Ion data for the compounds were extracted and compound formulas were generated using MassHunter Qualitative Analysis B.06.00 (Agilent). The purity for all final compounds was determined to be >95% by HPLC.

**General procedure 1**

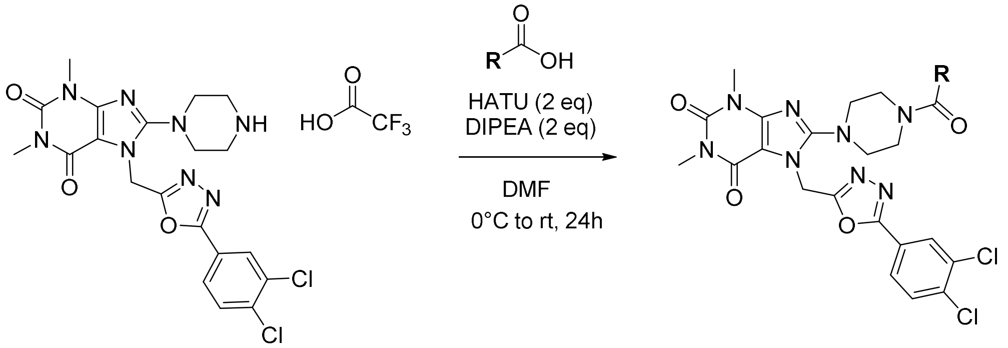

Under nitrogen, HATU (2 eq) was added to a solution of acid (1.1-1.2 eq) and DIPEA (2 or 3 eq) in DMF (0.03 M) at 0°C. The mixture was stirred for 15 min at 0°C. A solution of TH5427 TFA salt (1.2 eq) and DIPEA (1 eq) in DMF (0.03 M) was then added dropwise. The mixture was stirred for 24h at room temperature. After concentration under reduced pressure, the crude was purified by prep HPLC to afford the expected product.

**General procedure 2**

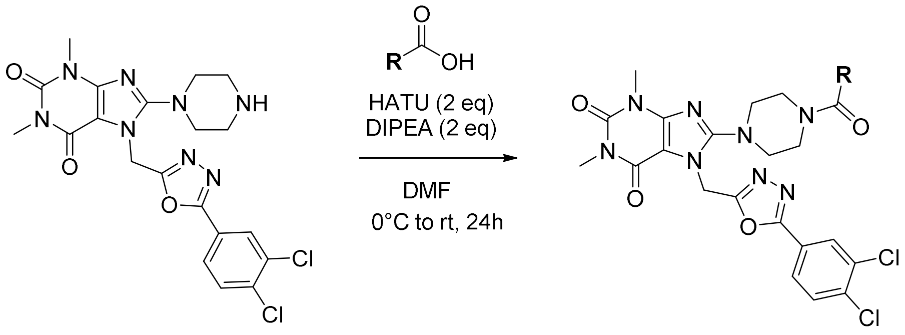

Under nitrogen, HATU was added to a solution of acid and DIPEA (2 or 3 eq) in DMF (0.010 M) at 0°C. The mixture was stirred for 15 min at 0°C. The amine was then added portion wise. The mixture was stirred for 24h at room temperature. After concentration under reduced pressure, the crude was purified by prep HPLC to afford the expected products.

**General procedure 3**

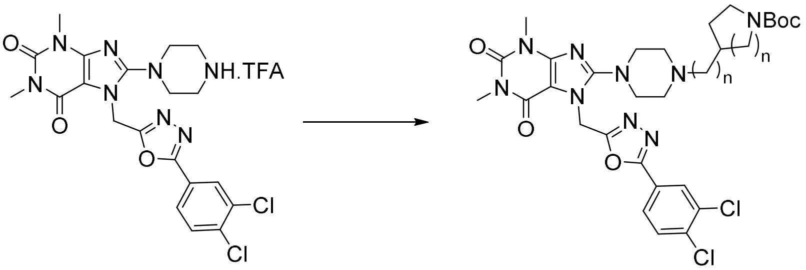

Under nitrogen, sodium triacetoxyborohydride was added to a solution of the corresponding aldehyde and TH5427 trifluoroacetic salt in THF. The resulting mixture was stirred at room temperature overnight. The crude was purified by flash column chromatography (Si-35, DCM/MeOH, gradient from 100/0 to 95/5) to afford the expected product.

**General procedure 4**

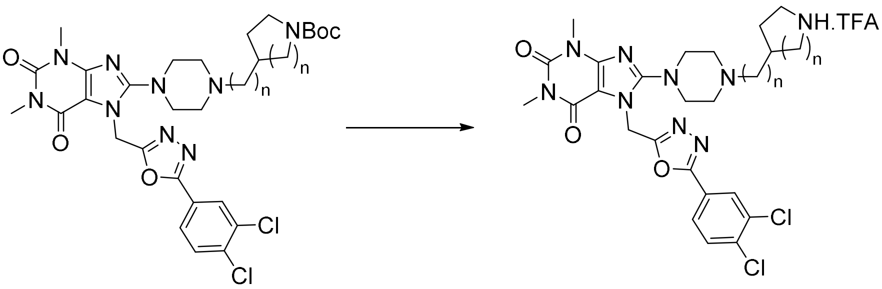

Under nitrogen, TFA was added to a solution of *N*-Boc protected compound in DCM at 0°C. The solution was stirred for the indicated time at room temperature. After completion, the crude was concentrated under reduced pressure and used in the next step without further purification.

**General procedure 5**

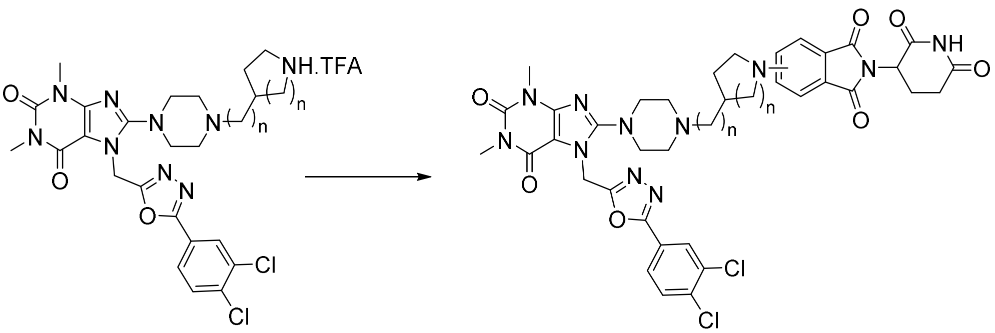

Under nitrogen, TEA (3 eq) was added to a solution of amine trifluoroacetic salt (1 eq) and fluoro thalidomide (1 eq) in DMSO. The mixture was heated under microwave at 200°C for 15-20 min. The crude was purified by prep HPLC to afford the expected product.

**General procedure 6**

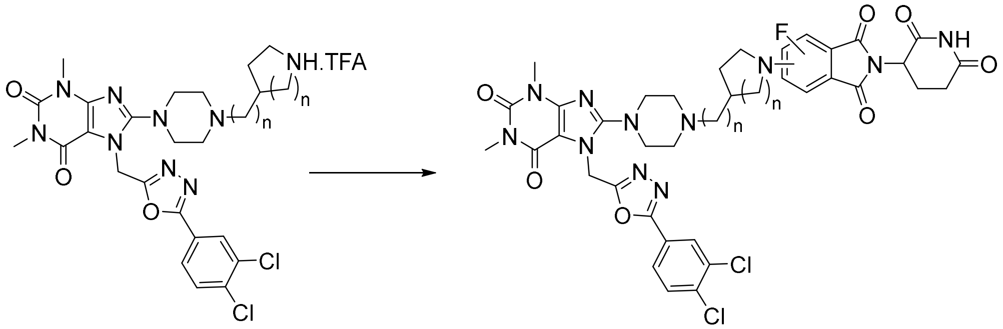

Under nitrogen, DIPEA (3 eq) was added to a solution of amine (1 eq) and mono- or di-fluoro thalidomide (1 eq) in DMSO. The mixture was heated at 100°C overnight. The crude was purified by prep HPLC to afford the expected product.

**ASM57**

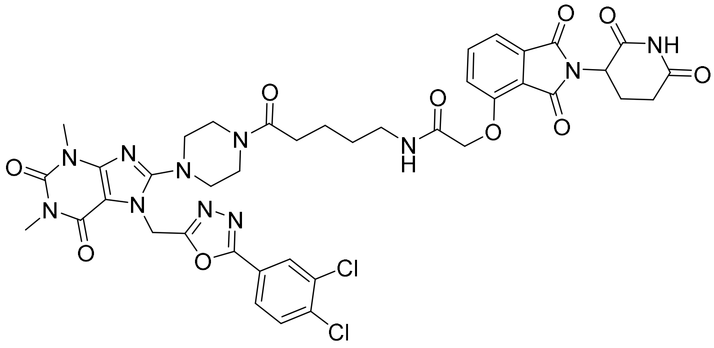

*N*-(5-(4-(7-((5-(3,4-dichlorophenyl)-1,3,4-oxadiazol-2-yl)methyl)-1,3-dimethyl-2,6-dioxo-2,3,6,7-tetrahydro-1*H*-purin-8-yl)piperazin-1-yl)-5-oxopentyl)-2-((2-(2,6-dioxopiperidin-3-yl)-1,3-dioxoisoindolin-4-yl)oxy)acetamide

ASM57 was prepared according to the general procedure 1 using TH5427 TFA salt (28.9 mg, 0.034 mmol, 1.1 eq), 5-(2-((2-(2,6-dioxopiperidin-3-yl)-1,3-dioxoisoindolin-4-yl)oxy)acetamido)pentanoic acid (10.3 mg, 0.031 mmol, 1 eq), DIPEA (22 µL, 0.12 mmol, 4 eq) and HATU (23.6 mg, 0.062 mmol, 2 eq). After purification, a white solid was obtained (3.3 mg, 0.024 mmol, 12%).

**$^{1}$H NMR (400 MHz, $CDCl_3$)** δ 8.99 (s, 1H), 8.10 (d, J = 2Hz, 1H), 7.85 (dd, J1 = 8.4 Hz, J2 = 2.0 Hz, 1H), 7.74 (dd, J1 = 7.4 Hz, J2 = 8.4 Hz, 1H), 7.68-7.65 (m, 1H), 7.60-7.54 (m, 2H), 7.21 (d, J = 8.4 Hz, 1H), 5.69 (s, 2H), 4.98-4.94 (m, 1H), 4.70-4.60 (m, 2H), 3.75-3.67 (m, 2H), 3.59-3.56 (m, 2H), 3.54 (s, 3H), 3.47-3.42 (m, 1H), 3.37 (s, 3H), 3.35-3.32 (m, 1H), 3.27-3.24 (m, 4H), 2.92-2.85 (m, 1H), 2.82-2.73 (m, 2H), 2.42-2.34 (m, 2H), 2.20-2.15 (m, 1H), 1.66-1.62 (m, 4H).

**$^{13}$C NMR (101 MHz, $CDCl_3$)** δ 171.6, 171.3, 168.4, 167.1, 166.7, 166.4, 164.2, 162.0, 155.9, 155.0, 154.9, 151.7, 147.7, 137.3, 137.0, 134.0, 133.7, 131.5, 128.9, 126.2, 123.1, 120.4, 118.5, 117.8, 105.0, 68.8, 50.6 (2C), 49.5, 45.0, 40.1, 39.0, 32.8, 31.5, 30.2, 30.0, 29.1, 28.0, 22.9, 22.5.

**HPLC-MS** $t_R$ = 1.475 min (93.2 %), ESI+ m/z 904.630; 906.071 $[M+H]^+$

**HRMS** was calculated for $C_{40}H_{40}Cl_2N_{11}O_{10}$: 904.2337, found: 906.2332

**ASM68**

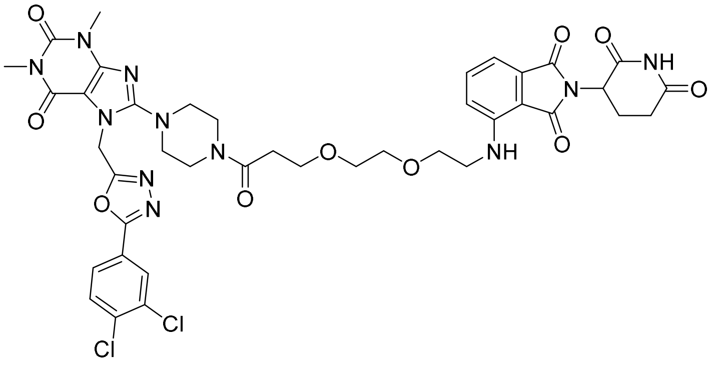

7-((5-(3,4-dichlorophenyl)-1,3,4-oxadiazol-2-yl)methyl)-8-(4-(3-(2-(2-((2-(2,6-dioxopiperidin-3-yl)-1,3-dioxoisoindolin-4-yl)amino)ethoxy)ethoxy)propanoyl)piperazin-1-yl)-1,3-dimethyl-3,7-dihydro-1*H*-purine-2,6-dione

ASM68 was prepared according to the general procedure 1 using TH5427 TFA salt (20 mg, 0.033 mmol, 1.1 eq), 3-(2-(2-((2-(2,6-dioxopiperidin-3-yl)-1,3-dioxoisoindolin-4-yl)amino)ethoxy)ethoxy) propanoic acid (13 mg, 0.030 mmol, 1 eq), DIPEA (21 µL, 0.12 mmol, 4 eq) and HATU (23 mg, 0.060 mmol, 2 eq). After purification, a yellow solid was obtained (21.4 mg, 0.024 mmol, 79%).

**$^{1}$H NMR (400 MHz, CDCl$_3$)** δ 8.88 (s, 1H), 8.07 (d, J = 2.0 Hz, 1H), 7.83 (dd, J1 = 8.4 Hz, J2 = 2.0 Hz, 1H), 7.57 (d, J = 8.4 Hz, 1H), 7.45 (dd, J1 = 7.0 Hz, J2 = 8.4 Hz, 1H), 7.06 (d, J = 7.0 Hz, 1H), 6.88 (d, J = 8.4 Hz, 1H), 6.45 (bs, 1H), 5.68 (s, 2H), 4.93-4.88 (m, 1H), 3.79 (t, J = 6.5 Hz, 2H), 3.71 (bt, J = 5.3 Hz, 4H), 3.65-3.63 (m, 4H), 3.61-3.59 (m, 2H), 3.53 (s, 3H), 3.46-3.42 (m, 2H), 3.35 (s, 3H), 3.29-3.23 (m, 4H), 2.88-2.81 (m, 1H), 2.77-2.72 (m, 2H), 2.64-2.60 (m, 2H), 2.17-2.10 (m, 1H).

**$^{13}$C NMR (101 MHz, CDCl$_3$)** δ 171.5, 170.1, 169.4, 168.8, 167.6, 164.1, 162.1, 155.9, 155.0, 151.6, 147.7, 146.88, 136.9, 136.1, 133.9, 132.5, 131.5, 128.8, 126.1, 123.0, 116.9, 111.7, 110.3, 104.9, 70.6, 70.5, 69.5, 67.7, 50.8, 50.3, 49.00, 45.2, 42.5, 41.0, 40.1, 33.7, 31.5, 30.0, 28.0, 22.9.

**HPLC-MS** $t_R$ = 1.54 min (100 %), ESI+ m/z 907.379 [M+H]$^+$

**HRMS** was calculated for $C_{40}H_{42}Cl_2N_{11}O_{10}$: 906.2493, found: 906.2487

**ASM76**

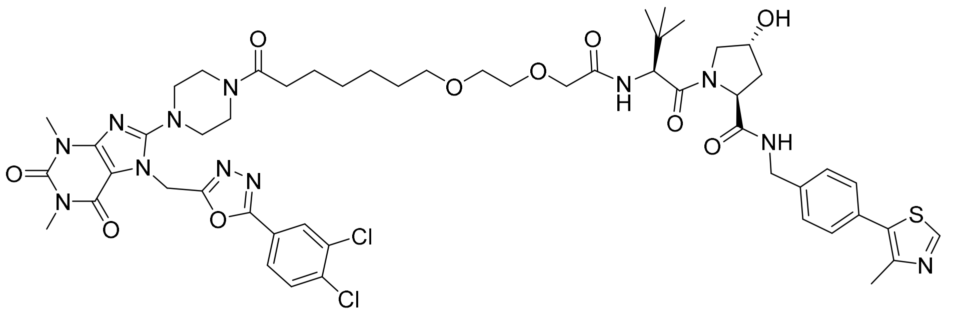

(2*S*,4*R*)-1-((*S*)-2-(2-(2-((7-(4-(7-((5-(3,4-dichlorophenyl)-1,3,4-oxadiazol-2-yl)methyl)-1,3-dimethyl-2,6-dioxo-2,3,6,7-tetrahydro-1*H*-purin-8-yl)piperazin-1-yl)-7-oxoheptyl)oxy)ethoxy)acetamido)-3,3-dimethylbutanoyl)-4-hydroxy-*N*-(4-(4-methylthiazol-5-yl)benzyl)pyrrolidine-2-carboxamide

ASM76 was prepared according to the general procedure 1 using TH5427 TFA salt (12 mg, 0.020 mmol, 1.1 eq), 7-(2-(2-(((S)-1-((2S,4R)-4-hydroxy-2-((4-(4-methylthiazol-5-yl)benzyl)carbamoyl) pyrrolidin-1-yl)-3,3-dimethyl-1-oxobutan-2-yl)amino)-2-oxoethoxy)ethoxy)heptanoic acid (12 mg, 0.018 mmol, 1 eq), DIPEA (9 µL, 0.054 mmol, 3 eq) and HATU (13.8 mg, 0.036 mmol, 2 eq). After purification, a white solid was obtained (12 mg, 0.011 mmol, 58%).

**$^{1}$H NMR (400 MHz, DMSO-*d6*)** δ (major rotamer described) 8.96 (s, 1H), 8.59 (t, J = 6 Hz, 1H), 8.14 (bd, J = 1.9 Hz, 1H) 7.93 (dd, J1 = 1.9 Hz, J2 = 8.5 Hz, 1H), 7.87 (d, J = 8.5 Hz, 1H), 7.42-7.36 (m, 5H), 5.72 (s, 2H), 5.15 (s, 1H), 4.55 (d, J = 9.5 Hz, 1H), 4.46-4.35 (m, 3H), 4.28-4.20 (m, 1H), 3.95 (s, 2H), 3.68-3.63 (m, 1H), 3.62-3.59 (m, 3H), 3.56-3.50 (m, 7H), 3.39 (s, 3H), 3.26-3.19 (m, 4H), 3.14 (s, 3H), 2.42 (s, 3H), 2.27 (t, J = 7.5 Hz, 2H), 2.08-2.03 (m, 1H), 1.93-1.87 (m, 1H), 1.49-1.40 (m, 4H), 1.26-1.22 (m, 4H), 0.94 (s, 9H). *OH not observed*

**$^{13}$C NMR (400 MHz, DMSO-*d6*)** δ 171.8, 170.9, 169.1, 168.6, 163.1, 162.7, 155.9, 153.8, 151.4, 150.9, 147.7, 147.2, 139.4, 135.0, 132.5, 132.0, 131.2, 129.7, 128.9, 128.7 (2C), 128.1, 127.5 (2C), 126.6, 123.5, 104.1, 70.6, 70.5, 69.6, 69.2, 68.9, 58.8, 56.6, 55.7, 50.0, 49.7, 44.4, 41.7, 40.4, 37.9, 35.7, 32.2, 29.6, 29.1, 28.6, 27.4, 26.2 (3C), 25.5, 24.7, 15.9.

**HPLC-MS** $t_R$ = 1.59 min (99.3 %), ESI+ m/z 1134.809; 1136.456 $[M+H]^+$

**ASM84**

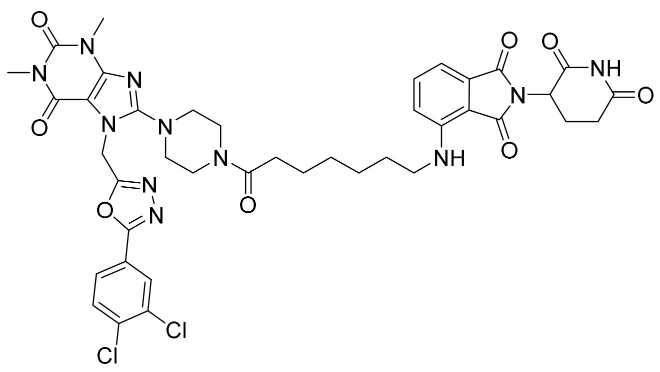

7-((5-(3,4-dichlorophenyl)-1,3,4-oxadiazol-2-yl)methyl)-8-(4-(7-((2-(2,6-dioxopiperidin-3-yl)-1,3-dioxoisoindolin-4-yl)amino)heptanoyl)piperazin-1-yl)-1,3-dimethyl-3,7-dihydro-1*H*-purine-2,6-dione

ASM84 was prepared according to the general procedure 1 using TH5427 TFA salt (16.6 mg, 0.027 mmol, 1.1 eq), 7-((2-(2,6-dioxopiperidin-3-yl)-1,3-dioxoisoindolin-4-yl)amino)heptanoic acid (10 mg, 0.025 mmol, 1 eq), DIPEA (13 µL, 0.075 mmol, 3 eq) and HATU (18.9 mg, 0.050 mmol, 2 eq). After purification, a yellow solid was obtained (18.2 mg, 0.021 mmol, 84%).

**$^1$H NMR (400 MHz, $CDCl_3$)** δ 8.46 (s, 1H), 8.10 (d, J = 2Hz, 1H), 7.86 (dd, J1 = 2.0 Hz, J2 = 8.5 Hz, 1H), 7.60 (d, J = 8.5 Hz, 1H) 7.49 (dd, J1 = 7.0 Hz, J2 = 8.4 Hz, 1H) 7.08 (d, J = 7.0 Hz, 1H) 6.88 (d, J = 8.4 Hz, 1H), 6.22 (d, J = 5.7 Hz, 1H), 5.70 (s, 2H), 4.93-4.89 (m, 1H), 3.74-3.71 (m, 2H), 3.60-3.57 (m, 2H), 3.55 (s, 3H), 3.37 (s, 3H), 3.29- 3.24 (m, 6H), 2.90-2.84 (m, 1H), 2.81-2.69 (m, 2H), 2.34 (t, J = 7.3 Hz, 2H), 2.15-2.10 (m, 1H), 1.71-1.63 (m, 4H), 1.49-1.36 (m, 4H).

**$^{13}$C NMR (101 MHz, $CDCl_3$)** δ 171.7, 171.2, 169.6, 168.6, 167.7, 164.2, 162.0, 155.9, 155.0, 151.7, 147.7, 147.1, 137.0, 136.3, 134.0, 132.6, 131.5, 128.9, 126.2, 123.0, 116.8, 111.5, 110.0, 105.0, 50.7, 50.6, 49.0, 45.0, 42.6, 40.9, 40.1, 33.1, 31.5, 30.0, 29.1, 29.1, 28.0, 26.8, 25.0, 22.9.

**HPLC-MS** $t_R$ = 1.68 min (99 %), ESI+ m/z 874.120 $[M+H]^+$

**ASM111**

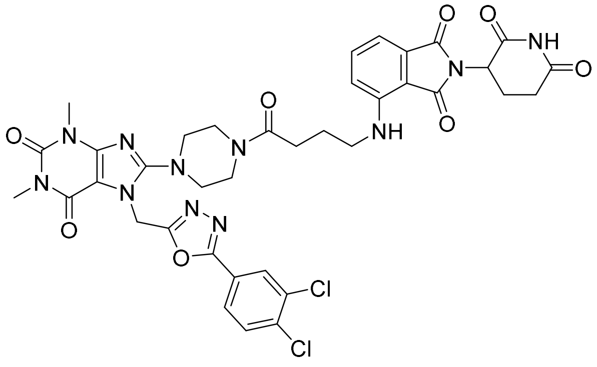

7-((5-(3,4-dichlorophenyl)-1,3,4-oxadiazol-2-yl)methyl)-8-(4-(4-((2-(2,6-dioxopiperidin-3-yl)-1,3-dioxoisoindolin-4-yl)amino)butanoyl)piperazin-1-yl)-1,3-dimethyl-3,7-dihydro-1*H*-purine-2,6-dione

ASM111 was prepared according to the general procedure 1 using TH5427 TFA salt (27.8 mg, 0.046 mmol, 1.1 eq), 4-((2-(2,6-dioxopiperidin-3-yl)-1,3-dioxoisoindolin-4-yl)amino)butanoic acid (15 mg, 0.042 mmol, 1 eq), DIPEA (22 µL, 0.125 mmol, 3 eq) and HATU (31.7 mg, 0.083 mmol, 2 eq). After purification, a yellow solid was obtained (10.1 mg, 0.012 mmol, 29%).

**$^{1}$H NMR (400 MHz, $CDCl_3$)** δ 8.44 (s, 1H), 8.12 (d, J = 2Hz, 1H), 7.87 (dd, J1 = 2.0 Hz, J2 = 8.4 Hz, 1H), 7.61 (d, J = 8.4 Hz, 1H) 7.51 (dd, J1 = 7.0 Hz, J2 = 8.4 Hz, 1H) 7.11 (d, J = 7.0 Hz, 1H) 6.98 (d, J = 8.4 Hz, 1H), 5.70 (S, 2H), 4.93-4.89 (m, 1H), 3.81-3.69 (m, 2H), 3.63-3.56 (m, 2H), 3.56 (s, 3H), 3.40 (t, J = 6.8 Hz, 2H), 3.38 (s, 3H), 3.30- 3.25 (m, 4H), 2.91-2.77 (m, 2H), 2.73-2.67 (m, 1H), 2.45 (t, J = 6.8 Hz, 2H), 2.17-2.12 (m, 1H), 2.09-2.03 (m, 2H). *NH-Ar not observed*

**$^{13}$C NMR (101 MHz, $CDCl_3$)** δ 171.2, 170.7, 169.6, 168.7, 167.7, 164.2, 162.0, 155.8, 155.0, 151.7, 147.7, 147.0, 137.0, 136.4, 134.0, 132.6, 131.5, 128.9, 126.2, 123.0, 117.1, 111.9, 110.4, 105.0, 50.6, 50.5, 49.0, 44.9, 42.2, 41.1, 40.1, 31.6, 30.0, 29.9, 28.0, 24.2, 23.0.

**ASM122**

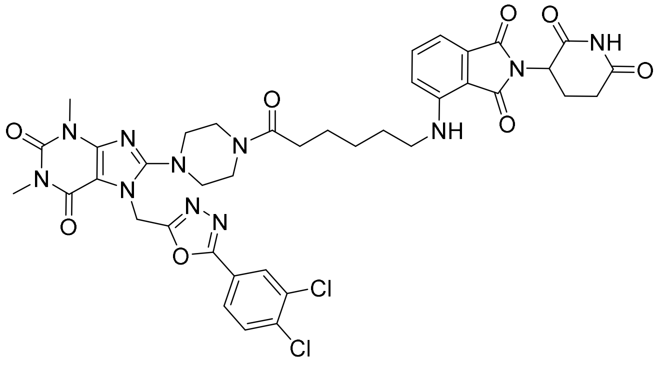

7-((5-(3,4-dichlorophenyl)-1,3,4-oxadiazol-2-yl)methyl)-8-(4-(6-((2-(2,6-dioxopiperidin-3-yl)-1,3-dioxoisoindolin-4-yl)amino)hexanoyl)piperazin-1-yl)-1,3-dimethyl-3,7-dihydro-1*H*-purine-2,6-dione

ASM122 was prepared according to the general procedure 1 using TH5427 TFA salt (33.2 mg, 0.055 mmol, 1.1 eq), 6-((2-(2,6-dioxopiperidin-3-yl)-1,3-dioxoisoindolin-4-yl)amino)hexanoic acid (25 mg, 0.050 mmol, 1 eq), DIPEA (26 μL, 0.15 mmol, 3 eq) and HATU (37.9 mg, 0.1 mmol, 2 eq). After purification, a yellow solid was obtained (8 mg, 0.009 mmol, 19%).

**$^1$H NMR (400 MHz, $CDCl_3$)** δ 8.19 (s, 1H), 8.11 (d, J = 2.0 Hz, 1H), 7.86 (dd, J = 2.0 Hz, J = 8.5 Hz, 1H), 7.60 (d, J = 8.5 Hz, 1H), 7.49 (dd, J = 7.1 Hz, J = 8.5 Hz, 1H), 7.09 (d, J = 7.1 Hz, 1H), 6.88 (d, J = 8.5 Hz, 1H), 6.23 (s, 1H), 5.69 (s, 2H), 4.93-4.88 (m, 1H), 3.75-3.72 (m, 2H), 3.59-3.57 (m, 2H), 3.55 (s, 3H), 3.37 (s, 3H), 3.30-3.24 (m, 6H), 2.91-2.86 (m, 1H), 2.84-2.78 (m, 1H), 2.76-2.71 (m, 1H), 2.36 (t, J = 7.3 Hz, 2H), 2.16-2.11 (m, 1H), 1.74-1.67 (m, 4H), 1.51-1.43 (m, 2H).

**$^{13}$C NMR (101 MHz, $CDCl_3$)** δ 171.5, 171.1, 169.7, 168.6, 167.7, 164.2, 162.0, 155.9, 155.1, 151.7, 147.8, 147.1, 137.0, 136.3, 134.0, 132.6, 131.5, 128.9, 126.2, 123.1, 116.8, 111.6, 110.0, 105.0, 50.7, 50.6, 49.0, 45.0, 42.4, 41.0, 40.1, 33.1, 31.6, 30.0, 29.1, 28.0, 26.8, 24.8, 23.0.

**ASM127**

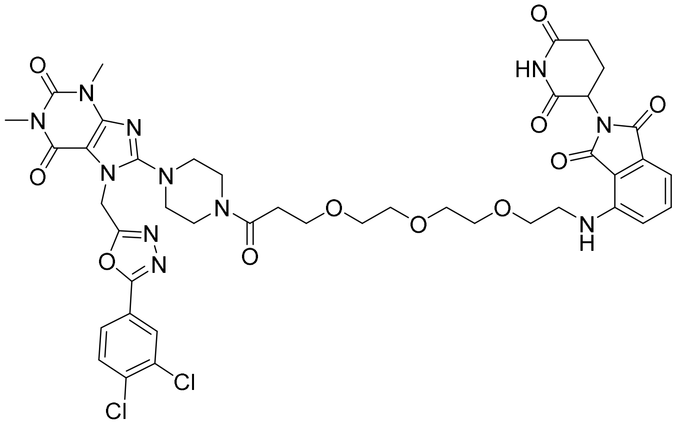

7-((5-(3,4-dichlorophenyl)-1,3,4-oxadiazol-2-yl)methyl)-8-(4-(3-(2-(2-(2-((2-(2,6-dioxopiperidin-3-yl)-1,3-dioxoisoindolin-4-yl)amino)ethoxy)ethoxy)ethoxy)propanoyl)piperazin-1-yl)-1,3-dimethyl-3,7-dihydro-1*H*-purine-2,6-dione

ASM127 was prepared according to the general procedure 2 using TH5427 (26.9 mg, 0.055 mmol, 1.1 eq), 3-(2-(2-(2-((2-(2,6-dioxopiperidin-3-yl)-1,3-dioxoisoindolin-4-yl)amino)ethoxy)ethoxy)ethoxy) propanoic acid (23.8 mg, 0.050 mmol, 1 eq), DIPEA (17 µL, 0.1 mmol, 2 eq) and HATU (37.9 mg, 0.1 mmol, 2 eq). After purification, a yellow solid was obtained (14.9 mg, 0.016 mmol, 31%).

**$^{1}$H NMR (400 MHz, CDCl$_3$)** δ 8.86 (s, 1H), 8.10 (d, J = 2Hz, 1H), 7.86 (dd, J1 = 2.0 Hz, J2 = 8.5 Hz, 1H), 7.59 (d, J = 8.5 Hz, 1H) 7.48 (dd, J1 = 7.0 Hz, J2 = 8.5 Hz, 1H) 7.09 (d, J = 7.0 Hz, 1H) 6.90 (d, J = 8.5 Hz, 1H), 5.68 (s, 2H), 4.94-4.89 (m, 1H), 3.80 (t, J = 6.5, 2H), 3.75-3.70 (m, 4H), 3.68-3.60 (m, 10H), 3.54 (s, 3H), 3.47-3.44 (m, 2H), 3.37 (s, 3H), 3.31-3.24 (m, 4H), 2.89-2.82 (m, 1H), 2.79-2.72 (m, 2H), 2.64 (t, J = 6.5, 2H), 2.17-2.10 (m, 1H). *NH-Ar not observed*

**$^{13}$C NMR (101 MHz, CDCl$_3$)** δ 171.5, 170.2, 169.4, 168.7, 167.7, 164.1, 162.1, 155.9, 155.0, 151.7, 147.8, 146.9, 136.9, 136.1, 134.0, 132.7, 131.5, 128.9, 126.2, 123.1, 116.8, 111.8, 110.5, 104.9, 71.0, 70.7, 70.6, 70.5, 69.5, 67.5, 50.6, 50.5, 49.0, 45.2, 42.5, 41.0, 40.1, 33.7, 31.6, 30.0, 28.0, 23.0.

**HPLC-MS** $t_R$ = 1.56 min (96 %), ESI+ m/z 951.535 $[M+H]^+$

**HRMS** was calculated for $C_{42}H_{46}Cl_2N_{11}O_{11}$: 950.2755, found: 950.2727

**ASM130**

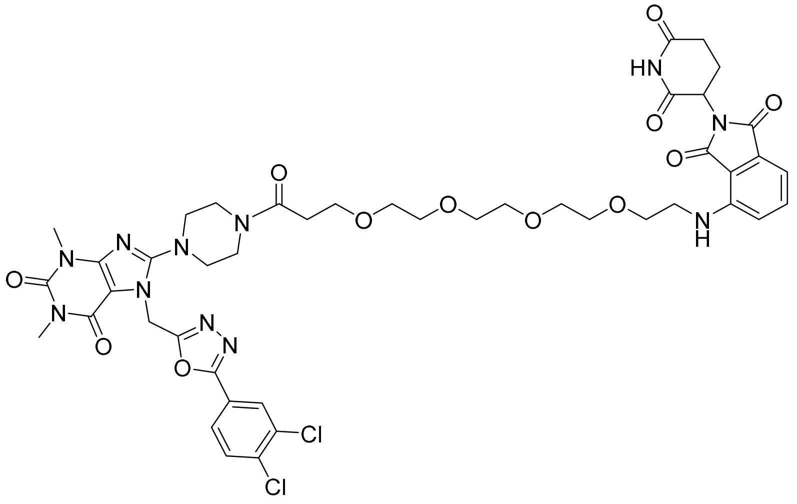

7-((5-(3,4-dichlorophenyl)-1,3,4-oxadiazol-2-yl)methyl)-8-(4-(1-((2-(2,6-dioxopiperidin-3-yl)-1,3-dioxoisoindolin-4-yl)amino)-3,6,9,12-tetraoxapentadecan-15-oyl)piperazin-1-yl)-1,3-dimethyl-3,7-dihydro-1*H*-purine-2,6-dione

ASM130 was prepared according to the general procedure 2 using TH5427 (10.3 mg, 0.021 mmol, 1.1 eq), 1-((2-(2,6-dioxopiperidin-3-yl)-1,3-dioxoisoindolin-4-yl)amino)-3,6,9,12-tetraoxa pentadecan-15-oic acid (9.9 mg, 0.019 mmol, 1 eq), DIPEA (10 µL, 0.057 mmol, 3 eq) and HATU (14.4 mg, 0.038 mmol, 2 eq). After purification, a yellow solid was obtained (4.9 mg, 0.005 mmol, 26%).

**$^1$H NMR (400 MHz, $CDCl_3$)** δ 8.57 (s, 1H), 8.11 (d, J = 2Hz, 1H), 7.86 (dd, J1 = 2.0 Hz, J2 = 8.5 Hz, 1H), 7.60 (d, J = 8.5 Hz, 1H) 7.49 (dd, J1 = 7.0 Hz, J2 = 8.5 Hz, 1H) 7.10 (d, J = 7.0 Hz, 1H) 6.92 (d, J = 8.5 Hz, 1H), 5.69 (s, 2H), 4.93-4.89 (m, 1H), 3.81 (t, J = 6.5, 2H), 3.74-3.71 (m, 4H), 3.68-3.64 (m, 8H), 3.64-3.61 (m, 6H), 3.56 (s, 3H), 3.47 (t, J = 5.5 Hz, 2H), 3.38 (s, 3H), 3.30-3.27 (m, 4H), 2.90-2.84 (m, 1H), 2.81-2.72 (m, 2H), 2.65 (t, J = 6.5, 2H), 2.16-2.10 (m, 1H). *NH-Ar not observed*

**$^{13}$C NMR (101 MHz, $CDCl_3$)** δ 171.3, 170.0, 169.4, 168.6, 167.7, 164.1, 162.1, 155.9, 155.0, 151.7, 147.8, 146.9, 136.9, 136.2, 134.0, 132.7, 131.5, 128.9, 126.2, 123.1, 116.9, 111.8, 110.5, 105.0, 70.9, 70.8, 70.7, 70.6, 70.6, 70.5, 69.6, 67.5, 50.7, 50.5, 49.0, 45.2, 42.5, 41.0, 40.1, 33.7, 31.6, 30.0, 28.0, 23.0.

**HPLC-MS** $t_R$ = 1.55 min (97.8 %), ESI+ m/z 995.315; 996.395 $[M+H]^+$

**HRMS** was calculated for $C_{44}H_{50}Cl_2N_{11}O_{12}$: 994.3018, found: 994.3007

**ASM131**

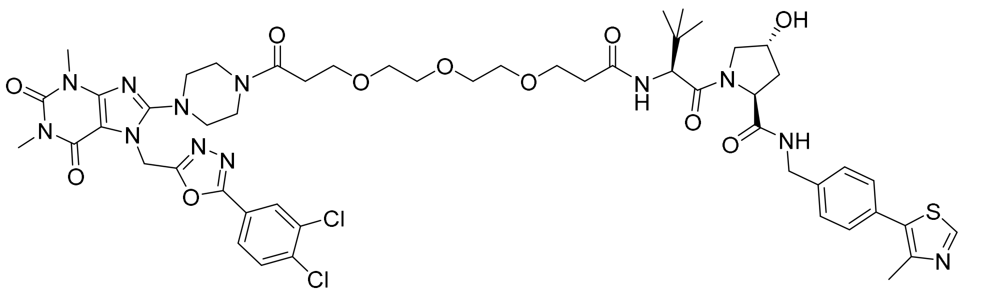

(2*S*,4*R*)-1-((*S*)-2-(*tert*-butyl)-16-(4-(7-((5-(3,4-dichlorophenyl)-1,3,4-oxadiazol-2-yl)methyl)-1,3-dimethyl-2,6-dioxo-2,3,6,7-tetrahydro-1*H*-purin-8-yl)piperazin-1-yl)-4,16-dioxo-7,10,13-trioxa-3-azahexadecanoyl)-4-hydroxy-N-(4-(4-methylthiazol-5-yl)benzyl)pyrrolidine-2-carboxamide

ASM131 was prepared according to the general procedure 2 using TH5427 (10.8 mg, 0.022 mmol, 1.1 eq), (*S*)-15-((2*S*,4*R*)-4-hydroxy-2-((4-(4-methylthiazol-5-yl)benzyl)carbamoyl)pyrrolidine-1-carbonyl)-16,16-dimethyl-13-oxo-4,7,10-trioxa-14-azaheptadecanoic acid (13.3 mg, 0.020 mmol, 1 eq), DIPEA (10 μL, 0.057 mmol, 3 eq) and HATU (15.2 mg, 0.040 mmol, 2 eq). After purification, a yellow solid was obtained (5.2 mg, 0.005 mmol, 24%).

**$^{1}$H NMR (400 MHz, $CDCl_3$)** δ 8.87 (s, 1H), 8.11 (d, J = 2 Hz, 1H), 7.86 (dd, J1 = 2.0 Hz, J2 = 8.5 Hz, 1H), 7.60 (d, J = 8.5 Hz, 1H), 7.50-7.47 (m, 1H), 7.40-7.35 (m, 5H), 7.02 (d, J = 8.4 Hz, 1H), 5.70 (s, 2H), 4.72 (t, J = 8.1 Hz, 1H), 4.62-4.57 (m, 1H), 4.53-4.50 (m, 1H), 4.48 (d, J = 8.5 Hz, 1H), 4.37-4.32 (m, 1H), 4.13 (bd, J = 11.6 Hz, 1H), 3.76 (t, J = 6.6 Hz, 2H), 3.74-3.70 (m, 4H), 3.64-3.61 (m, 6H), 3.60-3.58 (m, 5H), 3.55 (s, 3H), 3.37 (s, 3H), 3.32-3.29 (m, 2H), 3.27-3.24 (m, 2H), 2.66-2.62 (m, 2H), 2.57 (s, 3H), 2.53-2.45 (m, 4H), 2.19-2.13 (m, 1H).

**$^{13}$C NMR (101 MHz, $CDCl_3$)** δ 172.2, 171.9, 171.2, 170.0, 164.2, 162.1, 155.9, 155.0, 151.7, 151.2, 147.8, 145.9, 139.6, 137.0, 134.0, 131.6, 129.5 (2C), 128.9, 128.5 (2C), 126.2, 123.1, 105.0, 70.6, 70.6, 70.5, 70.5, 70.3, 67.4, 67.3, 58.6, 57.9, 56.9, 50.7, 50.5, 45.2, 43.2, 41.0, 40.2, 36.8, 36.3, 35.0, 33.7, 30.0, 28.0, 26.6 (3C), 15.0.

**HPLC-MS** $t_R$ = 1.55 min (97.8 %), ESI+ m/z 995.315 $[M+H]^+$

**HRMS** was calculated for $C_{52}H_{65}Cl_2N_{12}O_{11}S$: 1135.3994, found: 1135.3998

**ASM133**

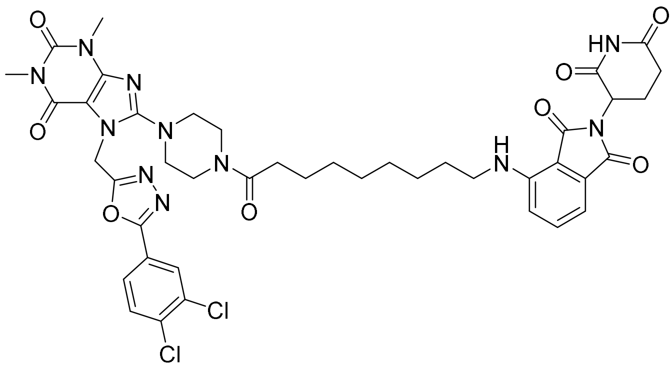

7-((5-(3,4-dichlorophenyl)-1,3,4-oxadiazol-2-yl)methyl)-8-(4-(9-((2-(2,6-dioxopiperidin-3-yl)-1,3-dioxoisoindolin-4-yl)amino)nonanoyl)piperazin-1-yl)-1,3-dimethyl-3,7-dihydro-1*H*-purine-2,6-dione

ASM133 was prepared according to the general procedure 2 using TH5427 (15.1 mg, 0.031 mmol, 1.1 eq), 9-((2-(2,6-dioxopiperidin-3-yl)-1,3-dioxoisoindolin-4-yl)amino)nonanoic acid (12.0 mg, 0.028 mmol, 1 eq), DIPEA (15 µL, 0.084 mmol, 3 eq) and HATU (21.2 mg, 0.056 mmol, 2 eq). After purification, a yellow solid was obtained (2.4 mg, 0.003 mmol, 10%).

**$^{1}$H NMR (400 MHz, $CDCl_3$)** δ 8.14 (s, 1H), 8.11 (d, J = 2.0 Hz, 1H), 7.86 (dd, J1 = 2.0 Hz, J2 = 8.5 Hz, 1H), 7.60 (d, J = 8.3 Hz, 1H), 7.51-7.47 (m, 1H), 7.08 (d, J = 7.0 Hz, 1H), 6.88 (d, J = 8.5 Hz, 1H), 5.69 (s, 2H), 4.93-4.89 (m, 1H), 3.71-3.62 (m, 4H), 3.55 (s, 3H), 3.37 (s, 3H), 3.29-3.24 (m, 6H), 2.91-2.80 (m, 2H), 2.78-2.68 (m, 2H), 2.33 (t, J = 7.3 Hz, 2H), 2.16-2.09 (m, 1H), 1.68-1.64 (m, 12H).

**HPLC-MS** $t_R$ = 1.79 min (97.7 %), ESI+ m/z 903.250 $[M+H]^+$

**ASM134**

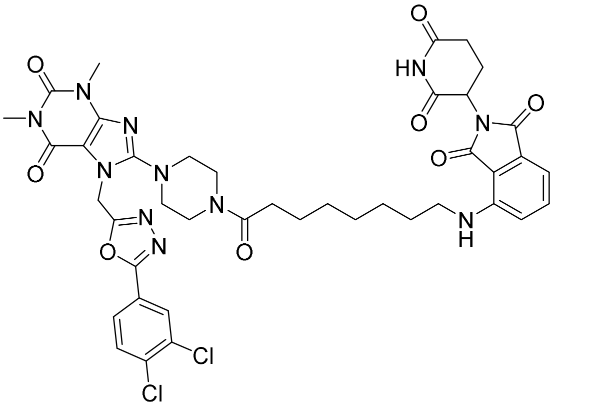

7-((5-(3,4-dichlorophenyl)-1,3,4-oxadiazol-2-yl)methyl)-8-(4-(8-((2-(2,6-dioxopiperidin-3-yl)-1,3-dioxoisoindolin-4-yl)amino)octanoyl)piperazin-1-yl)-1,3-dimethyl-3,7-dihydro-1*H*-purine-2,6-dione

ASM134 was prepared according to the general procedure 2 using TH5427 (15.0 mg, 0.030 mmol, 1.1 eq), 8-((2-(2,6-dioxopiperidin-3-yl)-1,3-dioxoisoindolin-4-yl)amino)octanoic acid (11.5 mg, 0.028 mmol, 1 eq), DIPEA (14 µL, 0.083 mmol, 3 eq) and HATU (21.1 mg, 0.055 mmol, 2 eq). After purification, a yellow solid was obtained (4.8 mg, 0.005 mmol, 19%).

**$^{1}$H NMR (400 MHz, $CDCl_3$)** δ 8.15 (s, 1H), 8.11 (d, J = 2.0 Hz, 1H), 7.86 (dd, J = 2.0 Hz, J = 8.4 Hz, 1H), 7.60 (d, J = 8.4 Hz, 1H), 7.49 (dd, J = 7.2 Hz, J = 8.4 Hz, 1H), 7.08 (d, J = 7.2 Hz, 1H), 6.88 (d, J = 8.6 Hz, 1H), 5.69 (s, 2H), 4.93-4.88 (m, 1H), 3.72-3.62 (m, 4H), 3.55 (s, 3H), 3.37 (s, 3H), 3.29- 3.24 (m, 6H), 2.91-2.80 (m, 2H), 2.78-2.68 (m, 2H), 2.33 (t, J = 7.4 Hz, 2H), 2.16-2.11 (m, 1H), 1.68-1.65 (m, 4H), 1.41-1.37 (m, 6H).

**$^{13}$C NMR (101 MHz, $CDCl_3$)** δ 171.9, 171.0, 169.7, 168.5, 167.7, 164.2, 162.0, 155.9, 155.1, 151.7, 147.8, 147.1, 137.0, 136.3, 134.0, 132.6, 131.6, 128.9, 126.2, 123.1, 116.8, 111.6, 110.0, 105.0, 50.7 (2C), 49.0, 45.0, 42.8, 40.9, 40.1, 33.3, 31.6, 30.0, 29.4, 29.3, 29.2, 28.0, 26.9, 25.2, 23.0.

**HPLC-MS** $t_R$ = 1.74 min (97.8 %), ESI+ m/z 888.726; 890.467 $[M+H]^+$

**HRMS** was calculated for $C_{41}H_{44}Cl_2N_{11}O_8$: 888.2751, found: 888.2767

**ASM135**

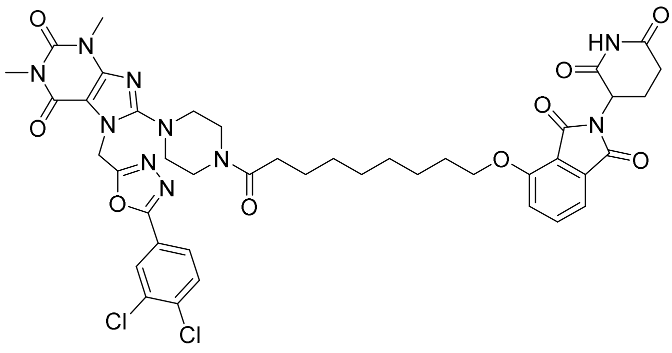

7-((5-(3,4-dichlorophenyl)-1,3,4-oxadiazol-2-yl)methyl)-8-(4-(9-((2-(2,6-dioxopiperidin-3-yl)-1,3-dioxoisoindolin-4-yl)oxy)nonanoyl)piperazin-1-yl)-1,3-dimethyl-3,7-dihydro-1*H*-purine-2,6-dione

ASM135 was prepared according to the general procedure 2 using TH5427 (15.1 mg, 0.031 mmol, 1.1 eq), 9-((2-(2,6-dioxopiperidin-3-yl)-1,3-dioxoisoindolin-4-yl)oxy)nonanoic acid (12 mg, 0.028 mmol, 1 eq), DIPEA (15 µL, 0.084 mmol, 3 eq) and HATU (21.2 mg, 0.056 mmol, 2 eq). After purification, a white solid was obtained (6.1 mg, 0.007 mmol, 24%).

**$^{1}$H NMR (400 MHz, $CDCl_3$)** δ 8.33 (s, 1H), 8.10 (d, J = 2Hz, 1H), 7.86 (dd, J1 = 2.0 Hz, J2 = 8.5 Hz, 1H), 7.66 (dd, J1 = 7.3 Hz, J2 = 8.5 Hz, 1H), 7.60 (d, J = 8.5 Hz, 1H), 7.44 (bd, J = 7.4 Hz, 1H) 7.20 (d, J = 8.4 Hz, 1H) 5.69 (s, 2H), 4.97-4.92 (m, 1H), 4.19-4.14 (m, 2H), 3.72 (bs, 2H), 3.59 (bs, 2H), 3.55 (s, 3H), 3.37 (s, 3H), 3.29-3.19 (m, 4H), 2.91-2.77 (m, 2H), 2.76-2.67 (m, 1H), 2.33 (t, J = 7.3 Hz, 2H), 2.16-2.10 (m, 1H), 1.90-1.83 (m, 2H), 1.74 (bs, 2H), 1.64-1.61 (m, 2H), 1.53-1.47 (m, 2H), 1.39-1.36 (m, 4H).

**$^{13}$C NMR (101 MHz, $CDCl_3$)** δ 172.1, 171.1, 168.3, 167.2, 165.8, 164.2, 162.0, 156.9, 155.9, 155.1, 151.7, 147.8, 137.0, 136.6, 134.0, 134.0, 131.5, 128.9, 126.2, 123.1, 119.1, 117.3, 115.9, 105.0, 69.6, 50.7 (2C), 49.2, 45.1, 40.9, 40.1, 33.4, 31.6, 30.0, 29.4, 29.3, 29.1, 28.9, 28.0, 25.9, 25.3, 22.8.

**HPLC-MS** $t_R$ = 1.73 min (98.2 %), ESI+ m/z 903.730; 905.531 $[M+H]^+$

**ASM136**

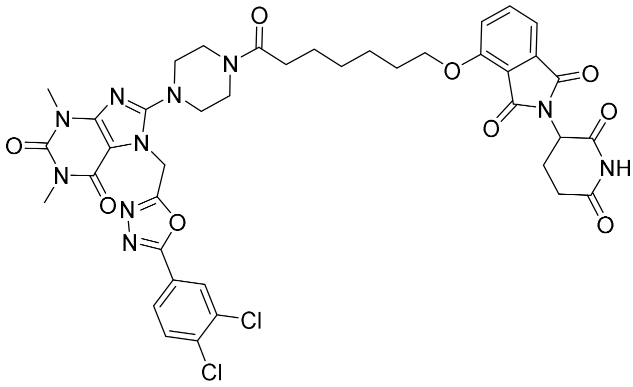

7-((5-(3,4-dichlorophenyl)-1,3,4-oxadiazol-2-yl)methyl)-8-(4-(7-((2-(2,6-dioxopiperidin-3-yl)-1,3-dioxoisoindolin-4-yl)oxy)heptanoyl)piperazin-1-yl)-1,3-dimethyl-3,7-dihydro-1*H*-purine-2,6-dione

ASM136 was prepared according to the general procedure 2 using TH5427 (15.0 mg, 0.030 mmol, 1.1 eq), 7-((2-(2,6-dioxopiperidin-3-yl)-1,3-dioxoisoindolin-4-yl)oxy)heptanoic acid (11.5 mg, 0.028 mmol, 1 eq), DIPEA (14 μL, 0.083 mmol, 3 eq) and HATU (21.1 mg, 0.055 mmol, 2 eq). After purification, a yellow solid was obtained (4.8 mg, 0.005 mmol, 19%).

**$^{1}$H NMR (400 MHz, CDCl$_3$)** δ 8.64 (s, 1H), 8.10 (d, J = 2Hz, 1H), 7.85 (dd, J1 = 2.0 Hz, J2 = 8.5 Hz, 1H), 7.67 (dd, J1 = 7.3 Hz, J2 = 8.5 Hz, 1H), 7.59 (d, J = 8.5 Hz, 1H), 7.45 (bd, J = 7.4 Hz, 1H), 7.21 (d, J = 8.5 Hz, 1H), 5.78-5.68 (m, 2H), 4.97-4.92 (m, 1H), 4.17 (t, J = 6.2 Hz, 1H), 3.72-3.62 (m, 4H), 3.62 (s, 3H), 3.38 (s, 3H), 3.29-3.25 (m, 4H), 2.91-2.70 (m, 3H), 2.46-2.30 (m, 2H), 2.17-2.10 (m, 1H), 1.91-1.84 (m, 2H), 1.71-1.66 (m, 4H), 1.61-1.54 (m, 2H), 1.50-1.45 (m, 2H).

**$^{13}$C NMR (101 MHz, CDCl$_3$)** δ 172.0, 171.1, 168.5, 167.2, 165.9, 164.2, 162.1, 156.8, 155.9, 155.1, 151.7, 147.7, 137.0, 136.7, 134.0, 133.9, 131.5, 128.9, 126.2, 123.0, 119.0, 117.3, 115.9, 105.0, 69.4, 50.8, 50.5, 49.2, 45.0, 40.9, 40.0, 32.8, 31.6, 30.0, 28.8, 28.7, 28.0, 25.6, 25.0, 22.8.

**HPLC-MS** $t_R$ = 1.63 min (98.0 %), ESI+ m/z; 875.643, 877.563 $[M+H]^+$

**HRMS** was calculated for $C_{40}H_{41}Cl_2N_{10}O_9$: 875.2435, found: 875.2448

**ASM137**

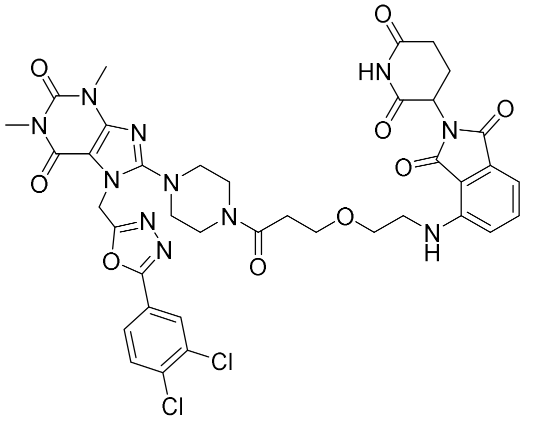

7-((5-(3,4-dichlorophenyl)-1,3,4-oxadiazol-2-yl)methyl)-8-(4-(3-(2-((2-(2,6-dioxopiperidin-3-yl)-1,3-dioxoisoindolin-4-yl)amino)ethoxy)propanoyl)piperazin-1-yl)-1,3-dimethyl-3,7-dihydro-1*H*-purine-2,6-dione

ASM137 was prepared according to the general procedure 2 using TH5427 (15.3 mg, 0.031 mmol, 1.1 eq), 3-(2-((2-(2,6-dioxopiperidin-3-yl)-1,3-dioxoisoindolin-4-yl)amino)ethoxy)propanoic acid (11.0 mg, 0.028 mmol, 1 eq), DIPEA (15 µL, 0.085 mmol, 3 eq) and HATU (21.5 mg, 0.057 mmol, 2 eq). After purification, a yellow solid was obtained (11.4 mg, 0.005 mmol, 47%).

**$^1$H NMR (400 MHz, CDCl$_3$)** δ 9.00 (s, 1H), 8.09 (d, J = 2Hz, 1H), 7.84 (dd, J1 = 2.0 Hz, J2 = 8.5 Hz, 1H), 7.58 (d, J = 8.5 Hz, 1H), 7.48 (dd, J1 = 7.1 Hz, J2 = 8.5 Hz, 1H), 7.10 (bd, J = 7.1Hz, 1H), 6.88 (d, J = 8.5 Hz, 1H), 5.79- 5.70 (m, 2H), 4.93-4.88 (m, 1H), 3.84 (t, J = 6.2 Hz, 2H), 3.78-3.60 (m, 6H), 3.53 (s, 3H), 3.45-3.43 (m, 2H), 3.38 (s, 3H), 3.28-3.22 (m, 4H), 2.88-2.85 (m, 1H), 2.80-2.71 (m, 2H), 2.70-2.59 (m, 2H), 2.16-2.11 (m, 1H). *NH-Ar not observed*

**$^{13}$C NMR (101 MHz, CDCl$_3$)** δ 171.4, 169.8, 169.5, 168.9, 167.6, 164.2, 162.1, 155.9, 155.1, 151.7, 147.7, 146.9, 137.0, 136.2, 134.0, 132.7, 131.5, 128.9, 126.2, 123.0, 116.9, 111.9, 110.5, 105.0, 69.1, 67.4, 50.6, 50.3, 49.0, 45.3, 42.2, 41.1, 40.0, 33.6, 31.5, 30.0, 28.0, 23.1.

**HPLC-MS** $t_R$ = 1.55 min (97.4 %), ESI+ m/z; 862.739, 864.600 $[M+H]^+$

**HRMS** was calculated for $C_{38}H_{38}Cl_2N_{11}O_9$: 862.2231, found: 862.2251

**Procedure for the synthesis of ASM124**

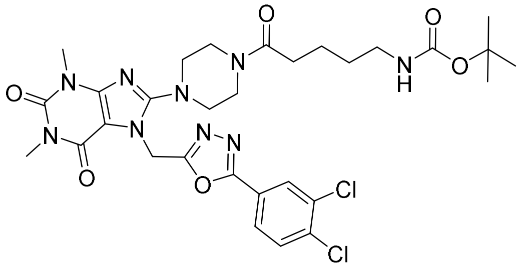

*Tert*-butyl (5-(4-(7-((5-(3,4-dichlorophenyl)-1,3,4-oxadiazol-2-yl)methyl)-1,3-dimethyl-2,6-dioxo-2,3,6,7-tetrahydro-1*H*-purin-8-yl)piperazin-1-yl)-5-oxopentyl)carbamate

*Tert*-butyl (5-(4-(7-((5-(3,4-dichlorophenyl)-1,3,4-oxadiazol-2-yl)methyl)-1,3-dimethyl-2,6-dioxo-2,3,6,7-tetrahydro-1*H*-purin-8-yl)piperazin-1-yl)-5-oxopentyl)carbamate was prepared from TH5427 TFA salt (75 mg, 0.124 mmol, 1.1 eq), 5-boc-amino-pentanoic acid (25 mg, 0.115 mmol, 1 eq), HATU (87.5 mg, 0.230 mmol, 2 eq) and DIPEA (60 µL, 0.345 mmol, 3 eq). Under nitrogen, DIPEA (2 eq) and HATU were added to a solution of acid in DMF (2.3 mL) at 0°C. After stirring for 15 min at 0°C, a solution of amine and DIPEA (1 eq) in DMF (2.3 mL) was added dropwise. The mixture was stirred overnight at room temperature. The crude was purified by flash column chromatography (Si-35, DCM/MeOH, gradient from 100/0 to 95/5) to afford the expected product as brown solid (19.1 mg, 0.028 mmol, 24%).

**$^1$H NMR (400 MHz, $CDCl_3$)** δ 8.10 (d, J = 1.9 Hz, 1H), 7.85 (dd, J = 1.9 Hz, J = 8.6 Hz, 1H), 7.59 (d, J = 8.6 Hz, 1H) 5.68 (s, 2H), 4.62 (s, 2H), 3.72 (bs, 2H), 3.58 (bs, 2H), 3.54 (s, 3H), 3.36 (s, 3H), 3.28-3.25 (m, 4H), 3.13-3.12 (m, 2H), 2.35 (t, J = 7.1 Hz, 2H), 1.71-1.63 (m, 2H), 1.56-1.49 (m, 2H), 1.42 (s, 9H).

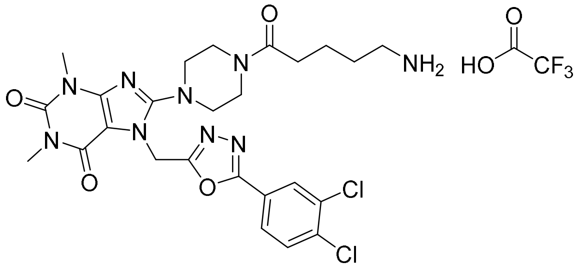

5-(4-(7-((5-(3,4-dichlorophenyl)-1,3,4-oxadiazol-2-yl)methyl)-1,3-dimethyl-2,6-dioxo-2,3,6,7-tetrahydro-1H-purin-8-yl)piperazin-1-yl)-5-oxopentan-1-aminium trifluoroacetate

*Tert*-butyl (5-(4-(7-((5-(3,4-dichlorophenyl)-1,3,4-oxadiazol-2-yl)methyl)-1,3-dimethyl-2,6-dioxo-2,3,6,7-tetrahydro-1*H*-purin-8-yl)piperazin-1-yl)-5-oxopentyl)carbamate (29 mg, 0.042 mmol, 1 eq) was dissolved in DCM (1.8 mL) then TFA (97 µL) was added at 0C. The mixture was stirred overnight at room temperature then water was added and the aqueous layer was recovered and evaporated under reduced pressure to afford the expected product as a pale white solid (29.6 mg, 0.042 mmol, quantitative).

**$^1$H NMR (400 MHz, $CDCl_3$)** δ 8.18 (d, J = 2.0 Hz, 1H), 7.95 (dd, J = 2.0 Hz, J = 8.5 Hz, 1H), 7.76 (d, J = 8.5 Hz, 1H) 5.78 (s, 2H), 3.76-3.71 (m, 4H), 3.53 (s, 3H), 3.40-3.37 (m, 2H), 3.34-3.32 (m, 2H), 3.28 (s, 3H), 3.00-2.96 (m, 2H), 2.52 (t, J = 6.6 Hz, 2H), 1.73-1.70 (m, 4H).

**ASM124**

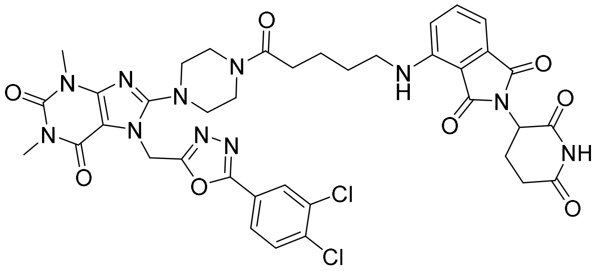

7-((5-(3,4-dichlorophenyl)-1,3,4-oxadiazol-2-yl)methyl)-8-(4-(5-((2-(2,6-dioxopiperidin-3-yl)-1,3-dioxoisoindolin-4-yl)amino)pentanoyl)piperazin-1-yl)-1,3-dimethyl-3,7-dihydro-1*H*-purine-2,6-dione

ASM124 was prepared from 5-(4-(7-((5-(3,4-dichlorophenyl)-1,3,4-oxadiazol-2-yl)methyl)-1,3-dimethyl-2,6-dioxo-2,3,6,7-tetrahydro-1H-purin-8-yl)piperazin-1-yl)-5-oxopentan-1-aminium trifluoroacetate (29.6 mg, 0.042 mmol, 1eq), 4-fluoro thalidomide (11.6 mg, 0.042 mmol, 1 eq) and triethylamine (18 µL, 0.126 mmol, 3eq). A microwave vial was charged with the reagents and DMSO (1 mL) then heated to 200°C for 15 min. The resulting mixture was then purified by prep HPLC to afford the expected product as yellow solid (1.6 mg, 0.002 mmol, 4%).

**$^1$H NMR (400 MHz, $CDCl_3$)** δ 8.34 (s, 1H), 8.10 (d, J = 2Hz, 1H), 7.86 (dd, J1 = 2.0 Hz, J2 = 8.4 Hz, 1H), 7.60 (d, J = 8.4 Hz, 1H), 7.50 (dd, J1 = 7.1 Hz, J2 = 8.4 Hz, 1H), 7.10 (d, J = 7.1 Hz, 1H), 6.89 (d, J = 8.4 Hz, 1H), 5.70 (s, 2H), 4.92-4.88 (m, 1H), 3.73-3.69 (m, 2H), 3.63-3.57 (m, 2H), 3.55 (s, 3H), 3.38 (s, 3H), 3.34-3.31 (m, 2H), 3.29-3.24 (m, 4H), 2.90-2.84 (m, 1H), 2.80-2.71 (m, 2H), 2.41 (t, J = 6.8 Hz, 2H), 2.16-2.09 (m, 1H), 1.81-1.73 (m, 4H). *NH not observed*

HPLC-MS $t_R$ = 1.59 min (97.7 %), ESI+ m/z 848.463 $[M+H]^+$

**Procedure for the synthesis of ASM102**

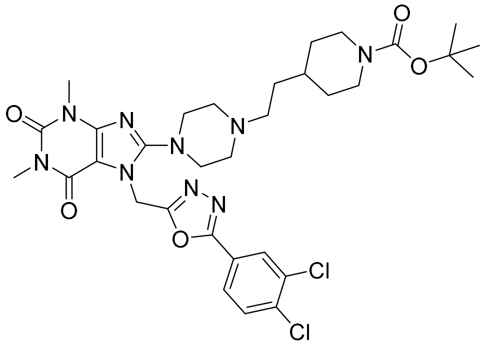

*Tert*-butyl 4-(2-(4-(7-((5-(3,4-dichlorophenyl)-1,3,4-oxadiazol-2-yl)methyl)-1,3-dimethyl-2,6-dioxo-2,3,6,7-tetrahydro-1*H*-purin-8-yl)piperazin-1-yl)ethyl)piperidine-1-carboxylate

*Tert*-butyl 4-(2-(4-(7-((5-(3,4-dichlorophenyl)-1,3,4-oxadiazol-2-yl)methyl)-1,3-dimethyl-2,6-dioxo-2,3,6,7-tetrahydro-1*H*-purin-8-yl)piperazin-1-yl)ethyl)piperidine-1-carboxylate was prepared according to the general procedure 3 using TH5427.TFA (33 mg, 0.055 mmol, 1 eq), *N*-Boc-4-piperidineacetaldehyde (14.9 mg, 0.065 mmol, 1.2 eq), sodium triacetoxyborohydride (17.3 mg, 0.082 mmol, 1.5 eq) and THF (1 mL). After purification, a light brown solid was obtained (33.1 mg, 0.047 mmol, 86%).

**$^1$H NMR (400 MHz, CDCl$_3$)** δ 8.08 (d, J = 2.0 Hz, 1H), 7.83 (dd, J = 2.0 Hz, J = 8.4 Hz, 1H), 7.57 (d, J = 8.4 Hz, 1H), 5.62 (s, 2H), 4.05 (bs, 3H), 3.53 (s, 3H), 3.34 (s, 3H), 3.33-3.32 (m, 4H), 2.68-2.61 (m, 3H) 2.58-2.55 (bs, 4H), 2.44-2.41 (m, 2H), 1.66-1.59 (m, 3H), 1.43 (s, 9H), 1.15-1.04 (m, 3H).

**$^{13}$C NMR (101 MHz, CDCl$_3$)** δ 164.0, 162.2, 156.5, 155.0, 154.9, 151.7, 148.0, 136.8, 133.9, 131.5, 128.9, 126.1, 123.2, 104.8, 79.4, 60.3, 55.9, 52.3 (2C), 50.1 (2C), 40.3 (2C), 39.4, 34.4, 32.6, 32.3 (2C), 30.0, 28.6 (3C), 27.9.

**HPLC-MS** $t_R$ = 1.86 min (94.5%), ESI+ m/z 702.824; 705.992 $[M+H]^+$

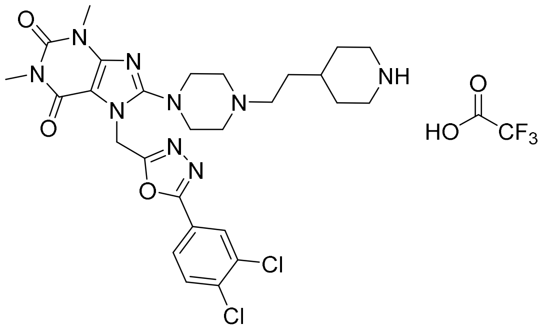

4-(2-(4-(7-((5-(3,4-dichlorophenyl)-1,3,4-oxadiazol-2-yl)methyl)-1,3-dimethyl-2,6-dioxo-2,3,6,7-tetrahydro-1*H*-purin-8-yl)piperazin-1-yl)ethyl)piperidin-1-ium trifluoroacetate

4-(2-(4-(7-((5-(3,4-dichlorophenyl)-1,3,4-oxadiazol-2-yl)methyl)-1,3-dimethyl-2,6-dioxo-2,3,6,7-tetrahydro-1*H*-purin-8-yl)piperazin-1-yl)ethyl)piperidin-1-ium trifluoroacetate was prepared from *tert*-butyl 4-((4-(7-((5-(3,4-dichlorophenyl)-1,3,4-oxadiazol-2-yl)methyl)-1,3-dimethyl-2,6-dioxo-2,3,6,7-tetrahydro-1*H*-purin-8-yl)piperazin-1-yl)ethyl)piperidine-1-carboxylate (33 mg, 0.047 mmol, 1eq), TFA (109 µL, 1.410 mmol, 30 eq) and DCM (2.8 mL) using the general procedure 4. The crude was used directly in the next step without further purification.

**$^1$H NMR (400 MHz, MeOD)** δ 8.15-8.14 (d, J = 2.0 Hz, 1H), 7.92 (dd, J = 2.0 Hz, J = 8.4 Hz, 1H), 7.73 (d, J = 8.4 Hz, 1H), 5.78 (s, 2H), 3.79-3.77 (m, 3H), 3.51 (s, 3H), 3.43-3.36 (m, 8H), 3.30 (s, 3H), 3.03-2.98 (m, 4H), 2.02-2.01 (m, 2H), 1.85-1.80 (m, 2H), 1.55-1.51 (m, 2H) *(NH.TFA not observed).*

**ASM102**

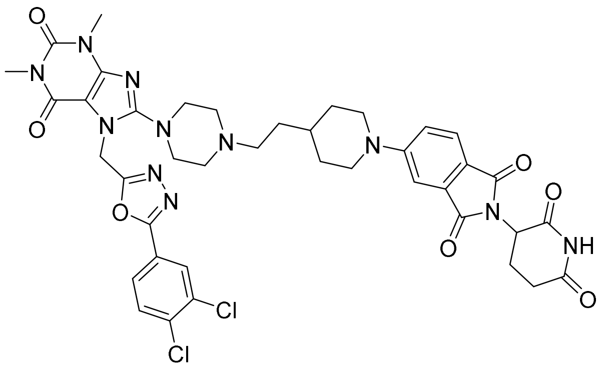

7-((5-(3,4-dichlorophenyl)-1,3,4-oxadiazol-2-yl)methyl)-8-(4-(2-(1-(2-(2,6-dioxopiperidin-3-yl)-1,3-dioxoisoindolin-5-yl)piperidin-4-yl)ethyl)piperazin-1-yl)-1,3-dimethyl-3,7-dihydro-1*H*-purine-2,6-dione

ASM102 was prepared according to the general procedure 5 using 4-(2-(4-(7-((5-(3,4-dichlorophenyl)-1,3,4-oxadiazol-2-yl)methyl)-1,3-dimethyl-2,6-dioxo-2,3,6,7-tetrahydro-1*H*-purin-8-yl)piperazin-1-yl)ethyl)piperidin-1-ium trifluoroacetate (33 mg, 0.046 mmol, 1 eq), 5-fluoro thalidomide (12.7 mg, 0.046 mmol, 1eq), TEA (19 µL, 0.138 mmol, 3 eq) and DMSO (1.2 mL). After purification, a yellow solid was obtained (1.7 mg, 0.002 mmol, 4%).

**$^{1}$H NMR (400 MHz, $CDCl_3$)** δ 8.12 (d, J = 2.0 Hz, 1H), 8.00 (bs, 1H), 7.86 (dd, J = 2.0 Hz, J = 8.5 Hz, 1H), 7.67 (d, J = 8.4 Hz, 1H) 7.61 (d, J = 8.4 Hz), 7.05- 7.02 (m, 1H), 5.65 (s, 2H), 4.96-4.91 (m, 1H), 3.96-3.93 (m, 2H), 3.55 (s, 3H), 3.37 (s, 3H), 3.00-2.92 (m, 4H), 2.88-2.70 (m, 5H), 2.16-2.12 (m, 2H), 1.85-1.82 (m, 3H), 1.62-1.58 (m, 9H).

**HRMS** was calculated for $C_{40}H_{42}Cl_2N_{11}O_7$: 858.2646, found: 858.2644

**Procedure for the synthesis of ASM105**

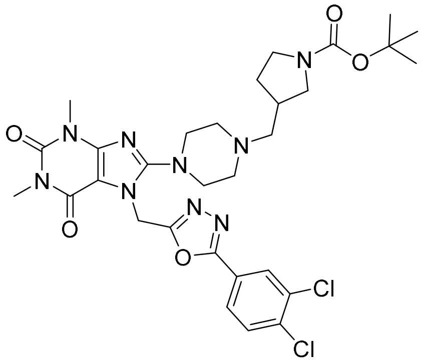

*Tert*-butyl 3-((4-(7-((5-(3,4-dichlorophenyl)-1,3,4-oxadiazol-2-yl)methyl)-1,3-dimethyl-2,6-dioxo-2,3,6,7-tetrahydro-1*H*-purin-8-yl)piperazin-1-yl)methyl)pyrrolidine-1-carboxylate

*Tert*-butyl 3-((4-(7-((5-(3,4-dichlorophenyl)-1,3,4-oxadiazol-2-yl)methyl)-1,3-dimethyl-2,6-dioxo-2,3,6,7-tetrahydro-1*H*-purin-8-yl)piperazin-1-yl)methyl)pyrrolidine-1-carboxylate was prepared according to the general procedure 3 using TH5427.TFA (50 mg, 0.083 mmol, 1 eq), *tert*-butyl 3-formylpyrrolidine-1-carboxylate (19.7 mg, 0.099 mmol, 1.2 eq), sodium triacetoxyborohydride (26.3 mg, 0.124 mmol, 1.5 eq) and THF (1.5 mL). After purification, an off-white solid was obtained (39.8 mg, 0.059 mmol, 71%).

**$^1$H NMR (400 MHz, CDCl$_3$)** δ 8.08 (d, J = 1.9 Hz, 1H), 7.84 (dd, J = 8.4 Hz, J = 2.0 Hz, 1H), 7.58 (d, J = 8.4 Hz, 1H), 5.63 (s, 2H), 3.54 (s, 3H), 3.50-3.42 (m, 4H), 3.35 (s, 3H), 3.31-3.17 (m, 4H), 3.11-2.99 (m, 2H), 2.60-2.51 (m, 3H) 2.43-2.32 (m, 3H), 1.98-1.93 (m, 1H), 1.44 (s, 9H).

**$^{13}$C NMR (101 MHz, CDCl$_3$)** δ 164.0, 162.2, 154.9, 154.7, 151.7, 148.0, 136.8, 133.9, 131.5, 128.9, 126.1, 123.2, 104.8, 79.3, 64.5, 61.4, 61.2, 52.7, 50.5, 50.2, 45.4, 45.2, 45.1, 40.3, 30.0, 28.6 (3C), 28.5, 27.9.

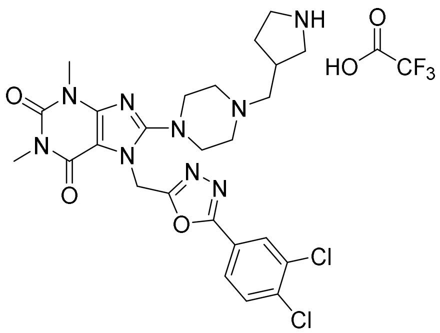

3-((4-(7-((5-(3,4-dichlorophenyl)-1,3,4-oxadiazol-2-yl)methyl)-1,3-dimethyl-2,6-dioxo-2,3,6,7-tetrahydro-1*H*-purin-8-yl)piperazin-1-yl)methyl)pyrrolidin-1-ium trifluoroacetate

3-((4-(7-((5-(3,4-dichlorophenyl)-1,3,4-oxadiazol-2-yl)methyl)-1,3-dimethyl-2,6-dioxo-2,3,6,7-tetrahydro-1*H*-purin-8-yl)piperazin-1-yl)methyl)pyrrolidin-1-ium trifluoroacetate was prepared according to the general procedure 4 using tert-butyl 3-((4-(7-((5-(3,4-dichlorophenyl)-1,3,4-oxadiazol-2-yl)methyl)-1,3-dimethyl-2,6-dioxo-2,3,6,7-tetrahydro-1*H*-purin-8-yl)piperazin-1-yl)methyl) pyrrolidine-1-carboxylate (39.8 mg, 0.059 mmol, 1eq), TFA (136 µL, 1.77 mmol, 30 eq) and DCM (2.0mL) using the general procedure 4. The crude was used directly in the next step without further purification.

**ASM105**

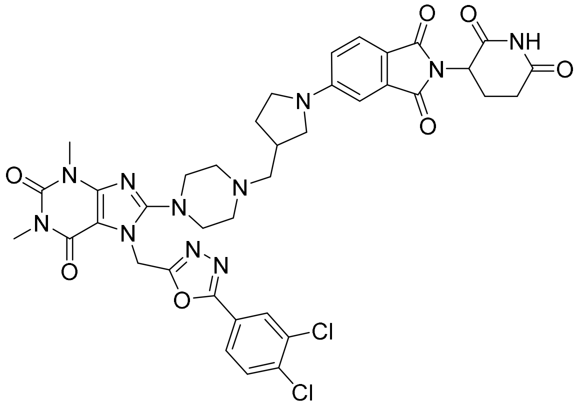

7-((5-(3,4-dichlorophenyl)-1,3,4-oxadiazol-2-yl)methyl)-8-(4-((1-(2-(2,6-dioxopiperidin-3-yl)-1,3-dioxoisoindolin-5-yl)pyrrolidin-3-yl)methyl)piperazin-1-yl)-1,3-dimethyl-3,7-dihydro-1*H*-purine-2,6-dione

ASM105 was prepared according to the general procedure 5 using 3-((4-(7-((5-(3,4-dichlorophenyl)-1,3,4-oxadiazol-2-yl)methyl)-1,3-dimethyl-2,6-dioxo-2,3,6,7-tetrahydro-1*H*-purin-8-yl)piperazin-1-yl)methyl)pyrrolidin-1-ium trifluoroacetate (40 mg, 0.058 mmol, 1 eq), 5-fluoro thalidomide (16 mg, 0.058 mmol, 1 eq), TEA (24 µL, 0.174 mmol, 3 eq) and DMSO (1.5 mL). After purification, a yellow solid was obtained (15.2 mg, 0.018 mmol, 32%).

**$^1$H NMR (400 MHz, CDCl$_3$)** δ 8.11 (d, J = 2.0 Hz, 1H), 8.06 (bs, 1H), 7.86 (dd, J = 2.0 Hz, J = 8.4 Hz, 1H), 7.65 (d, J = 8.4 Hz, 1H), 7.60 (d, J = 8.4 Hz, 1H), 6.94 (d, J = 2.0 Hz, 1H), 6.70-6.68 (m, 1H), 5.66 (s, 2H), 4.96-4.91 (m, 1H), 3.56 (s, 3H), 3.51 (bs, 1H), 3.46-3.41 (m, 2H), 3.37 (s, 3H), 3.34-3.29 (m, 2H), 3.24-3.19 (bs, 1H), 2.91-2.81 (m, 2H), 2.80-2.69 (m, 2H), 2.66-2.42 (m, 6H), 2.15-2.10 (m, 1H), 1.65-1.62 (m, 4H).

**$^{13}$C NMR (101 MHz, CDCl$_3$)** δ 171.3, 168.7, 168.3, 167.7, 164.1, 162.2, 155.0, 152.1, 151.7, 147.9, 136.9, 134.6, 133.9, 131.5, 128.9, 126.2, 125.6, 123.1, 116.8, 115.3, 106.3, 104.9, 61.4, 52.6, 50.4, 50.2 (2C), 49.2, 47.6, 40.3, 36.1, 35.9 (2C), 31.6, 30.0, 29.8, 28.0, 22.9.

**HPLC-MS** $t_R$ = 1.66 min (99.0 %), ESI+ m/z 830.598 $[M+H]^+$

**HRMS** was calculated for $C_{38}H_{38}Cl_2N_{11}O_7$: 830.2333, found: 830.2330

**Procedure for the synthesis of ASM110**

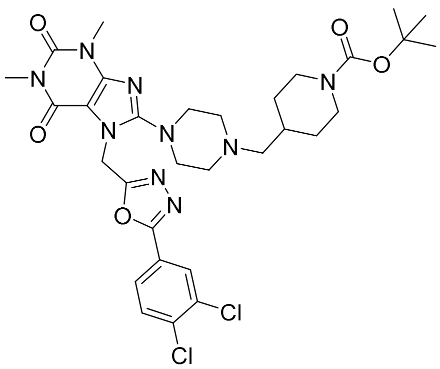

*Tert*-butyl 4-((4-(7-((5-(3,4-dichlorophenyl)-1,3,4-oxadiazol-2-yl)methyl)-1,3-dimethyl-2,6-dioxo-2,3,6,7-tetrahydro-1*H*-purin-8-yl)piperazin-1-yl)methyl)piperidine-1-carboxylate

*Tert*-butyl 4-((4-(7-((5-(3,4-dichlorophenyl)-1,3,4-oxadiazol-2-yl)methyl)-1,3-dimethyl-2,6-dioxo-2,3,6,7-tetrahydro-1*H*-purin-8-yl)piperazin-1-yl)methyl)piperidine-1-carboxylate was prepared according to the general procedure 3 using TH5427.TFA (50 mg, 0.083 mmol, 1 eq), 1-Boc-piperidine-4-carboxaldehyde (21.1 mg, 0.099 mmol, 1.2 eq), sodium triacetoxyborohydride (26.3 mg, 0.124 mmol, 1.5 eq) and THF (1.5 mL). After purification, an off-white solid was obtained (40 mg, 0.058 mmol, 70%).

**$^{1}$H NMR (400 MHz, $CDCl_3$)** δ 8.08 (d, J = 2.0 Hz, 1H), 7.83 (dd, J = 2.0 Hz, J = 8.4 Hz, 1H), 7.57 (d, J = 8.4 Hz, 1H), 5.63 (s, 2H), 4.06 (bs, 2H), 3.53 (s, 3H), 3.34 (s, 3H), 3.30-3.27 (m, 4H), 2.66 (t, J = 12.7, 2H) 2.49 (bs, 4H), 2.19 (d, J = 6.7 Hz, 2H), 1.70 (bd, J = 13.2 Hz, 2H), 1.61-1.59 (m, 1H), 1.43 (s, 9H), 1.10-1.00 (m, 2H).

**$^{13}$C NMR (101 MHz, $CDCl_3$)** δ 163.9, 162.2, 156.7, 155.0, 154.9, 151.7, 148.0, 136.8, 133.9, 131.4, 128.8, 126.1, 123.2, 104.8, 79.4, 64.4, 52.9 (2C), 50.5 (2C), 40.3 (2C), 33.6, 30.7 (2C), 29.9, 28.6 (3C), 27.9.

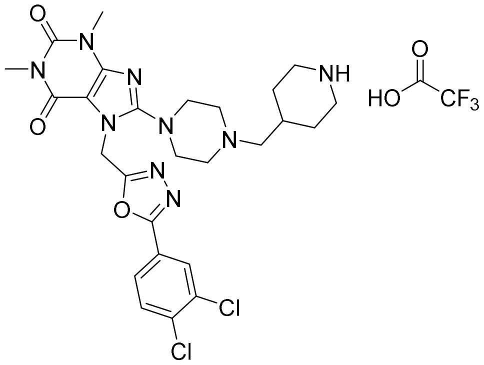

4-((4-(7-((5-(3,4-dichlorophenyl)-1,3,4-oxadiazol-2-yl)methyl)-1,3-dimethyl-2,6-dioxo-2,3,6,7-tetrahydro-1*H*-purin-8-yl)piperazin-1-yl)methyl)piperidin-1-ium trifluoroacetate

4-((4-(7-((5-(3,4-dichlorophenyl)-1,3,4-oxadiazol-2-yl)methyl)-1,3-dimethyl-2,6-dioxo-2,3,6,7-tetrahydro-1H-purin-8-yl)piperazin-1-yl)methyl)piperidin-1-ium trifluoroacetate was prepared from *tert*-butyl 4-((4-(7-((5-(3,4-dichlorophenyl)-1,3,4-oxadiazol-2-yl)methyl)-1,3-dimethyl-2,6-dioxo-2,3,6,7-tetrahydro-1H-purin-8-yl)piperazin-1-yl)methyl)piperidine-1-carboxylate (40 mg, 0.058 mmol, 1eq), TFA (134 µL, 1.743 mmol, 30 eq) and DCM (2.5 mL) using the general procedure 4. The crude was used directly in the next step without further purification.

**$^1$H NMR (400 MHz, MeOD)** δ 8.12-8.11 (d, J = 2.0 Hz, 1H), 7.90 (dd, J = 2.0 Hz, J = 8.4 Hz, 1H), 7.71 (d, J = 8.4 Hz, 1H), 5.77 (s, 2H), 3.71-3.59 (m, 7H), 3.49 (s, 3H), 3.48-3.42 (m, 3H), 3.24 (s, 3H), 3.23-3.21 (m, 2H), 3.10-3.01 (m, 2H), 2.35-2.27 (m, 1H), 2.11-2.07 (m, 2H), 1.63-1.50 (m, 2H) *NH.TFA not observed.*

**HPLC-MS** $t_R$ = 1.33 min (90.9 %), ESI+ m/z 588.714; 590.634 $[M+H]^+$ *(free NH)*

**ASM110**

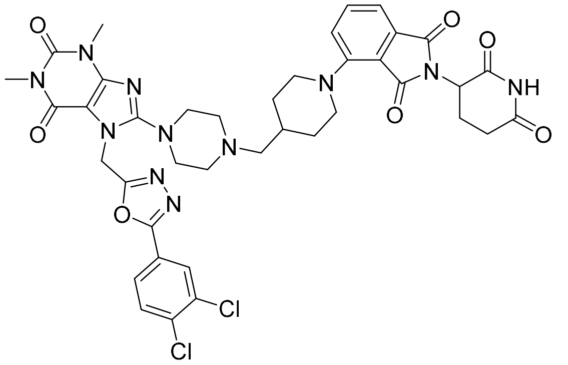

7-((5-(3,4-dichlorophenyl)-1,3,4-oxadiazol-2-yl)methyl)-8-(4-((1-(2-(2,6-dioxopiperidin-3-yl)-1,3-dioxoisoindolin-4-yl)piperidin-4-yl)methyl)piperazin-1-yl)-1,3-dimethyl-3,7-dihydro-1*H*-purine-2,6-dione

ASM110 was prepared according to the general procedure 5 using 4-((4-(7-((5-(3,4-dichlorophenyl)-1,3,4-oxadiazol-2-yl)methyl)-1,3-dimethyl-2,6-dioxo-2,3,6,7-tetrahydro-1*H*-purin-8-yl)piperazin-1-yl)methyl)piperidin-1-ium trifluoroacetate (23 mg, 0.033 mmol, 1 eq), 4-fluoro thalidomide (9 mg, 0.033 mmol, 1 eq), TEA (14 µL, 0.098 mmol, 3 eq) and DMSO (0.9 mL). After purification, a yellow solid was obtained (12.2 mg, 0.014 mmol, 44%).

**$^{1}$H NMR (400 MHz, $CDCl_3$)** δ 8.26 (bs, 1H), 8.10 (d, J = 2.0 Hz, 1H), 7.85 (dd, J = 2.0 Hz, J = 8.4 Hz, 1H), 7.59 (d, J = 8.4 Hz, 1H), 7.56-7.54 (m, 1H), 7.36 (d, J = 7.1 Hz, 1H), 7.16 (d, J = 8.4 Hz, 1H), 5.65 (s, 2H), 4.97-4.93 (m, 1H), 3.76- 3.71 (m, 2H), 3.55 (s, 3H), 3.45-3.38 (m, 2H), 3.36 (s, 3H), 2.91-2.82 (m, 4H), 2.80-2.68 (m, 3H), 2.37 (bs, 1H), 2.14-2.08 (m, 1H), 1.94-1.91 (m, 2H), 1.79-1.66 (m, 5H), 1.50-1.47 (m, 2H).

**$^{13}$C NMR (101 MHz, $CDCl_3$)** δ 171.1, 168.4, 167.5, 166.8, 164.1, 162.2, 155.0, 151.8, 150.9, 148.0, 136.9, 135.6, 134.2, 133.9, 131.5, 128.9, 126.2, 123.8, 123.2, 117.4, 115.5, 104.9, 64.4, 51.9 (2C), 51.6, 50.1, 49.2, 41.1, 40.3, 32.9, 31.6 (2C), 31.0 (2C), 30.0, 29.8, 28.0, 22.8.

**HPLC-MS** $t_R$ = 1.71 min (96.6 %), ESI+ m/z 844.535 $[M+H]^+$

**HRMS** was calculated for $C_{39}H_{40}Cl_2N_{11}O_7$: 844.2489, found: 844.2487

**ASM70**

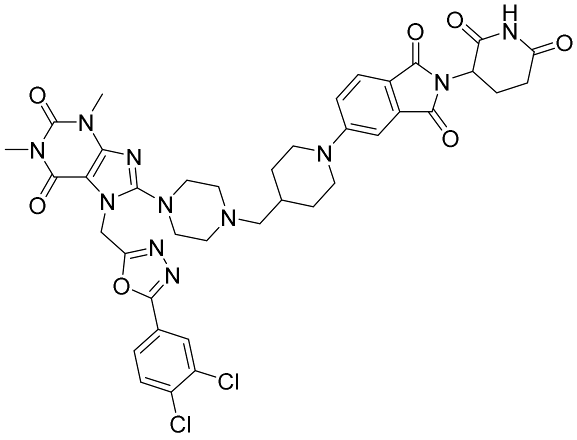

7-((5-(3,4-dichlorophenyl)-1,3,4-oxadiazol-2-yl)methyl)-8-(4-((1-(2-(2,6-dioxopiperidin-3-yl)-1,3-dioxoisoindolin-5-yl)piperidin-4-yl)methyl)piperazin-1-yl)-1,3-dimethyl-3,7-dihydro-1*H*-purine-2,6-dione

ASM70 was prepared according to the general procedure 6 using 4-((4-(7-((5-(3,4-dichlorophenyl)-1,3,4-oxadiazol-2-yl)methyl)-1,3-dimethyl-2,6-dioxo-2,3,6,7-tetrahydro-1*H*-purin-8-yl)piperazin-1-yl)methyl)piperidin-1-ium trifluoroacetate (27 mg, 0.038 mmol, 1 eq), 5-fluoro thalidomide (10.6 mg, 0.038 mmol, 1 eq), DIPEA (14 µL, 0.098 mmol, 3 eq) and DMSO (1.1 mL). After purification, a yellow solid was obtained (7.4 mg, 0.009 mmol, 23%).

**$^{1}$H NMR (400 MHz, $CDCl_3$)** δ 8.18 (s, 1H), 8.10 (d, J = 2.0 Hz, 1H), 7.85 (dd, J = 2.0 Hz, J = 8.5 Hz, 1H), 7.66 (d, J = 8.5 Hz, 1H) 7.59 (d, J = 8.5 Hz, 1H ), 7.27 (d, J = 2.3 Hz, 1H), 7.03 (dd, J = 2.3 Hz, J = 8.5 Hz, 1H), 5.65 (s, 2H), 4.95-4.91 (m, 1H), 3.96-3.93 (m, 2H), 3.56 (s, 3H), 3.37 (s, 3H), 3.32 (bs, 3H), 2.98-2.90 (m, 2H), 2.86-2.68 (m, 4H), 2.53 (bs, 4H), 2.25 (bs, 2H), 2.14-2.09 (m, 1H), 1.90-1.87 (m, 3H), 1.74, 1.32-1.26 (m, 2H).

**$^{13}$C NMR (101 MHz, $CDCl_3$)** δ 171.2, 168.5, 168.2, 167.4, 164.0, 162.2, 156.7, 155.5, 155.0, 151.8, 148.1, 136.9, 134.5, 134.0, 131.5, 128.9, 126.2, 125.6, 123.2, 118.7, 118.0, 108.8, 104.9, 64.2, 53.0 (2C), 50.5 (2C), 49.3, 48.1 (2C), 40.3, 33.4, 31.6, 30.1 (2C), 30.0, 28.0, 22.9.

**HPLC-MS** $t_R$ = 1.68 min (96.6 %), ESI+ m/z 844.795; 845.995 $[M+H]^+$

**HRMS** was calculated for $C_{39}H_{40}Cl_2N_{11}O_7$: 844.2489, found: 844.2483

**ASM71**

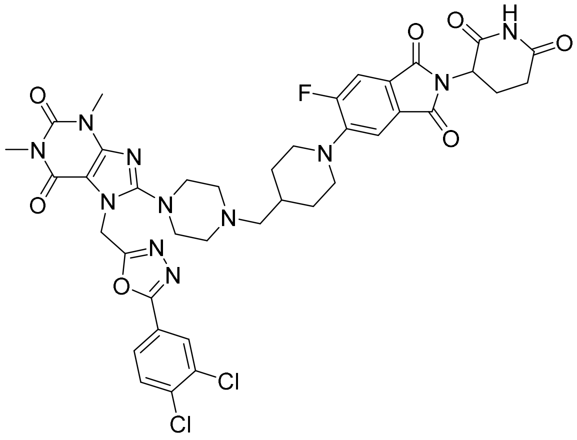

7-((5-(3,4-dichlorophenyl)-1,3,4-oxadiazol-2-yl)methyl)-8-(4-((1-(2-(2,6-dioxopiperidin-3-yl)-6-fluoro-1,3-dioxoisoindolin-5-yl)piperidin-4-yl)methyl)piperazin-1-yl)-1,3-dimethyl-3,7-dihydro-1*H*-purine-2,6-dione

ASM71 was prepared according to the general procedure 6 using 4-((4-(7-((5-(3,4-dichlorophenyl)-1,3,4-oxadiazol-2-yl)methyl)-1,3-dimethyl-2,6-dioxo-2,3,6,7-tetrahydro-1*H*-purin-8-yl)piperazin-1-yl)methyl)piperidin-1-ium trifluoroacetate (27 mg, 0.038 mmol, 1 eq), 4,5-difluoro thalidomide (11.3 mg, 0.038 mmol, 1 eq), DIPEA (14 µL, 0.098 mmol, 3 eq) and DMSO (1.1 mL). After purification, a yellow solid was obtained (6.5 mg, 0.008 mmol, 20%).

**$^{1}$H NMR (400 MHz, CDCl$_3$)** δ 8.17 (s, 1H), 8.11 (d, J = 2.0 Hz, 1H) 7.85 (dd, J = 2.0 Hz, J = 8.4 Hz, 1H), 7.59 (d, J = 8.4 Hz, 1H), 7.45 (d, J = 11.0 Hz, 1H) 7.37 (d, J = 7.2 Hz, 1H), 5.65 (s, 2H), 4.95-4.90 (m, 1H), 3.67-3.64 (m, 2H), 3.56 (s, 3H), 3.36 (s, 3H), 3.34-3.28 (m, 2H), 2.92-2.82 (m, 4H), 2.80-2.69 (m, 3H), 2.61-2.55 (m, 3H), 2.30 (bs, 1H), 2.17-2.09 (m, 1H), 1.91 (bs, 2H), 1.42 (bs, 2H), 1.28-1.25 (m, 3H).

**$^{19}$F NMR (376 MHz, CDCl$_3$)** δ -110.98

**HPLC-MS** $t_R$ = 1.77 min (97.8 %), ESI+ m/z 862.919; 864.240 $[M+H]^+$

**HRMS** was calculated for $C_{39}H_{39}Cl_2FN_{11}O_7$: 862.2395, found: 862.2414

**ASM32**

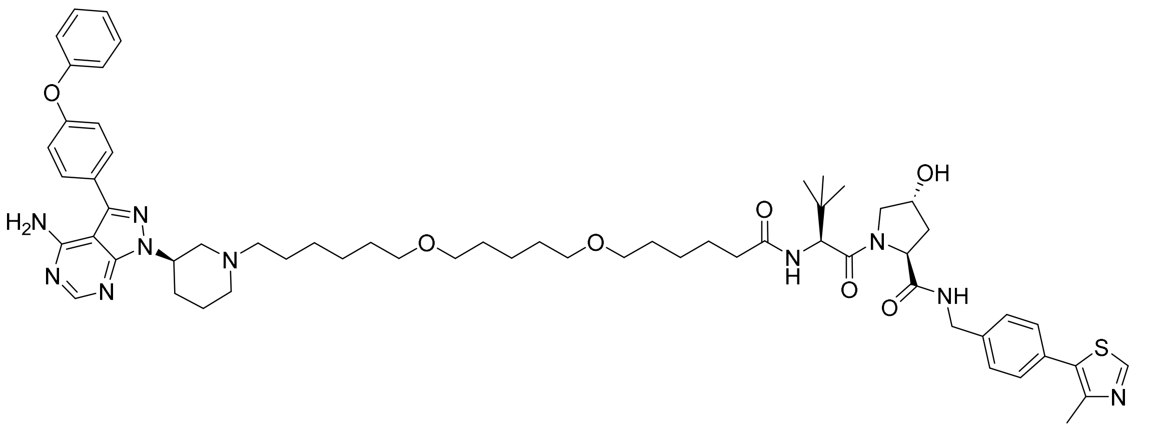

(2*S*,4*R*)-1-((*S*)-2-(6-((5-((6-((*R*)-3-(4-amino-3-(4-phenoxyphenyl)-1*H*-pyrazolo[3,4-*d*]pyrimidin-1-yl)piperidin-1-yl)hexyl)oxy)pentyl)oxy)hexanamido)-3,3-dimethylbutanoyl)-4-hydroxy-*N*-(4-(4-methylthiazol-5-yl)benzyl)pyrrolidine-2-carboxamide

(*R*)-3-(4-phenoxyphenyl)-1-(piperidin-3-yl)-1*H*-pyrazolo[3,4-*d*]pyrimidin-4-amine* (15.5 mg, 0.021 mmol, 1 eq) was dissolved in DMF (0.6 mL) then cesium carbonate (8.1 mg, 0.025 mmol, 1.2 eq) and (2*S*,4*R*)-1-((*S*)-2-(6-((5-((6-chlorohexyl)oxy)pentyl)oxy)hexanamido)-3,3-dimethylbutanoyl)-4-hydroxy-*N*-(4-(4-methylthiazol-5-yl)benzyl)pyrrolidine-2-carboxamide (8.0 mg, 0.021 mmol, 1 eq) were added. The mixture was stirred at 70°C overnight. After concentration under reduced pressure, the crude was purified by preparative HPLC to afford the expected product as white solid (3 mg, 0.003 mmol, 13%).

**$^{1}$H NMR (400 MHz, CDCl$_3$)** δ 8.66 (s, 1H), 8.35 (s, 1H), 7.64-7.62 (m, 2H), 7.40-7.29 (m, 7H), 7.18-7.12 (m, 3H), 7.08-7.05 (m, 2H), 6.51 (d, J = 8.9 Hz, 1H) 5.69 (bs, 2H), 5.04 (s, 1H), 4.72 (t, J = 8.1 Hz, 1H), 4.60-4.51 (m, 3H), 4.32 (dd, J = 5.2 Hz, 14.8 Hz, 1H), 4.13-4.10 (m, 1H), 3.59 (dd, J = 3.4 Hz, 11.3 Hz, 1H), 3.39-3.33 (m, 8H), 3.20-3.19 (m, 1H), 2.99-2.97 (m, 1H), 2.65-2.64 (m, 1H), 2.56-2.51 (m, 1H), 2.49 (s, 3H), 2.45-2.43 (m, 2H), 2.22-2.11 (m, 7H), 1.87 (bs, 2H), 1.63-1.48 (m, 12H), 1.40-1.29 (m, 8H), 0.93 (s, 9H).

**$^{13}$C NMR (101 MHz, CDCl$_3$)** δ 174.0, 172.2, 170.9, 158.5, 157.9, 156.6, 155.7, 154.2, 150.4, 148.6, 148.1, 143.7, 138.2, 131.7, 131.1, 130.2 (2C), 130.1 (2C), 129.7 (2C), 128.3 (2C), 128.1, 124.1, 119.6 (2C), 119.3 (2C), 100.0, 98.6, 71.0, 70.9, 70.8, 70.7, 70.1, 58.6, 57.7, 57.0, 53.8, 53.3, 43.4, 36.4, 36.0, 35.0, 31.0, 30.1, 29.8, 29.7, 29.6, 29.5, 27.4, 26.6 (3C), 26.2, 26.0, 25.5, 24.4, 22.9, 16.2.

**HPLC-MS** $t_R$ = 1.82 min (97.9 %), ESI+ m/z 1100.346; 1103.450 $[M+H]^+$

* Synthesized according to literature[12]

**ASM73**

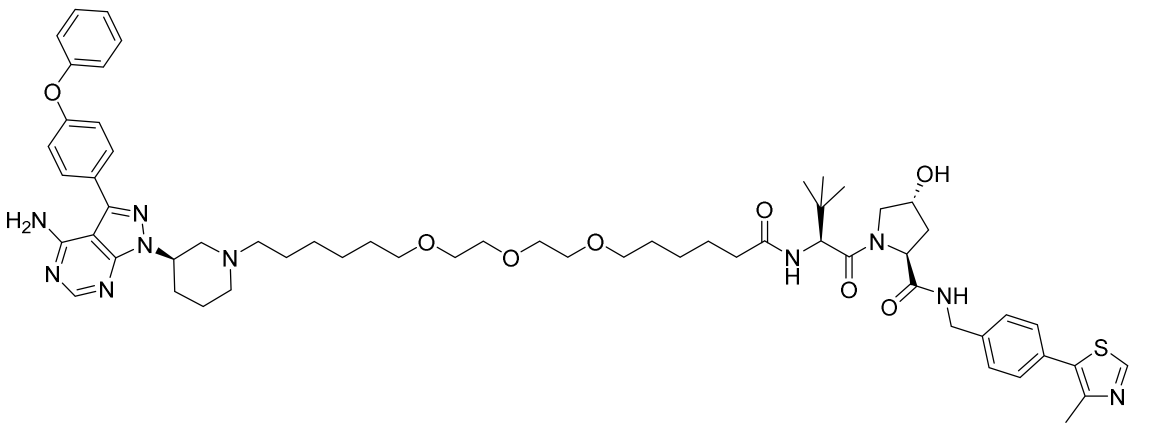

(2*S*,4*R*)-1-((*S*)-22-((*R*)-3-(4-amino-3-(4-phenoxyphenyl)-1*H*-pyrazolo[3,4-d]pyrimidin-1-yl)piperidin-1-yl)-2-(tert-butyl)-4-oxo-10,13,16-trioxa-3-azadocosanoyl)-4-hydroxy-*N*-(4-(4-methylthiazol-5-yl)benzyl)pyrrolidine-2-carboxamide

(*R*)-3-(4-phenoxyphenyl)-1-(piperidin-3-yl)-1*H*-pyrazolo[3,4-*d*]pyrimidin-4-amine (11.3 mg, 0.029 mmol, 1 eq) was dissolved in DMF (0.84 mL) then cesium carbonate (11.4 mg, 0.035 mmol, 1.2 eq) and (2*S*,4*R*)-1-((*S*)-2-(*tert*-butyl)-22-chloro-4-oxo-10,13,16-trioxa-3-azadocosanoyl)-4-hydroxy-*N*-(4-(4-methylthiazol-5-yl)benzyl) pyrrolidine-2-carboxamide (22 mg, 0.029 mmol, 1 eq) were added. The mixture was stirred at 70°C overnight. After concentration under reduced pressure, the crude was purified by preparative HPLC to afford the expected product as white solid (3.5 mg, 0.003 mmol, 11%).

Major rotamer

**$^1$H NMR (400 MHz, $CDCl_3$)** δ 8.67 (s, 1H), 7.64-7.61 (m, 1H), 7.40-7.30 (m, 6H), 7.18-7.11 (m, 2H), 6.23 (d, J = 8.8 Hz, 1H), 4.70 (t, J = 8.1 Hz, 1H), 4.59-4.50 (m, 3H), 4.35-4.29 (m, 1H), 4.10-4.07 (m, 1H), 3.63-3.50 (m, 13H), 3.46-3.39 (m, 5H), 2.83-2.66 (m, 5H), 2.50 (s, 3H), 2.49-2.45 (m, 2H), 2.21-2.10 (m, 4H), 1.80-1.72 (m, 2H), 1.64-1.51 (m, 8H), 1.48-1.29 (m, 8H), 0.92 (s, 9H).

**ASM74**

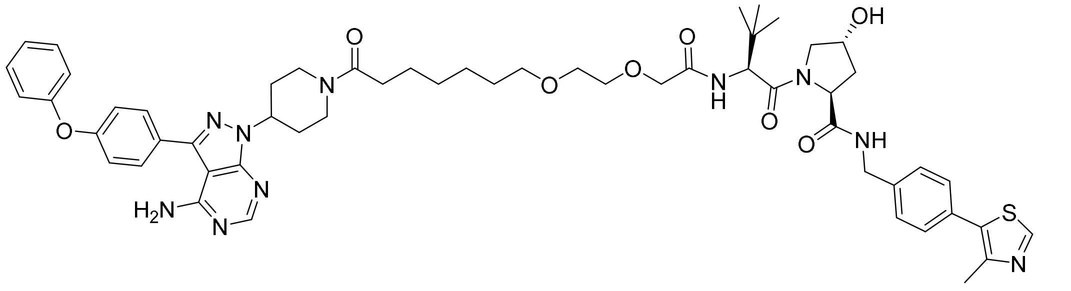

(2*S*,4*R*)-1-((*S*)-2-(2-(2-((7-(4-(4-amino-3-(4-phenoxyphenyl)-1*H*-pyrazolo[3,4-*d*]pyrimidin-1-yl)piperidin-1-yl)-7-oxoheptyl)oxy)ethoxy)acetamido)-3,3-dimethylbutanoyl)-4-hydroxy-*N*-(4-(4-methylthiazol-5-yl)benzyl)pyrrolidine-2-carboxamide

ASM74 was prepared according to the general procedure 2 using 3-(4-phenoxyphenyl)-1-(piperidin-3-yl)-1*H*-pyrazolo[3,4-*d*]pyrimidin-4-amine (7.7 mg, 0.020 mmol, 1.1 eq), 7-(2-(2-(((*S*)-1-((2*S*,4*R*)-4-hydroxy-2-((4-(4-methylthiazol-5-yl)benzyl)carbamoyl)pyrrolidin-1-yl)-3,3-dimethyl-1-oxobutan-2-yl)amino)-2-oxoethoxy)ethoxy)heptanoic acid (12 mg, 0.018 mmol, 1 eq), DIPEA (6 μL, 0.036 mmol, 2 eq) and HATU (13.8 mg, 0.036 mmol, 2 eq). After purification, a white solid was obtained (7.2 mg, 0.007 mmol, 39%).

**$^{1}$H NMR (400 MHz, $CDCl_3$)** δ 8.66 (s, 1H), 8.35 (s, 1H), 7.64-7.60 (m, 2H), 7.57-7.52 (m, 1H), 7.40-7.31 (m, 7H), 7.19-7.12 (m, 3H), 7.08-7.06 (m, 2H), 5.70 (s, 2H), 5.02-4.96, 4.79-4.69 (m, 2H), 4.59-4.50 (m, 3H), 4.34 (dd, J = 15.0 Hz, J = 5.5 Hz, 1H), 4.10-4.03 (m, 2H), 4.00-3.99 (m, 2H), 3.67-3.57 (m, 4H), 3.46 (t, J = 6.5 Hz, 2H), 3.25 (t, J = 13.1 Hz, 1H), 2.82-2.78 (m, 1H), 2.50 (s, 3H), 2.47-2.45 (m, 1H), 2.38-2.31 (m, 2H), 2.27-2.20 (m, 2H), 2.16-2.03 (m, 5H), 1.65-1.55 (m, 4H), 1.36-1.34 (m, 4H), 0.96 (s, 9H).

**HPLC-MS** $t_R$ = 1.59 min (98.7 %), ESI+ m/z 1030.154 $[M+H]^+$

**ASM75**

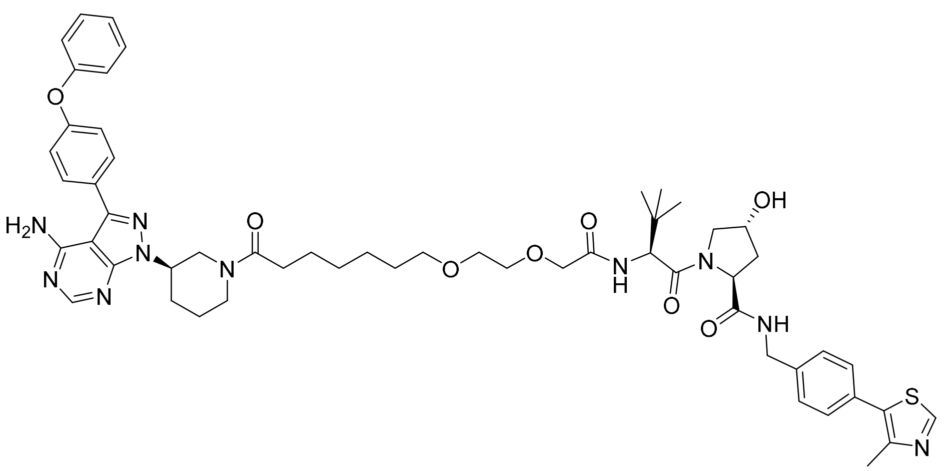

(2*S*,4*R*)-1-((*S*)-2-(2-(2-((7-((*R*)-3-(4-amino-3-(4-phenoxyphenyl)-1*H*-pyrazolo[3,4-d]pyrimidin-1-yl)piperidin-1-yl)-7-oxoheptyl)oxy)ethoxy)acetamido)-3,3-dimethylbutanoyl)-4-hydroxy-*N*-(4-(4-methylthiazol-5-yl)benzyl)pyrrolidine-2-carboxamide

ASM75 was prepared according to the general procedure 2 using (*R*)-3-(4-phenoxyphenyl)-1-(piperidin-3-yl)-1*H*-pyrazolo[3,4-*d*]pyrimidin-4-amine (7.7 mg, 0.020 mmol, 1.1 eq), 7-(2-(2-(((*S*)-1-((2*S*,4*R*)-4-hydroxy-2-((4-(4-methylthiazol-5-yl)benzyl)carbamoyl)pyrrolidin-1-yl)-3,3-dimethyl-1-oxobutan-2-yl)amino)-2-oxoethoxy)ethoxy)heptanoic acid (12 mg, 0.018 mmol, 1 eq), DIPEA (6 µL, 0.036 mmol, 2 eq) and HATU (13.8 mg, 0.036 mmol, 2 eq). After purification, a white solid was obtained (8.7 mg, 0.008 mmol, 46%).

Major rotamer

**$^1$H NMR (400 MHz, $CDCl_3$)** δ 8.65 (s, 1H), 8.34 (s, 1H), 7.64-7.55 (m, 3H), 7.41-7.32 (m, 7H), 7.19-7.13 (m, 3H), 7.09-7.06 (m, 2H), 5.75 (s, 2H), 4.85-4.69 (m, 2H), 4.59-4.49 (m, 3H), 4.36-4.31 (m, 1H), 4.09-4.05 (m, 1H), 4.02-3.97 (m, 2H), 3.67-3.53 (m, 6H), 3.48-3.40 (m, 2H), 3.30 (t, J = 12.6 Hz, 1H), 2.49 (s, 3H), 2.48-2.44 (m, 1H), 2.39-2.29 (m, 3H), 2.25-2.19 (m, 3H), 2.16-2.11 (m, 2H), 2.00-1.95 (m, 1H), 1.68-1.52 (m, 5H), 1.35-1.28 (m, 4H), 0.95 (s, 9H).

**HPLC-MS** $t_R$ = 1.66 min (98.8 %), ESI+ m/z 1030.090 $[M+H]^+$

**Procedures for the synthesis of dNUDT5***

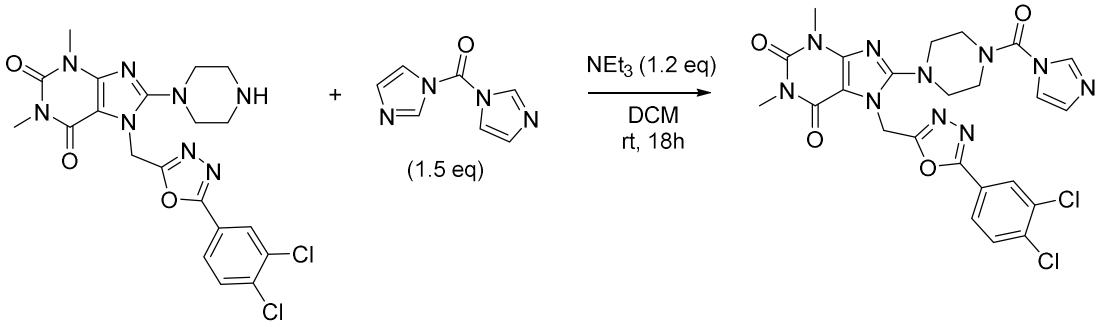

8-(4-(1*H*-imidazole-1-carbonyl)piperazin-1-yl)-7-((5-(3,4-dichlorophenyl)-1,3,4-oxadiazol-2-yl)methyl)-1,3-dimethyl-3,7-dihydro-1*H*-purine-2,6-dione

Under nitrogen, 1,1′-carbonyldiimidazole (88.6 mg, 0.546 mmol, 1.5 eq) was added at -5°C to a mixture of TH5427 (179 mg, 0.364 mmol, 1 eq) and triethylamine (61 µL, 0.437 mmol, 1.2 eq) in DCM (14.3 mL). The mixture was then stirred overnight at room temperature. After concentration under reduced pressure, methanol was added and the resulting solid was collected by filtration to afford the expected product as white solid (213 mg, 0.364 mmol, quantitative).

**$^{1}$H NMR (400 MHz, $CDCl_3$)** δ 8.12 (d, 1H, J = 2.0 Hz), 8.03 (bs, 1H), 7.87 (dd, 1H, J = 2.0 Hz J = 8.4 Hz), 7.61 (d, 1H, J = 8.4 Hz), 7.24 (bs, 1H), 7.16 (bs, 1H), 5.77 (s, 2H), 3.79-3.77 (m, 4H), 3.56 (s, 3H), 3.41-3.40 (m, 4H), 3.39 (s, 3H).

**$^{13}$C NMR (101 MHz, $CDCl_3$)** δ 164.3, 161.9, 155.2, 155.1, 151.6, 150.9, 147.6, 137.1, 137.0, 134.0, 131.6, 129.8, 128.9, 126.2, 123.0, 118.1, 105.1, 50.5, 45.9 (2C), 40.0 (2C), 30.0, 28.0.

**HPLC-MS** $t_R$ = 1.46 min (97.2 %), ESI+ m/z 585.533; 587.153 $[M+H]^+$

*Adapted from[13]

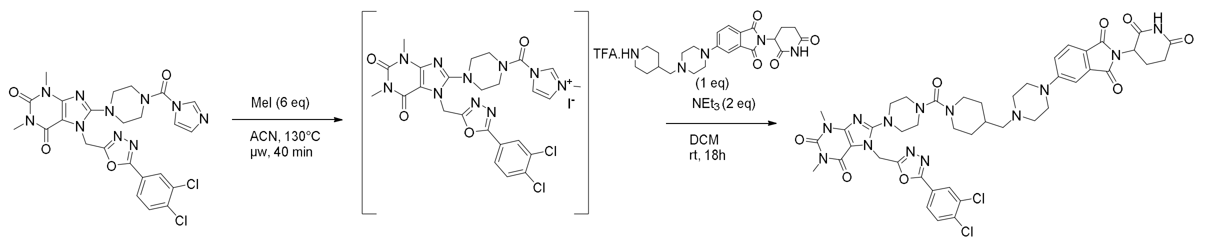

7-((5-(3,4-dichlorophenyl)-1,3,4-oxadiazol-2-yl)methyl)-8-(4-(4-((4-(2-(2,6-dioxopiperidin-3-yl)-1,3-dioxoisoindolin-5-yl)piperazin-1-yl)methyl)piperidine-1-carbonyl)piperazin-1-yl)-1,3-dimethyl-3,7-dihydro-1*H*-purine-2,6-dione

Under $N_2$, methyl iodide (16 µL, 0.256 mmol, 6 eq) was added to a mixture of 8-(4-(1H-imidazole-1-carbonyl)piperazin-1-yl)-7-((5-(3,4-dichlorophenyl)-1,3,4-oxadiazol-2-yl)methyl)-1,3-dimethyl-3,7-dihydro-1H-purine-2,6-dione (25 mg, 0.043 mmol, 1 eq) in acetonitrile (1.7 mL). The mixture was stirred for 40 min at 130°C under microwave heating (the reaction was checked by LCMS – 100% conversion).* After concentration under reduced pressure, the crude was dissolved in DCM (2 mL). Under $N_2$, triethylamine (12 µL, 0.086 mmol, 2 eq) and 2-(2,6-dioxopiperidin-3-yl)-5-(4-((1-(2,2,2-trifluoroacetyl)-1λ4-piperidin-4-yl)methyl)piperazin-1-yl)isoindoline-1,3-dione, trifluoroacetic salt (23.8 mg, 0.043 mmol, 1 eq) were added. The mixture was stirred overnight at room temperature. After concentration under reduced pressure, the crude was purified by HPLC to afford the expected product as yellow solid (6.9 mg, 0.007 mmol, 17% over 2 steps).

**$^1$H NMR (400 MHz, $CDCl_3$)** δ 8.11 (d, J = 2.0 Hz, 1H), 8.06 (s, 1H), 7.86 (dd, J = 2.0 Hz, J = 8.4 Hz, 1H), 7.67 (d, J = 8.4 Hz, 1H), 7.60 (d, J = 8.4 Hz, 1H), 7.28 (d, J = 2.2 Hz, 1H), 7.05 (dd, J = 2.0 Hz, J = 8.5 Hz, 1H), 5.66 (s, 2H), 4.96-4.91 (m, 1H), 3.75-3.72 (m, 2H), 3.55 (s, 3H), 3.43-3.40 (m, 4H), 3.37 (s, 3H), 3.35-3.33 (m, 3H), 3.30-3.28 (m, 4H), 2.91-2.81 (m, 2H), 2.80-2.69 (m, 3H), 2.60-2.55 (m, 3H), 2.27-2.26 (m, 2H), 2.17-2.10 (m, 1H), 1.81-1.78 (m, 2H), 1.74-1.69 (m, 2H), 1.21-1.12 (m, 2H)

**$^{13}$C NMR (101 MHz, $CDCl_3$)** δ 171.0, 168.3, 168.0, 167.3, 164.1, 164.0, 162.1, 156.3, 155.6, 155.0, 151.7, 147.9, 136.9, 134.4, 134.0, 131.5, 128.9, 126.2, 125.5, 123.1, 119.7, 118.0, 108.8, 104.9, 64.4, 53.2 (2C), 50.5 (2C), 49.3, 47.6 (2C), 47.1 (2C), 46.5 (2C), 40.2, 33.9, 31.6, 30.8 (2C), 30.0, 28.0, 22.9.

**HPLC-MS** $t_R$ = 1.56 min (96.9 %), ESI+ m/z 957.9246; 959.9651 $[M+H]^+$

**HRMS** was calculated for $C_{44}H_{48}Cl_2N_{13}O_8$: 956.3126, found: 956.3120

*Adapted from[14]

**Procedures for the synthesis of dNUDT5nc**

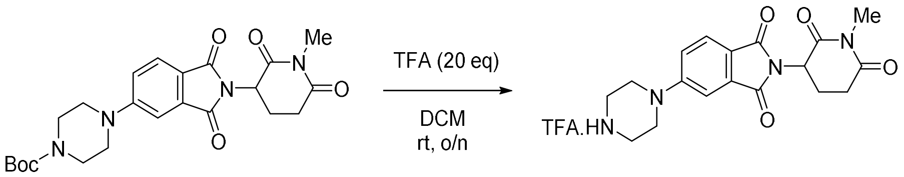

2-(1-methyl-2,6-dioxopiperidin-3-yl)-5-(piperazin-1-yl)isoindoline-1,3-dione (trifluoroacetic salt)

Trifluoroacetic acid (429 µL, 5.564 mmol, 20 eq) was added dropwise to a solution of tert-butyl 4-(2-(1-methyl-2,6-dioxopiperidin-3-yl)-1,3-dioxoisoindolin-5-yl)piperazine-1-carboxylate (127 mg, 0.278 mmol, 1 eq) in DCM (4 mL) at 0°C. The mixture was stirred for 3 h at room temperature. Water was added and the aqueous layer was recovered and concentrated under reduced pressure to afford the expected product as a yellow oil (99.5 mg, 0.212 mmol, 76%).

**$^1$H NMR (400 MHz, MeOD)** δ 7.74 (d, J = 8.5 Hz, 1H), 7.44 (d, J = 2.3 Hz, 1H), 7.32 (dd, J = 2.3 Hz, J = 8.5 Hz, 1H), 5.11 (dd, J = 5.5 Hz, J = 13.0 Hz, 1H), 3.71-3.69 (m, 4H), 3.41-3.38 (m, 4H), 3.31 (s, 3H), 2.91-2.87 (m, 2H), 2.75-2.68 (m, 1H), 2.13-2.06 (m, 1H).

**$^{13}$C NMR (101 MHz, MeOD)** δ 173.6, 171.4, 169.0, 168.7, 156.2, 135.5, 126.1, 120.6, 110.5, 51.2, 49.0, 46.0 (2C), 44.3 (2C), 32.5, 27.3, 22.9 (TFA not detected).

**HPLC-MS** $t_R$ = 1.08 min (100%), ESI+ m/z 357.4402 [M+H]$^+$

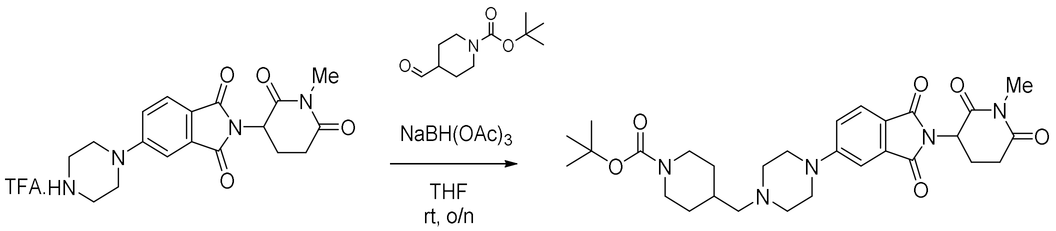

*Tert*-butyl 4-((4-(2-(1-methyl-2,6-dioxopiperidin-3-yl)-1,3-dioxoisoindolin-5-yl)piperazin-1-yl)methyl)piperidine-1-carboxylate

Sodium triacetoxyborohydride (67 mg, 0.316 mmol, 1.5 eq) was added to a mixture of 2-(1-methyl-2,6-dioxopiperidin-3-yl)-5-(piperazin-1-yl)isoindoline-1,3-dione, trifluoroacetic salt (99 mg, 0.210 mmol, 1 eq) and tert-butyl 4-formylpiperidine-1-carboxylate (54 mg, 0.253 mmol, 1.2 eq) in THF (4.6 mL). The mixture was stirred overnight at room temperature. After concentration under reduced pressure, the crude was purified by column chromatography (Si-35, DCM/MeOH gradient from 100% DCM to 95%) to afford the expected product as yellow solid (112.3 mg, 0.203 mmol, 96%).

**$^{1}$H NMR (400 MHz, $CDCl_3$)** δ 7.65 (d, J = 8.6 Hz, 1H), 7.24 (d, J = 2.3 Hz, 1H), 7.02 (dd, J = 2.3 Hz, J = 8.6 Hz, 1H), 4.94-4.90 (m, 1H), 4.07 (bs, 2H), 3.41 (bs, 4H), 3.16 (s, 3H), 2.98-2.89 (m, 1H), 2.81-2.65 (m, 4H), 2.58-2.56 (m, 4H), 2.24 (bs, 2H), 2.08-2.04 (m, 1H), 1.75-1.67 (m, 3H), 1.43 (s, 9H), 1.13-1.03 (m, 2H).

**$^{13}$C NMR (101 MHz, $CDCl_3$)** δ 171.3, 169.1, 168.1, 167.4, 155.4, 154.9, 134.3, 134.3, 125.3, 117.9, 108.6, 79.3, 64.3, 53.0 (4C), 49.9, 47.4 (2C), 33.5, 32.0 (2C), 30.7, 28.5, 27.2, 22.1 (C=O Boc not detected).

**HPLC-MS** $t_R$ = 1.77 min (97.3 %), ESI+ m/z 554.8069 $[M+H]^+$

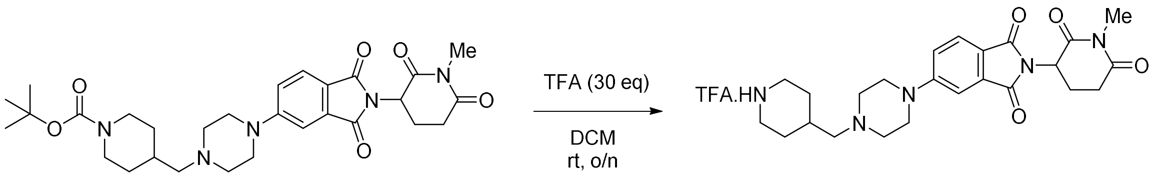

2-(1-methyl-2,6-dioxopiperidin-3-yl)-5-(4-(piperidin-4-ylmethyl)piperazin-1-yl)isoindoline-1,3-dione (trifluoroacetic salt)

Trifluoroacetic acid (468 µL, 6.069 mmol, 30 eq) was added dropwise to a solution of tert-butyl 4-((4-(2-(1-methyl-2,6-dioxopiperidin-3-yl)-1,3-dioxoisoindolin-5-yl)piperazin-1-yl)methyl)piperidine-1-carboxylate (835 mg, 1.547 mmol, 1 eq) in DCM (10 mL) at 0°C. The mixture was stirred for 3h at room temperature. Water was added and the aqueous layer was recovered and concentrated under reduced pressure to afford the expected product as a yellow solid (863 mg, 1.54 mmol, quantitative).

**$^{1}$H NMR (400 MHz, MeOD)** δ 7.77 (d, J = 8.5 Hz, 1H), 7.48 (d, J = 2.3 Hz, 1H), 7.35 (dd, J = 2.3 Hz, J = 8.5 Hz, 1H), 5.15-5.10 (m, 1H), 3.78 (bs, 3H), 3.48-3.45 (m, 6H), 3-18-3.16 (m, 2H), 3.14 (s, 3H), 3.09-3.02 (m, 2H), 2.91-2.87 (m, 2H), 2.75-2.64 (m, 1H), 2.32-2.27 (m, 1H), 2.12-2.08 (m, 3H), 1.61-1.50 (m, 2H) (TFA not detected).

**$^{13}$C NMR (101 MHz, MeOD)** δ 173.6, 171.4, 169.0, 168.7, 155.9, 135.6, 126.1, 122.9, 120.5, 110.4, 62.3, 53.1 (2C), 51.2, 46.1 (2C), 44.4 (2C), 32.5, 30.3, 27.8 (2C), 27.3, 23.0 (TFA not detected).

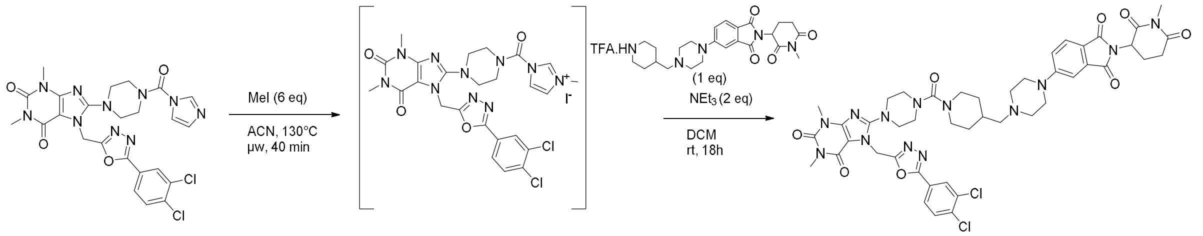

7-((5-(3,4-dichlorophenyl)-1,3,4-oxadiazol-2-yl)methyl)-1,3-dimethyl-8-(4-(4-((4-(2-(1-methyl-2,6-dioxopiperidin-3-yl)-1,3-dioxoisoindolin-5-yl)piperazin-1-yl)methyl)piperidine-1-carbonyl)piperazin-1-yl)-3,7-dihydro-1*H*-purine-2,6-dione

Under $N_2$, methyl iodide (16 µL, 0.256 mmol, 6 eq) was added to a mixture of 8-(4-(1H-imidazole-1-carbonyl)piperazin-1-yl)-7-((5-(3,4-dichlorophenyl)-1,3,4-oxadiazol-2-yl)methyl)-1,3-dimethyl-3,7-dihydro-1*H*-purine-2,6-dione (25 mg, 0.043 mmol, 1 eq) in acetonitrile (1.7 mL). The mixture was stirred for 40 min at 130°C under microwave heating (the reaction was checked by LCMS – 95% conversion). After concentration under reduced pressure, the crude was dissolved in DCM (1.8 mL). Under $N_2$, were added triethylamine (12 µL, 0.086 mmol, 2 eq) and 2-(1-methyl-2,6-dioxopiperidin-3-yl)-5-(4-(piperidin-4-ylmethyl)piperazin-1-yl)isoindoline-1,3-dione, trifluoroacetic salt (24.4 mg, 0.043 mmol, 1 eq). The mixture was stirred overnight at room temperature. After concentration under reduced pressure, the crude was purified by HPLC to afford the expected product as yellow solid (5.9 mg, 0.006 mmol, 14% over 2 steps).

**$^1$H NMR (400 MHz, $CDCl_3$)** δ 8.12 (d, J = 2.0 Hz, 1H), 7.87 (dd, J = 2.0 Hz, J = 8.4 Hz, 1H), 7.70 (d, J = 8.4 Hz, 1H), 7.60 (d, J = 8.4 Hz, 1H), 7.29 (d, J = 2.0 Hz, 1H), 7.06 (dd, J = 2.0 Hz, J = 8.4 Hz, 1H), 5.67 (s, 2H), 4.97-4.92 (m, 1H), 3.76-3.73 (m, 2H), 3.56 (s, 3H), 3.44-3.41 (m, 4H), 3.38 (s, 3H), 3.36-3.34 (m, 3H), 3.31-3.29 (m, 4H), 3.32 (s, 3H), 3.00-2.94 (m, 1H), 2.85-2.74 (m, 4H), 2.60-2.57 (m, 4H), 2.27 (d, J = 6.7 Hz, 2H), 2.13-2.08 (m, 1H), 1.83-1.79 (m, 2H), 1.22-1.11 (m, 4H).

**$^{13}$C NMR (101 MHz, $CDCl_3$)** δ 171.4, 169.1, 168.2, 167.5, 164.1, 164.0, 162.1, 156.3, 155.6, 155.0, 151.7, 147.9, 137.0, 134.4, 134.0, 131.5, 128.9, 126.2, 125.5, 123.1, 119.8, 118.0, 108.7, 105.0, 77.2, 64.4, 53.2 (2C), 50.5 (2C), 50.1, 47.6 (2C), 47.1 (2C), 46.5 (2C), 40.2, 33.9, 32.1, 30.8 (2C), 30.0, 28.0, 27.4, 22.2.

**HPLC-MS** $t_R$ = 1.71 min (98.7 %), ESI+ m/z 970.7936; 971.9340 $[M+H]^+$
